# Potent, Novel SARS-CoV-2 PLpro Inhibitors Block Viral Replication in Monkey and Human Cell Cultures

**DOI:** 10.1101/2021.02.13.431008

**Authors:** Zhengnan Shen, Kiira Ratia, Laura Cooper, Deyu Kong, Hyun Lee, Youngjin Kwon, Yangfeng Li, Saad Alqarni, Fei Huang, Oleksii Dubrovskyi, Lijun Rong, Gregory Thatcher, Rui Xiong

**Affiliations:** Department of Pharmaceutical Sciences, College of Pharmacy, University of Illinois at Chicago (UIC), Chicago, IL, USA; UICentre (Drug Discovery @ UIC), UIC; Department of Microbiology, College of Medicine, UIC; Research Resources Center, University of Illinois at Chicago (UIC), Chicago, IL, USA; Department of Pharmacology & Toxicology, College of Pharmacy, University of Arizona, Tucson, AZ, USA

**Author notes:** Corresponding authors: Rui Xiong, and Gregory Thatcher. These authors contributed equally to this work (Zhengnan Shen, Kiira Ratia, Laura Cooper).

## Abstract

Antiviral agents blocking SARS-CoV-2 viral replication are desperately needed to complement vaccination to end the COVID-19 pandemic. Viral replication and assembly are entirely dependent on two viral cysteine proteases: 3C-like protease (3CLpro) and the papain-like protease (PLpro). PLpro also has deubiquitinase (DUB) activity, removing ubiquitin (Ub) and Ub-like modifications from host proteins, disrupting the host immune response. 3CLpro is inhibited by many known cysteine protease inhibitors, whereas PLpro is a relatively unusual cysteine protease, being resistant to blockade by such inhibitors. A high-throughput screen of biased and unbiased libraries gave a low hit rate, identifying only CPI-169 and the positive control, GRL0617, as inhibitors with good potency (IC_50_ < 10 µM). Analogues of both inhibitors were designed to develop structure-activity relationships; however, without a co-crystal structure of the CPI-169 series, we focused on GRL0617 as a starting point for structure-based drug design, obtaining several co-crystal structures to guide optimization. A series of novel 2-phenylthiophene-based non-covalent SARS-CoV-2 PLpro inhibitors were obtained, culminating in low nanomolar potency. The high potency and slow inhibitor off-rate were rationalized by newly identified ligand interactions with a “BL2 groove” that is distal from the active site cysteine. Trapping of the conformationally flexible BL2 loop by these inhibitors blocks binding of viral and host protein substrates; however, until now it has not been demonstrated that this mechanism can induce potent and efficacious antiviral activity. In this study, we report that novel PLpro inhibitors have excellent antiviral efficacy and potency against infectious SARS-CoV-2 replication in cell cultures. Together, our data provide structural insights into the design of potent PLpro inhibitors and the first validation that non-covalent inhibitors of SARS-CoV-2 PLpro can block infection of human cells with low micromolar potency.

---

The COVID-19 pandemic, caused by the novel severe acute respiratory syndrome coronavirus 2 (SARS-CoV-2), has caused profound socioeconomic challenges for humankind.^1 2,3^ By the end of January 2021, the number of cases exceeded 100 million and over 2.2 million people have died from COVID-19, with worldwide daily deaths still increasing and deaths in the USA amplified by health disparities.^4^ Currently approved antiviral agents have not effectively addressed the current pandemic and we are learning belatedly that it is essential to proactively create new antiviral agents for future outbreaks of this and other zoonotic viruses. The expedited approval, distribution and administration of the first vaccines is one important step in ending the pandemic. However, the resurgence of COVID-19 in a population with high seroprevalence in Manaus, Brazil,^5^ questions the long-term effects of immunoprotection. With the evolution and spread of new variants there exists an urgent need to develop small molecule antiviral agents to treat patients who do not respond to or cannot tolerate vaccines and to address future outbreaks.

The early sequencing of the SARS-CoV-2 genome has allowed comparisons with other coronaviruses including the Middle East Respiratory Syndrome CoV (MERS-CoV) and the earlier SARS-CoV, which like SARS-CoV-2 uses the angiotensin-converting enzyme 2 (ACE2) receptor to enter host cells **(Figure S1**).^6,7,8^ SARS-CoV-2 shares 86% overall amino acid sequence identity with SARS-CoV and ∼ 50% with MERS-CoV.^2^ The high homology of SARS-CoV-2 to other coronaviruses has allowed the rapid understanding of viral biology, from particle attachment, entry, replication and primary translation (polyprotein processing), assembly, maturation, to release and shedding.^9,10^ The SARS-CoV-2 spike protein recognizes and attaches to ACE2 and utilizes the cell surface serine protease TMPRSS2 to promote viral entry. ^6,7 8,11^ Following entry, viral RNA is translated by the host ribosome to yield two large overlapping open reading frames (ORFs), ORF1a and ORF1b. Two viral cysteine proteases, the coronavirus main protease (3CLpro; nsp5) and the papain-like protease (PLpro; nsp3), proteolytically process these two viral polyproteins to yield individual non-structural proteins (nsps) that then assemble in complexes with host membrane components.^12^ The RNA-dependent RNA polymerase encoded by nsp12, a proteolytic product of 3CLpro, is a molecular target of the FDA-approved COVID-19 treatment, remdesivir.^13^ PLpro recognizes the P4–P1 sequence LxGG and cleaves at three sites to release nsps 1-3. Nsp3 (1922aa, 215 kDa) incorporates PLpro itself (residues 1602–1855) and is the largest component of the replication and transcription complex.^14,15^ The catalytic activity of 3CLpro and PLpro is essential for viral replication, making inhibition of these enzymes a compelling strategy for antiviral therapy.

PLpro supports viral replication beyond the role of viral polyprotein processing, since host proteins are substrates for PLpro. Specifically, PLpro disrupts the host innate immune response by cleaving the isopeptide bond that ligates ubiquitin (Ub) and ubiquitin-like proteins (UbL) such as interferon-stimulated gene product 15 (ISG15) to lysine sidechains of host proteins.^16-22^ Post translational modification of host proteins by Ub and UbL regulates the protein’s cellular localization, stability, or involvement in specialized responses such as antiviral immunity. PLpro recognizes the C-terminal RLRGG sequence of many Ub and UbLs acting as a deubiquitinase (DUB) towards these host proteins. DUB activity is hypothesized to cause dysregulation of both the initial inflammatory and subsequent interferon response. Substantial SARS-CoV-2-related mortality is associated with a dysregulated inflammatory response culminating in a cytokine storm.^23^ Thus, targeting PLpro is an attractive, strategy to inhibit viral replication and to prevent disruption of the host immune response to viral infection.

There has been a flurry of interest in drug repurposing as an expedited approach to address the COVID-19 pandemic, despite the lack of tangible success from repurposing approaches to the SARS-CoV and MERS-CoV outbreaks in 2002/3 and 2012/3, respectively. ^24,25^ Of the two essential cysteine proteases of SARS-CoV-2, 3CLpro is inhibited by many known cysteine protease inhibitors, the majority of which act via covalent modification of the active site cysteine and would seem to be a more amenable target for repurposing. The promiscuity of many human cysteine protease inhibitors has held back the progress of these agents into clinical use;^26^ however, off-target inhibition by calpain-1 inhibitors of cathepsin-L and 3CLpro may be opportunistically exploited, since cathepsin-L facilitates viral entry.^27^ Discovery of PF-008352313 as a covalent active-site-directed inhibitor of SARS-CoV 3CLpro in 2003 allowed the relatively rapid translation of this agent into clinical trials for SARS-CoV-2 in 2020.^28^

Like 3CLpro, PLpro from SARS-CoV-2 has 100% active site homology with the enzyme from SARS-CoV. In contrast to 3CLpro, there are very few potent inhibitors of SARS-CoV PLpro with efficacy validated experimentally; therefore, targeting PLpro with repurposed drugs is likely to be futile.^29-32^ A key reason for the lack of potent PLpro inhibitors is the absence of well-defined binding pockets at the S1 and S2 sites (Gly-Gly recognition). This presents severe challenges for inhibitor design and precludes a rapid drug discovery campaign.^33^ Furthermore, the relatively speedy resolution of the SARS-CoV outbreak in 2003 prematurely curtailed a number of early drug discovery campaigns. Notably, the resolution of crystal structures of the SARS-CoV PLpro apoenzyme by Ratia et al. demonstrated a conformationally flexible BL2 loop, remote from the active site cysteine, which could be stabilized by small molecule SARS-CoV PLpro inhibitors. ^34^ However, several of these inhibitors had reported poor metabolic stability, and the best, GRL0617, attained 14.5 µM potency in inhibition of host cell death to infectious SARS-CoV. ^35-37^

Utilizing our experience with SARS-CoV PLpro, we established a robust high-throughput screening (HTS) assay to identify SARS-CoV-2 PLpro inhibitors from both an unbiased and a targeted biased library. Based on observations with SARS-CoV PLpro, a low hit rate was expected; nevertheless, we identified CPI-169 as a validated and non-covalent SARS-CoV-2 PLpro inhibitor. We attempted to derive structure-activity relationships for synthetic analogues of CPI-169, which we computationally docked to the BL2 loop of PLpro; however, we were unable to obtain co-crystal structures. In parallel, we explored synthetic analogues of the known SARS-CoV PLpro inhibitor, GRL0619, for which a SARS-CoV PLpro co-crystal structure was available. We synthesized a chemical library of almost 100 compounds and identified nine compounds with low nanomolar potency against SARS-CoV-2 PLpro. Co-crystal structures of a number of these inhibitors with SARS-CoV-2 PLpro were obtained, revealing a novel “BL2 groove” formed by closing of the BL2 loop (blocking loop 2, AA266-271) and the palm domain. Structure-based optimization led to PLpro inhibitors with increased metabolic stability, low nanomolar potency against PLpro enzyme activity, a relatively slow dissociation rate, and low micromolar potency against infectious SARS-CoV-2. These novel compounds represent the most potent inhibitors of PLpro reported to date with activity translating to infectious virus assays in mammalian and human host cells.

## RESULTS

### High-throughput screening to identify inhibitors of SARS-CoV-2 Plpro

To discover novel chemical scaffolds that inhibit SARS-CoV-2 PLpro, we performed an HTS assay measuring PLpro protease activity with the short peptide substrate Z-RLRGG-AMC. Assay performance was excellent, with plate Z’ values ranging from 0.85-0.90. An unbiased ChemDiv library (10,000-compound SMART library subset excluding PAINS compounds) and a biased, annotated TargetMol Bioactive library (5,370 compounds) were screened. The Bioactive library contains 1,283 FDA-approved drugs, 761 drugs approved by regulatory bodies in other countries such as European Medicines Agency (EMA), and 3,326 advanced-stage developmental candidates. Compounds were screened against PLpro at a final compound concentration of 20 µM, which is a more stringent threshold than other contemporary screens of PLpro.^38^ Assay of PLpro in the presence of 5 mM DTT, as reducing agent and electrophile trap, strongly biases against reactive (electrophilic and redox) hits.

The 28 hit compounds from HTS were counter-assayed to remove false positives associated with signal interference and then further pruned to remove frequent hitters and known redox cyclers (**Figure 1 & Figure S2A, B**). A set of five compounds producing >40% inhibition of SARS-CoV-2 PLpro activity along with the SARS-CoV PLpro inhibitor GRL0617 were selected for follow-up 8-point dose-response assays (**Figures 2A**). All six active compounds were also tested in an orthogonal binding assay using surface plasmon resonance (SPR) (8-point titration) (**Figure 2B**). Only GRL0617 and CPI-169 inhibited PLpro with IC_50_ < 10 µM in the primary enzyme inhibition assay (IC_50_ values of 1.6 µM and 7.3 µM, respectively (**Figure 2A**). These values compare well with the binding affinities measured by SPR: GRL0617 K_D_= 1.9 µM, CPI-169 K_D_ = 10.2 µM (**Figure 2C**).

**Figure 1.**
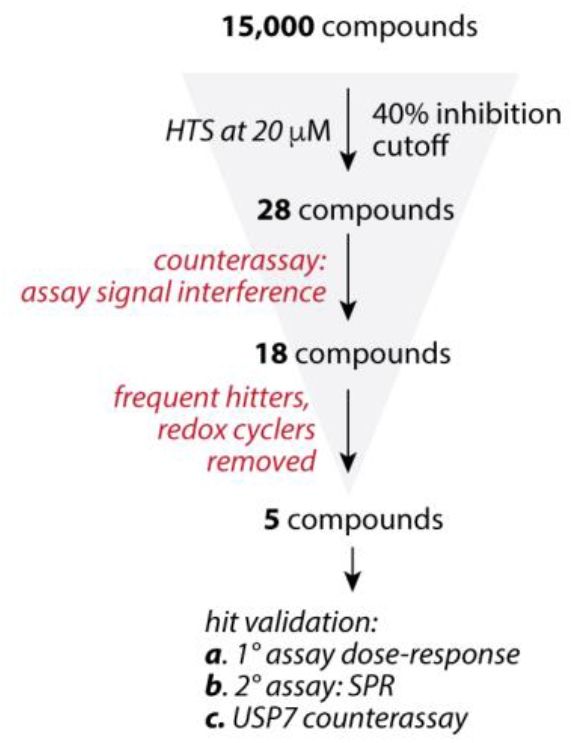
HTS for SARS-CoV-2 PLpro inhibitors. Hit triage and validation workflow.

**Figure 2.**
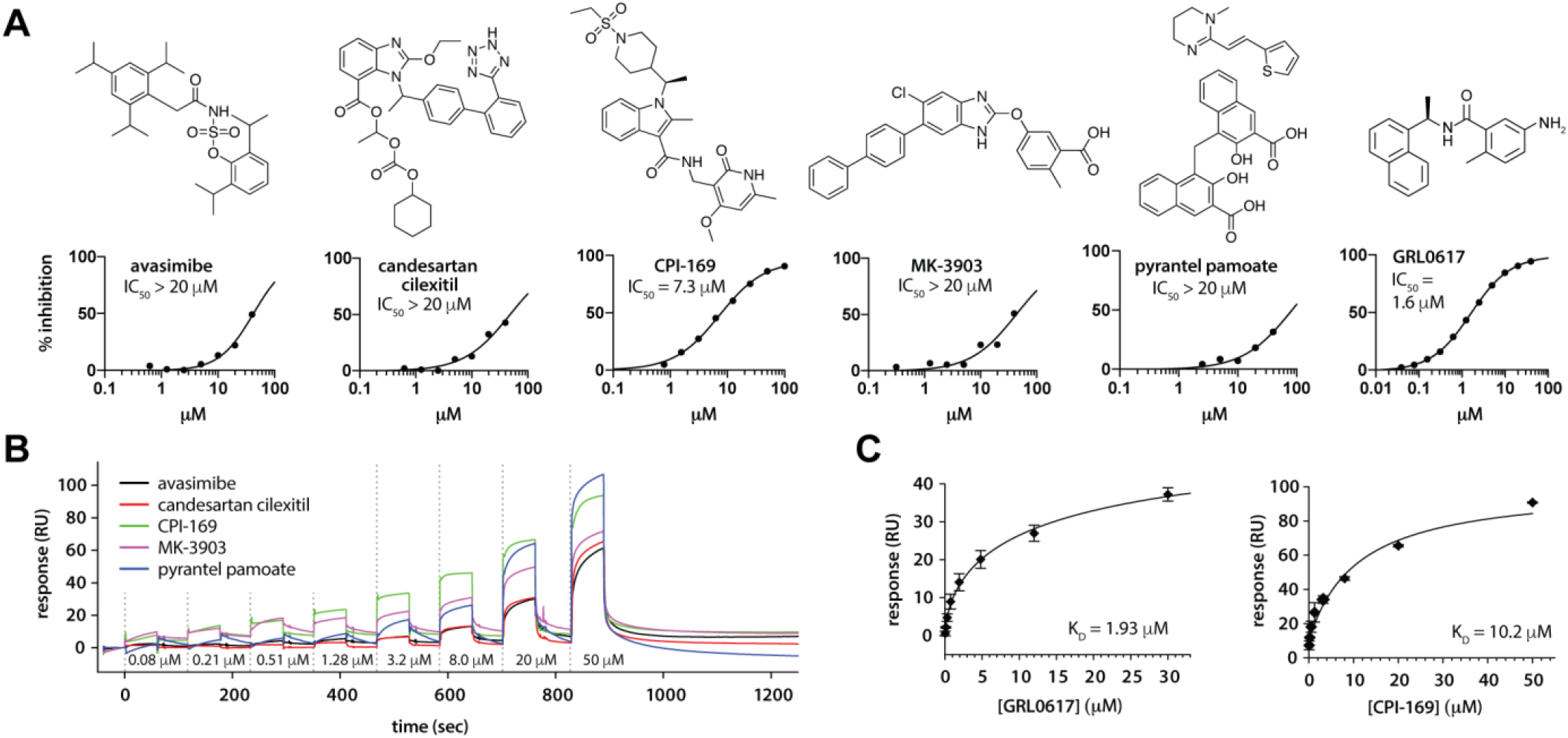
HTS for SARS-CoV-2 PLpro inhibitors. **A)** Chemical structures and dose-dependent SARS-CoV-2 PLpro inhibition in the primary HTS assay. **B)** Overlaid SPR sensorgrams of the single-cycle kinetics for HTS hits (0.08 µM to 50 µM, 2.5-fold dilutions). **C)** Binding of GRL0617 and CPI-169 to SARS-CoV-2 PLpro as measured by SPR.

PLpro has DUB enzyme activity; therefore, human DUBs represent off-targets for any PLpro inhibitor. To confirm that hits were selective for SARS-CoV-2 PLpro over host cell DUBs, the catalytic domain of human USP7, the closest structural homolog of PLpro, was used as an additional counter-assay. Consistent with the initial findings, GRL0617 did not inhibit USP7-catalyzed Ub-AMC hydrolysis.^29^ Similarly, CPI-169 was not able to inhibit USP7 up to a concentration of 30 µM (**Figure S2C**). This data confirmed and validated GRL0617 and CPI-169 as selective SARS-CoV-2 PLpro inhibitors.

### Structure-based design of PLpro inhibitors

With these two inhibitors in hand, we embarked on a medicinal chemistry campaign to optimize both GRL0617 and CPI-169 (SAR for CPI-169 will be reported in a separate paper). As shown above, GRL0617 displayed only modest potency against SARS-CoV-2 PLpro. Furthermore, this family of compounds was reported to be unstable to metabolism by liver cytochrome P450s, probably due to the presence of aniline and naphthalene groups, which are known culprits for rapid metabolism, precluding use as *in vivo* probe compounds.^32,39^ Any modifications to the GRL0617 series would need to incorporate improved potency and metabolic stability.

SARS-CoV-2 PLpro has 83% sequence identity to SARS-CoV PLpro and 100% identity at the active site; therefore, the GRL0617:PLpro (SARS-CoV) co-crystal structure (PDB: 3E9S) can be used to guide initial structure-based optimization of this series. The PLpro monomer is comprised of four distinct domains, including an N-terminal ubiquitin-like (Ubl) domain (first 62 residues) and an extended right-hand architecture with distinct palm, thumb, and finger domains (**Figure 3A**). The active site formed by the catalytic triad, Cys111, His272, and Asp286 (SARS-CoV-2 PLpro numbering, PDB: 7JRN), sits in a solvent-exposed cleft at the interface of the thumb and palm domains. Binding of host and viral protein substrates is controlled by the flexible β-hairpin BL2 loop, which contains an unusual beta-turn formed by Tyr268 and Gln269 (**Figures 3A, B**), controlling access to the active site.

**Figure 3.**
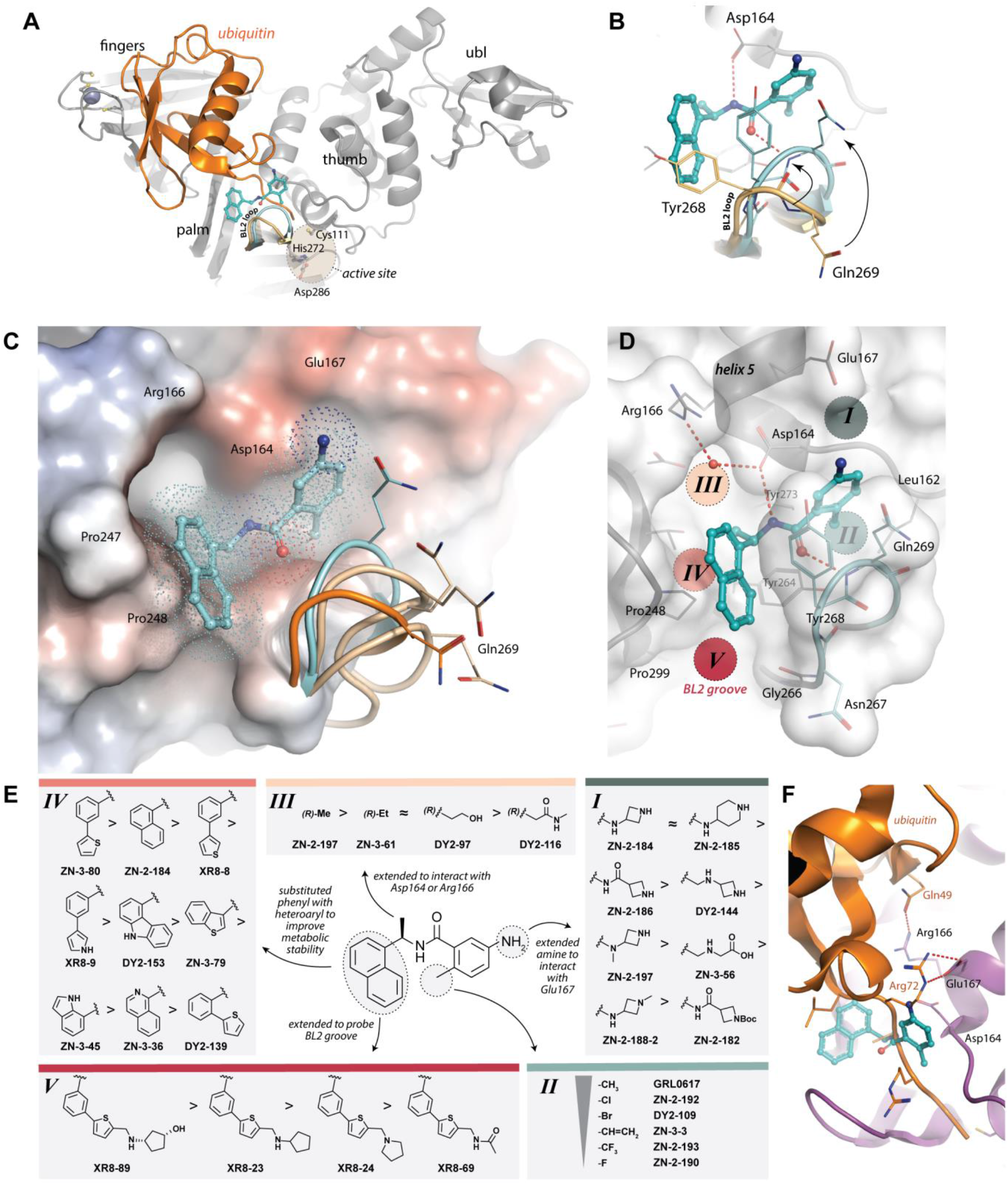
Structure-guided design of SARS-CoV-2 PLpro inhibitors based on GRL0617. **A)** Overall structure and domain organization of PLpro and the PLpro-ubiquitin complex. GRL0167 is shown in cyan. **B)** Twisting of BL2 loop induced by GRL0617 binding. The conformation of the ubiquitin-bound BL2 loop is shown in pale orange; the GRL0167-bound loop is shown in cyan. **C)** BL2 loop conformational flexibility. The structure of the GRL0617-bound PLpro (electrostatic surface representation) and associated BL2 loop (cyan cartoon) superimposed with Ub-bound (orange; pdb: 6XAA) and apo structures (wheat; pdbs 6WZU, 7D47, 7JCD). Gln269 is shown for reference. **D)** Structural detail of *Sites I-V* targeted by drug design; **E)** Brief summary of structure activity relationship of selected compounds (full SAR in shown in Tables S1-S5); **F)** PLpro Glu167 (magenta) interacts with Arg72 of ubiquitin (orange) in *Site I*. GRL0617 is aligned and shown in cyan.

Superposition of crystal structures of the SARS-CoV-2 PLpro apoenzyme (PDBs 6WZU, 7D47, 7JCD) highlights the flexibility of the BL2 loop (**Figure 3C)**. The binding of GRL0617, presumably by induced-fit, requires reorganization of the PLpro secondary structure, and thus the association rate of the ligand is anticipated to be slower than the rate of diffusion. In SPR experiments, the association rate was measured to be 1.8 × 10^5^ M^-1^s^-1^, which is significantly slower than the expected 1×10^9^ M^-1^s^-1^ rate of diffusion-controlled association. The unique structural reorganization of the BL2 loop, in part, explains the low hit rate from our HTS campaign (as also reported by others^40^) and represents a challenge and an opportunity for developing potent, selective small molecule PLpro inhibitors.

In the GRL0617:PLpro (SARS-CoV) co-crystal structure, the ligand stabilizes the closed BL2 loop, blocking access to the active site; thus, unusually for a cysteine protease inhibitor, GRL0617 does not interact with the active site cysteine and the closest point of contact is 7.6Å distant. We hypothesized that CPI-169 also stabilizes the BL2 loop; however, we were unable to obtain co-crystal structures of CPI-169 or its analogues with SARS-CoV-2 PLpro to guide optimization.

### Optimization of novel benzamide PLpro inhibitors

GRL0617 forms two key hydrogen bonding interactions with the mainchain nitrogen of Gln269 and sidechain of Asp164 in PLpro (**Figure 3B**), causing the BL2 loop to adopt a twisted conformation that narrows the solvent-exposed surface and exposes a hydrophobic binding site. To test the importance of this hydrogen bonding interaction in regulating the BL2 loop conformation, we synthesized two tool compounds reducing the amide to amine (DY2-64) or replacing the amide with a sulfonamide bioisostere (DY3-63). Both modifications led to a sharp decline in potency, therefore the benzamide was conserved moving forward (**Table S1**).

A detailed analysis of residues within the GRL0617 binding site of PLpro revealed five potential regions that we hypothesized could be targeted to increase affinity and potency for BL2-binding ligands (*I-V*, **Figure 3D**,**E**). Compounds extended into *Site I* explore potential interactions with Glu167, which forms electrostatic contacts with the Arg72 of ubiquitin in the Ub:PLpro SARS-CoV co-crystal structure (PDB: 4MM3) (**Figure 3F**).^19^ We envisioned that a basic amine appended to the aniline group would capture this interaction and improve binding affinity. A library of 16 compounds was synthesized to identify suitable basic side chains (**Figure 3E**). The azetidine-substituted ZN2-184 was the most potent analogue targeting *Site I*, with a two-fold improvement relative to GRL0617, which correlated with affinity measured by SPR. The increase in affinity and potency was also accompanied by a twofold increase in rate of association (**Figure 4A-D**).

**Figure 4.**
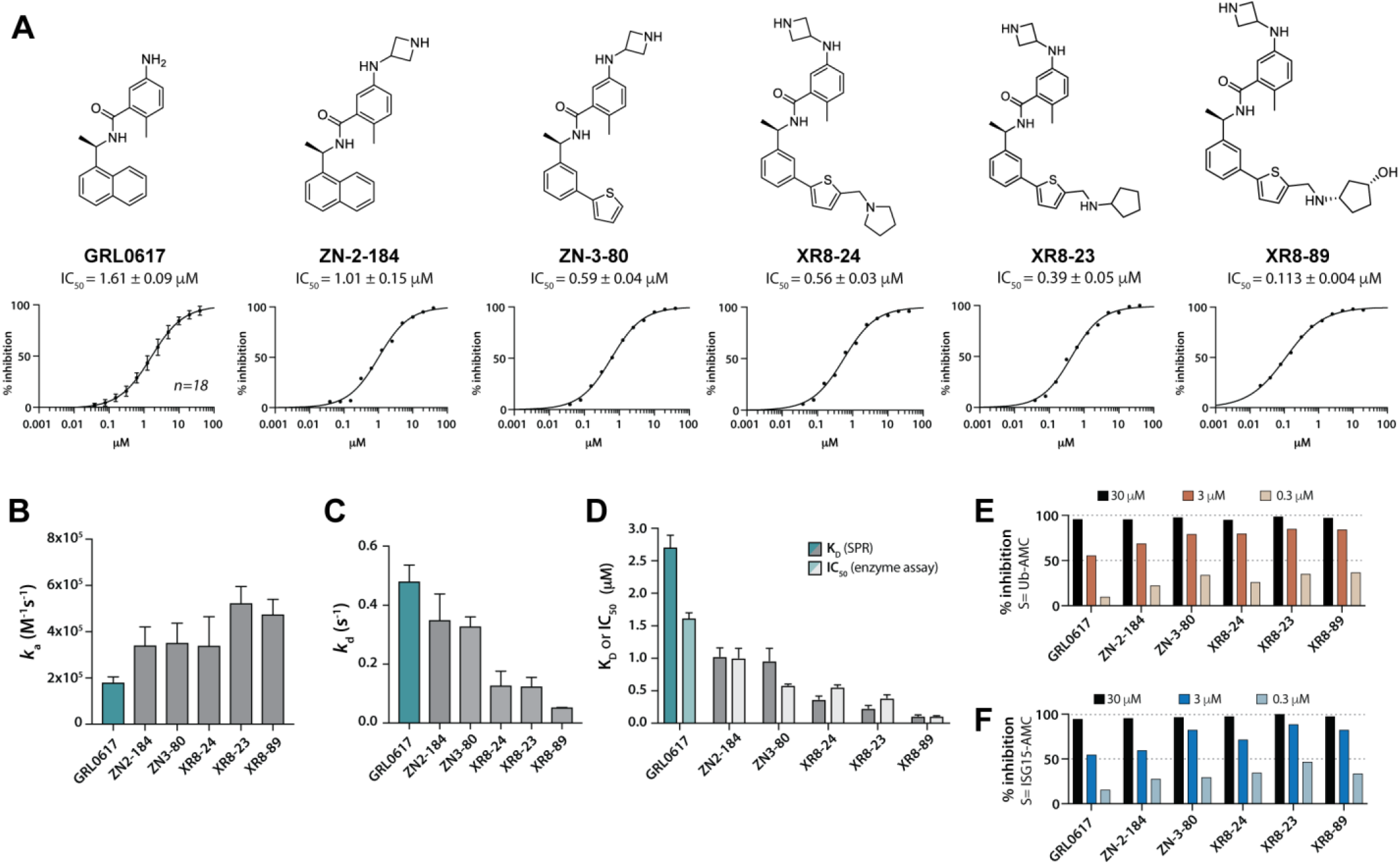
**A)** Chemical structures and dose responses of the most potent PLpro inhibitors. **B)** Association rates and **C)** dissociation rates of PLpro inhibitors as determined by SPR. **D)** Comparison of K_D_ measured by SPR and IC_50_ of enzyme inhibition assay of select PLpro inhibitors. Inhibition of **E)** deubiquitinating and **F)** de-ISGylating activities of PLpro by select inhibitors at three compound concentrations.

*Site II* is located at the S3 site of the substrate-binding channel, which is formed by the BL2 loop, helix 5, and neighboring hydrophobic residues Tyr264, Tyr273, and Leu162 (**Figure 3D**). Small hydrophobic moieties such as a halogen or trifluoromethyl group were synthesized to probe the hydrophobic interaction at this site (Figure 3E, **Table S2**). Interestingly, small substitutions such as methyl to fluorine at *Site II* led to a dramatic decrease in potency. Only bromo and chloro substituents did not significantly right-shift potency. Attempts to make fused-ring indole analogs to replace aniline also did not lead to any improvement in potency.

*Site III* is positioned next to the charged side chains of Arg166 and Asp164 (**Figure 3D**). Arg166 forms an electrostatic interaction with Asp164 via its charged guanidino group, leaving the other guanidine nitrogens available for hydrogen bonding interactions. In the Ub:PLpro complex, this interaction is captured by hydrogen bonding to the Gln49 of ubiquitin (**Figure 3F**). To exploit hydrogen bonding with Arg166 at *Site III*, we synthesized analogues modified at the: 1) 2-napthelene position; 2) α-methyl position; and 3) aniline nitrogen, all without success (**Tables S2-4**). Indeed, minor modifications, such as ZN3-36 with a 2-isoquinoline, designed to engage with a structurally conserved water molecule at *Site III*, significantly lost activity against PLpro (IC_50_ = 56 µM, **Figure 3D, Table S4**). Conformational minimization (B3LYP/6-31G* with a polarizable continuum model for aqueous solvation) indicated a dihedral angle of 27.9° between the amide and isoquinoline planes of ZN3-36 (**Figure S3)**. This angle is significantly different from that seen in the crystal structure of GRL-0617 (81.7°, PDB: 7JRN), which may highlight the importance of maintaining a dihedral angle ∼ 90° for optimal hydrogen bonding with the BL2 loop (**Figure S3**).

Extending from the α-methyl position also proved to be futile. A minor ethyl modification led to a significant decrease in potency (ZN3-61); further expansion from this position resulted in almost completely inactive compounds such as DY2-97 and DY2-116 (**Table S3**). Only ZN3-56 with a glycine tail extended from the aniline site could engage with *Site III* and led to a slight improvement of potency over GRL0617. The proposed binding model of ZN3-56 predicts an electrostatic interaction with Arg 166 (**Figure S4**); however, this incremental improvement in potency suggests that electrostatic stabilization at this highly solvent-exposed region is counterbalanced by a solvation penalty. No further exploration of *Site III* interactions was attempted.

### Scaffold hopping: naphthalene ring replacement

Although the active sites of PLpro from SARS-CoV and SARS CoV-2 are identical, there are several amino acids that differ between the two enzymes, which are within approximately 10Å of the binding site of BL2-stabilizing ligands GRL-0617 (e.g., E251/Q250 and N263/S262). It is reasonable to assume that these substitutions might subtly alter ligand binding to the BL2 loop in SARS-CoV-2 PLpro versus SARS-CoV. This observation alone supports exploration of scaffolds to replace the naphthalene of GRL0617. Retaining the essential geometry between the benzamide and naphthalene rings should be possible using heteroaryl or bi-aryl group replacements. Replacement of the naphthalene ring is also anticipated to reduce metabolic instability.^30^ Modifications at this site should also allow favorable interactions with *Site IV* (**Figure 3D**). Fused heteroaryls such as benzothiophene, indole, and carbazole with various linkages were prepared and tested (**Table S4**); however, most modifications weakened or lost activity. Only the 3-benzothiophene (ZN3-79) and the carbazole-based (DY2-153) analogues showed comparable potency to GRL0617 (IC_50_ = 1.9 µM and 1.8 µM, respectively; **Table S4**). In contrast, the biaryl analogues not only retained the activity of GRL0617 but also gained potency. Both 2-phenylthiophene (ZN-3-80; IC_50_ = 0.59 µM) and 3-phenylthiophene (XR8-8; IC_50_ = 1.3 µM) demonstrated enhanced potency (Table S5). Importantly, ZN3-80, the most potent analog in this subset, was found to be more stable than GRL0617 in human liver microsome stability assays (**Table S6**), which encouraged us to explore further optimization.

### Engaging the BL2 groove decreases inhibitor off-rate and improves potency

Examination of crystal structures identified a ligand binding site, coined the “BL2 groove” (*Site V*, **Figure 3D**), which is positioned at the N-terminal side of the BL2 loop and features hydrophobic residues such as Pro248 and Pro299 and potential hydrogen bonding partners such as the backbone amide of Gly266. With an optimized 2-phenylthiophene (ZN3-80) in hand, we explored derivatization to exploit further interactions with the BL2 groove, synthesizing 22 compounds. Interestingly, nine compounds in this family significantly impacted potency, improving IC_50_ to below 500 nM (**Figure 4A; Table S5**).

We hypothesized that once bound, these extended ligands interacting with *Sites I-V* might have slower off-rates because of the increased conformational reorganization of the BL2 loop required for ligand release (Figure S5 & Table S8). Association and dissociation rates were measured by SPR (**Figure 4B**). The extended ligands, designed to engage the BL2 groove, showed a marked decrease in dissociation rates (**Figure 4C**). For example, both XR8-23 and XR8-89, with basic amine side chains extending from the thiophene of ZN3-80, produced 4-fold and 9-fold reductions in off rates compared with GRL0617, respectively; suggesting that BL2 groove engagement is a novel strategy for development of potent PLpro inhibitors with slower off-rates.

To confirm that the improved analogs are capable of inhibiting the DUB activity of SARS-CoV-2 PLpro, we studied both Ub-AMC (**Figure 4E**) and ISG-15-AMC (**Figure 4F**) as substrates for the enzyme at three inhibitor concentrations. Complete ablation of DUB activity was observed with all tested inhibitors at 30 µM and at the approximate IC_50_ concentration for GRL0617, all novel inhibitors produced greater inhibition of DUB activity. The data support the ability of these novel inhibitors to block SARS-CoV-2 PLpro-mediated deubiquitination and deISGylation of host proteins involved in host immune response.

### Co-crystal structures of XR8-23 and XR8-89 with SARS-CoV-2 PLpro

To examine the binding mode of our novel PLpro inhibitors and test the proposed binding site hypotheses, we obtained co-crystal structures of XR8-24, XR8-65, XR8-69, XR8-83, and XR8-89 complexed with SARS-CoV-2 PLpro. Superposition of the ligand-bound structures shows all inhibitors enforcing the same binding mode of the BL2 loop (**Figure S6**). Focusing on the co-crystal structures of XR8-24 and XR8-89, we observe that the azetidine ring extends into *Site I* to within 3 Å of Glu168, likely engaging in the postulated electrostatic stabilizing interaction (**Figure 5A, B**). The amide group of XR8-24 and XR8-89 is aligned closely with that of GRL0617 in SARS-CoV-2 PLpro (PDB: 7JRN) with the expected: i) carbonyl hydrogen bonding to the mainchain of Gln269 on the BL2 loop; and ii) amide nitrogen hydrogen bonding to Asp164 of helix 5. In *Site IV*, the 2-phenylthiophene of the ligand retains the T-shaped pi-interaction with Tyr268, as seen for the naphthalene ring of GRL0617; however, there is a shift in the biaryl ring of XR8-24 relative to the naphthalene of GRL0617, suggesting that the biaryl substituent maximizes interactions that may not be present in PLpro from SARS-CoV. These interactions place the thiophene firmly in the BL2 groove (*Site V*), where it takes part in van der Waals interactions with residues surrounding the cavity (Pro248, Tyr264, Tyr268; **Figure 5D**).

**Figure 5.**
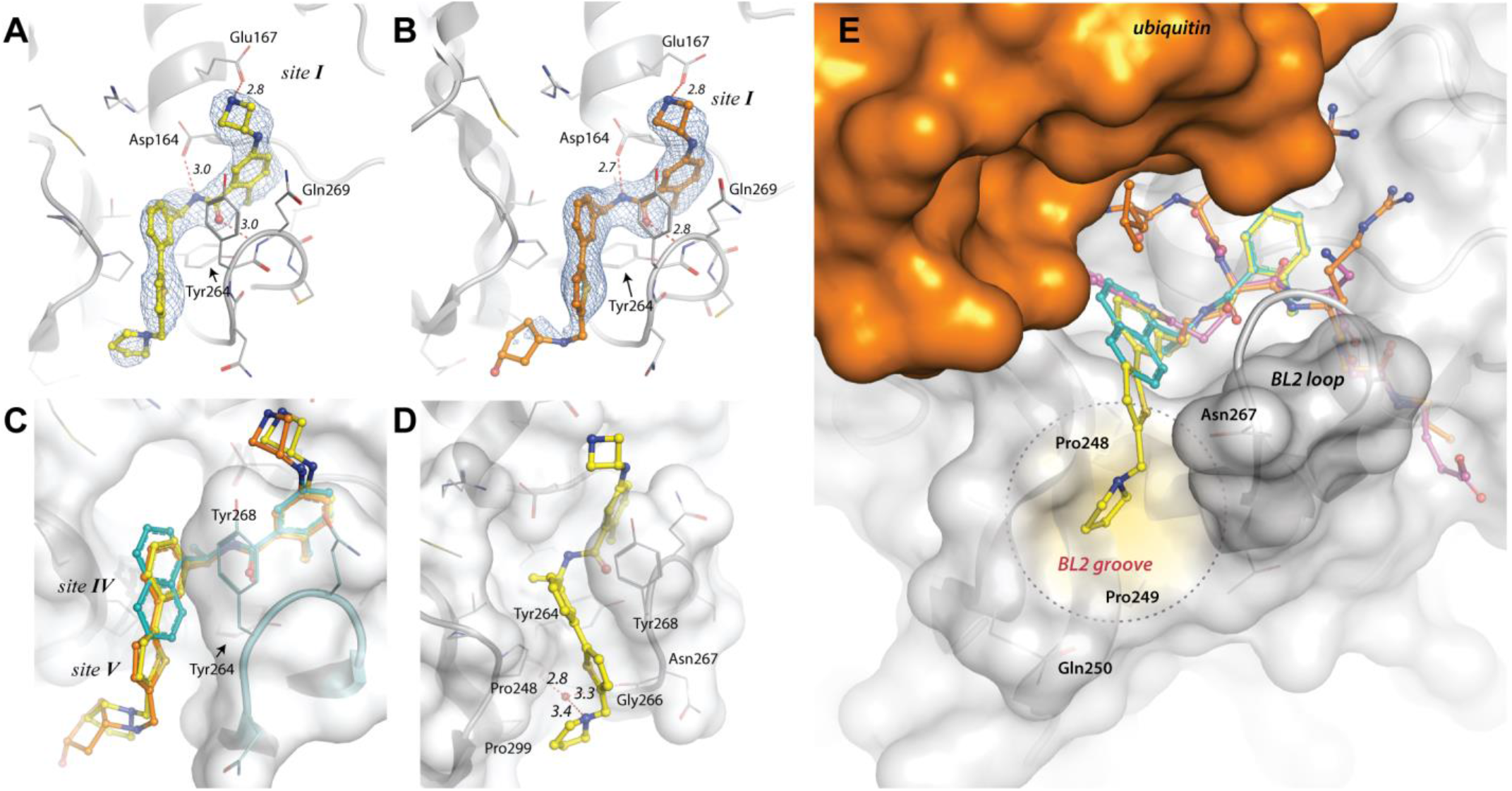
Structural details of SARS CoV-2 PLpro complexed with XR8-24 and XR8-89. 2F_o_-F_c_ electron density maps of A) XR8-24 and B) XR8-89. The maps are shown as blue mesh and are contoured at 1 sigma around the inhibitors. Hydrogen bonds are indicated by dashed lines, with distances (in Å) indicated in italics. C) Superposition of SARS-CoV-2 PLpro-bound GRL0617 (cyan; pdb 7JRN) with XR8-24 (yellow) and XR8-89 (orange). D) Interaction of XR8-24 with the BL2 groove. E) A comparison of PLpro ligand and inhibitor binding surfaces on PLpro. The surface of the body of ubiquitin is shown in orange and its 5 C-terminal residues are shown as orange sticks (pdb 6XAA). GRL0617 is shown in cyan (pdb 7JRN) and a covalent peptide-based inhibitor (pdb 6WUU) is shown in magenta. XR8-24 is shown in yellow, and the binding surface unique to XR8-24 and close analogs is highlighted by a yellow circle.

The “tail” of both XR8-24 and XR8-89, which we postulated to provide interactions with the BL2 groove in these co-crystal structures sits perpendicular to the thiophene and adjacent to the body of the protein near Pro248 and Pro299. The lack of electron density for the tail of XR8-89 (**Figure 5B**), also observed for the co-crystal structures of XR8-65, XR8-69 and XR8-83 indicates that this region is largely disordered, which may be due in part both to high solvent-exposure and crystal packing forces induced by the proximity to a second symmetry-related monomer. However, the tail of XR8-24 is better defined, with the pyrrolidine ring forming a putative water-mediated hydrogen bond to the mainchain carbonyl oxygens of Tyr264 and Gly266 (**Figure 5D**). The new co-crystal structures show that engagement of the BL2 groove contributes to binding affinity and we emphasize that this binding interaction is unique and is not observed in structures of SARS CoV-2 PLpro in complex with ubiquitin or ISG15, nor with any known PLpro inhibitors (**Figure 5E**).

### Improved potency translates to improved antiviral activity

Of the potent, novel PLpro inhibitors, selection for antiviral testing was based on the following rationale: XR8-89 demonstrated highest potency for PLpro inhibition (IC_50_=113 nM); XR8-23 demonstrated a high association/dissociation rate ratio; and XR8-24 yielded a superior co-crystal structure. No toxicity was observed under assay conditions in Vero E6 cells for these compounds at < 50 µM; therefore, we evaluated all three compounds compared with GRL0617 in a plaque formation assay using the SARS-CoV-2 USA/WA1/2020 strain.

Vero E6 cells are known to express high levels of efflux transporter proteins.^41^ In a report on the 3CLpro inhibitor, PF-00835231, currently in clinical trials for COVID-19, co-treatment with the dual P-gp/BCRP inhibitor, CP-100356, was required to elicit antiviral activity in Vero E6 cells. ^42^ In Caco-2 cells, XR8-24 demonstrated a high efflux ratio (**Table S7**), implying efficient P-gp mediated efflux; therefore CP-100356 co-treatment (1.5 µM) was used to test antiviral activity in Vero E6 cells. Consistent with reports on PF-00835231, CP-100356 itself had no antiviral activity and at the tested concentrations did not influence cytotoxicity alone or in combination with our PLpro inhibitors. Consistent with contemporary reports on GRL0617,^21^ an EC_50_ of 21.7 ± 1.6 µM was measured against infectious SARS-CoV-2 in an eight-point dose response assay. Both XR8-23 and XR8-24 were significantly more potent than GRL0617 with EC_50_ measured at 2.8 ± 0.4 µM and 2.5 ± 1.9 µM, respectively (**Figure 6A** and **Figure S7**). XR8-89 also demonstrated superior antiviral potency to GRL017; however, antiviral potency did not correlate with the superior potency of this inhibitor in biochemical assays. The lack of observable cytotoxicity for XR8-89 might indicate attenuated cell permeability as a cause of lower antiviral potency.

**Figure 6.**
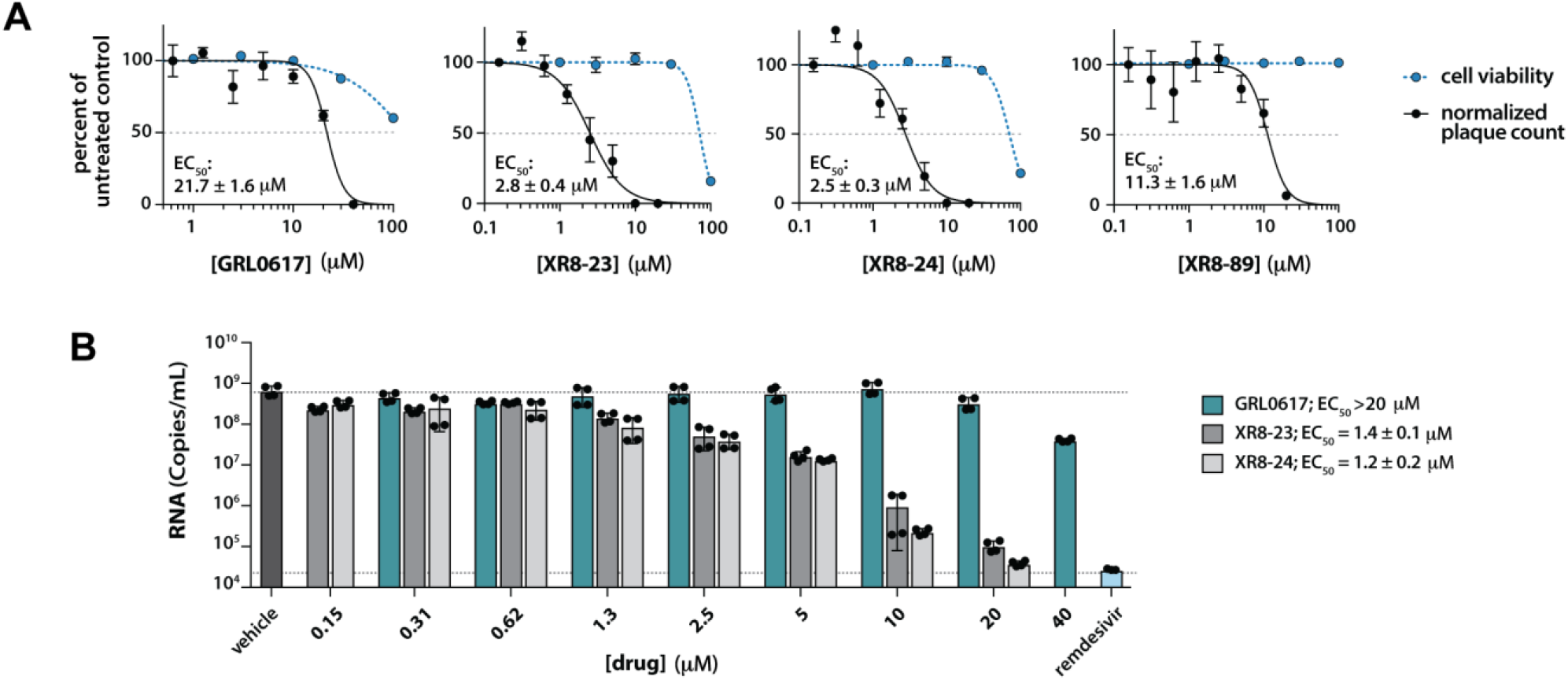
Improved PLpro inhibitors shows potent antiviral efficacy. **A**) Plaque reduction of SARS-CoV-2 infected Vero E6 cells at MOI = 0.0001 treated with GRL0617, XR8-23, XR8-24, and XR8-89 from 20 µM to 0.156 µM in the presence of 1.5 µM CP-100356. Cytotoxicity in Vero E6 cells was measured by CellTiter-Glo assay **B**) To measure reduction in virus yield, A549-hACE2 cells were infected with MOI = 0.01 of SARS-CoV-2 cultured in Vero E6 cells with and without various concentrations GRL0617, XR8-23, XR8-24 after 48 hours, supernatants were harvested, RNA isolated and quantified by RT-qPCR. The data show mean ± S.D.

The two most potent antiviral agents in Vero E6 cells, XR8-23 and XR8-24, were tested and compared to GRL0617 and remdesivir, an FDA-approved COVID-19 antiviral agent, in the human lung epithelial A549 cell line stably overexpressing the human ACE2 receptor. Although monkey Vero E6 cells are a standard model for antiviral testing, a human cell line provides an orthogonal and more relevant model system. Viral RNA was assayed by RT-qPCR as a measure of replication of infectious SARS-CoV-2 USA/WA1/2020. The assay was conducted in the absence of CP-100356 and cytotoxicity was not observed under assay conditions at < 50 µM for XR8-24 and < 10 µM for XR8-23 (Figure S8). The antiviral activity of novel PLpro inhibitors was markedly superior to that of GRL0617 in this model system (**Figure 6B)**. Both XR8-23 and XR8-24 were again significantly more potent than GRL0617 (IC_50_ > 20 µM) with IC_50_ measured at 1.4 ± 0.1 µM and 1.2 ± 0.2 µM, respectively. By unpaired nonparametric t-test: 1) the effect of treatment with XR8-23 and XR8-24 (1.3 µM) was significantly different from vehicle control; and 2) the effect of treatment with XR8-24 (20 µM) was not significantly different from that of remdesivir (10 µM).

## DISCUSSION

The pathogenesis of the SARS-CoV-2 infection is characterized by a strong dysregulation of the innate immune and the type I interferon (IFN-I) responses.^43^ The viral protein, PLpro, represents an excellent therapeutic target owing to multi-functional roles: i) in mediating viral replication via processing of the viral polyprotein; and ii) in reversing host-mediated post-translational modifications in response to viral infection via its actions as a DUB. The DUB enzyme activity of PLpro is responsible for removing ubiquitin chains and the ISG15 ubiquitin-like (Ubl) modification from host proteins. ISGylation of proteins is induced during viral infection as a host antiviral signaling mechanism.^44^ Interestingly, despite the close homology of PLpro from SARS-CoV-2 and from the original SARS-CoV coronovirus, PLpro has been claimed differentially to modulate the host immune system: specifically, it is reported that SARS-CoV-2 PLpro preferentially cleaves ISG15, whereas PLpro from SARS-CoV predominantly targets ubiquitin chains.^20,45^ In addition to Ub- and Ubl-modified host proteins, the autophagy-activating kinase, ULK1, is also a substrate for PLpro, cleaving the N-terminal kinase domain from a C-terminal substrate recognition region to disrupt autophagy during early viral replication.^46^ Pharmacological inhibitors of PLpro are needed to probe the effects of PLpro proteolytic activity on host cell immune response and autophagy; however, more urgent is the need for PLpro inhibitors that effectively block viral replication, since many SARS-CoV-2 gene products in addition to PLpro have immune modulatory effects.

Towards developing potent inhibitors, we developed a robust high-throughput assay for SARS-CoV-2 PLpro using Z-RLRGG-AMC to screen chemical libraries including FDA-approved drugs and molecules in clinical trials (15,370 molecules). Consistent with contemporary reports,^38,40,47^ an extremely low hit rate was observed, which makes repurposing of approved drugs as therapeutically useful PLpro inhibitors problematic. As recent reports have noted, the experimental data is not in accordance with *in silico* repurposing predictions:^48,49^ for example, isotretinoin reportedly in clinical trials for COVID-19 as a PLpro inhibitor, ^*46*^ was inactive in our screen. We did identify avasimibe and candesartan as weak PLpro inhibitors; however, only GRL0617 and CPI-169 (initially developed as an EZH2 inhibitor) were validated as hits with potency/affinity < 10 µM in both enzymatic assay and SPR binding assays. CPI-169 represents a novel chemical scaffold and a novel addition to the very limited PLpro inhibitor chemotypes identified in the literature.^33,50^

Given the availability of co-crystal structures of GRL0617:PLpro (SARS-CoV-2), structure-guided drug design was pursued exploring this chemical scaffold. Five binding sites on PLpro were explored by modification of the benzamide scaffold to identify additional interactions to increase inhibitor potency. *Site I* contains Glu167, which forms a salt bridge with Arg72 of ubiquitin in the Ub:PLpro complex. This electrostatic interaction was captured with addition of a basic amine side chain in novel inhibitors to yield a significant increase in binding and potency. Additional modifications to engage with the S1 or S2 sites in the channel leading to the catalytic site were unsuccesful, consistent with recent findings profiling substrate specificity using a combinatorial library.^51^ *Site III* is defined by Arg166, which forms a hydrogen bonding interaction with Gln49 of ubiquitin; however, none of the modifications designed to mimic this interaction increased the affinity of inhibitors for PLpro. *Site III*, therefore, remains to be exploited in future work.

The BL2 groove is a new binding site identified in the process of inhibitor optimization, which was confirmed and validated by obtaining SARS-CoV-2 PLpro co-crystal structures. This BL2 groove is not involved in the binding of any PLpro substrates, such as Ubs and Ubls, by the enzyme. Novel inhibitors, such as XR8-23 and XR8-24, modified with BL2-interacting side chains, showed both improved binding affinity and slower off-rates, suggesting that BL2 groove interactions can yield more efficacious PLpro inhibitors. Gratifyingly, these enhanced biochemical properties translated to antiviral efficacy against infectious SARS-CoV-2 (USA/WA1/2020) in Vero E6 green monkey kidney epithelial cells and A549 human lung epithelial cells. The low micromolar potency observed in inhibition of viral plaque formation was superior to GRL0617 and suggests that optimization of PLpro inhibitors as therapeutic agents for SARS-CoV-2 is feasible. Vero E6 cells are highly susceptible to the cytopathic effects of SARS-CoV-2 infection in contrast to many human cell lines.^52^ The observations in a human lung epithelial cell line of inhibition of SARS-CoV-2 viral replication is therefore very promising. Novel PLpro inhibitors were markedly more efficacious than GRL0617, with significant suppression of viral RNA at low micromolar concentrations.

PLpro inhibitors such as XR8-23 and XR8-24 provide an opportunity to study combination therapy with FDA-approved RdRp inhibitors such as remdesivir, or 3CLPro inhibitors such as PF-00835231, now in Phase I/II clinical trials. Genotyping of SARS-CoV-2 virus strains circulating worldwide has identified multiple recurrent non-synonymous mutations in the receptor-binding domain (RBD) of the spike protein. For example, the SARS-CoV-2 B.1.1.7 strain identified in London contains a N501Y mutation in the RBD domain. Variants with multiple mutations in the spike protein pose a risk of resistance to current FDA-approved vaccines and therapeutic antibodies; mutations in the cysteine proteases 3CLpro and PLpro have not been reported.

In conclusion, we identified a new drug-like PLpro inhibitor chemotype, CPI-169, adding to the very limited examples of PLpro inhibitor scaffolds. Guided by new SARS-CoV-2 PLpro co-crystal structures, we designed novel non-covalent PLpro inhibitors that exhibited superior nanomolar potency and inhibited PLpro DUB activity, without inhibiting human DUBs. Biochemical potency, affinity, and slow off-rates translated to low micromolar potency against infectious SARS-CoV-2 in primate and human cell lines. Non-covalent inhibitors that stabilize the BL2 loop by induced fit do not occupy the active site of PLpro and published data on SARS-CoV PLpro inhibitors showed relatively weak potency against infectious virus; therefore, the therapeutic relevance of this approach in SARS-CoV-2 antiviral therapy was problematic. We have shown that BL2-stabilizing PLpro inhibitors have therapeutically relevant activity against SARS-CoV-2.

We synthesized almost 100 analogues during structure-based optimization, identifying the novel BL2 groove as an important ligand binding site. The newly identified non-covalent inhibitors described herein are the most potent SARS-CoV-2 PLpro inhibitors reported to date, with improved potency and metabolic stability. The infectious virus data suggests that administration in combination with remdesivir or 3CLpro inhibitors may be therapeutically beneficial. Moreover, potent PLpro inhibitors such as XR8-23 and XR8-24 represent chemical probe tool compounds to study the details of PLpro-mediated disruption of host immune response and autophagy; and their contribution to COVID-19 infection and progression, including “long-COVID” and potential genetic bias.^53,54^

## MATERIALS AND METHODS

### SARS-CoV-2 PLpro expression and purification

pET11a vector containing SARS-CoV-2 PLpro protein (pp1ab aa 1564-1878) with N-terminal, TEV-cleavable His-tag was transformed into BL21(DE3) cells and maintained in media containing 100 ug/mL carbenicillin. Protein expression was induced using an auto-induction protocol modified from Studier et al.^55^ Briefly, 1 mL day cultures were used to inoculate a 2L flask of 500 mL of Super LB containing 100 ug/mL carbenicillin. Cells were grown for 24h at 25°C and then harvested by centrifugation. All steps of SARS-CoV2 PLpro purification were performed at 4°C. Protein yield at each step was monitored by Bradford assay using BSA as a standard. Frozen cells pellets were lysed by sonication in Buffer A (50 mM HEPES, pH 8, 0.5 M NaCl) containing 10 ug/mL lysozyme. The lysate was clarified by centrifugation and loaded onto a 2-mL HiTrap Talon crude column equilibrated with Buffer A. Bound His6-PLpro was eluted with a linear gradient of 0-150 mM imidazole in Buffer A, and fractions containing His6-PLpro were pooled and exchanged into cleavage buffer (20 mM Tris-HCl pH 8.5, 5 mM DTT, 0.5 mM EDTA, 5% glycerol). A 1:100 molar ratio of TEV protease to PLpro was incubated at 4°C overnight to cleave the His6-tag. To remove the tag and TEV protease, the reaction was loaded onto a UNO-Q column equilibrated with 20 mM Tris HCl, pH 8.5, 3 mM DTT. Cleaved PLpro eluted first in a gradient from 0-150 mM NaCl over 20 column volumes. Fractions containing cleaved PLpro were pooled and concentrated to 12 mg/mL, frozen in liquid nitrogen, and stored at −80 °C.

### PLpro primary assay

The PLpro primary assay, which measures protease activity with the short peptide substrate Z-RLRGG-AMC (Bachem), was performed in black, flat-bottom 384-well plates containing a final reaction volume of 50 μL. The assays were assembled at room temperature as follows: 40 µL of 50 nM PLpro in Buffer B (50 mM HEPES, pH 7.5, 0.1 mg/mL BSA, 0.01% Triton-X 100, and 5 mM DTT) was dispensed into wells containing 0.1-1 uL of inhibitor in DMSO or appropriate controls. The enzyme was incubated with inhibitor for 10 min prior to substrate addition. Reactions were initiated with 10 µL of 62.5 μM RLRGG-AMC in Buffer B. Plates were shaken vigorously for 30 s, and fluorescence from the release of AMC from peptide was monitored continuously for 15 min on a Tecan Infinite M200 Pro plate reader (λ_excitation_=360 nm; λ_emission_=460 nm). Slopes from the linear portions of each progress curve were recorded and normalized to plate-based controls. Positive control wells, representing 100% inhibition, included 10 µM GRL0617; negative control wells, representing 0% inhibition, included vehicle.

### PLpro high-throughput screening

High-throughput screening for inhibitors of PLpro was performed using the primary assay above. Test compounds (20 uM final concentration) and controls were delivered via 100 nL pin tool (V&P Scientific). The libraries included in the screen were purchased from TargetMol (Bioactive Library) and ChemDiv (a 10,000-compound SMART library subset). Each 384-well plate contains 32 positive control wells and 32 negative control wells. Average Z’ values for this assay ranged from 0.85-0.90. Compounds producing >40% inhibition of PLpro activity were selected for follow-up analysis. To eliminate compounds that interfered with AMC fluorescence and thus produced false positives, the fluorescence of 10 μM free AMC was measured in the presence of 20 µM compound in Buffer B. Inhibitors that produced a >25% decrease in AMC fluorescence signal were eliminated from further analysis. Similarly, compounds that were frequent hitters in in-house screens, or that were documented redox-cycling compounds, were eliminated from follow-up studies.

The dose responses of remaining hit compounds were tested in the primary assay over 10 compound concentrations. Percent inhibition (%I) of each data point was calculated using Equation 1:

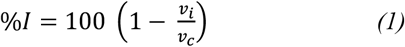

where *ν*_*i*_ is the reaction rate in the presence of inhibitor and *ν*_*c*_ is the reaction rate in the absence of inhibitor (DMSO control). Data were fit to Equation 2 using GraphPad Prism to establish an IC50:

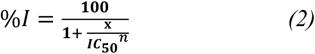

where *x* is the concentration of inhibitor and n is the Hill coefficient.

The selectivity of the most potent inhibitors was tested against the human deubiquitinating enzymes USP7 and USP14 (Boston Biochem). Assay conditions were similar to the PLpro primary assay, with the following substitutions: USP7 assays contained 4 nM USP7 and 0.5 uM Ub-AMC (Boston Biochem); USP14 assays contained 1.7 uM USP14, 4 uM Ub-AMC, and the addition of 5% glycerol to Buffer B.

PLpro activity with ISG15-AMC and Ub-AMC were assayed in a manner similar to the PLpro primary assay. PLpro and substrate concentrations were modified as follows: 80 nM PLpro was assayed with 0.5 uM Ub-AMC, and 4 nM PLpro was assayed with 0.5 uM ISG15-AMC.

### Crystallization

Crystals of SARS-CoV-2 PLpro complexed with XR compounds were grown by hanging drop vapor diffusion at 16°C. Prior to crystallization, 12 mg/mL PLpro protein was incubated with 2 mM XR824 (or XR865, XR869, XR883, XR889) for 30 min on ice. Crystals of the complexes were grown by mixing 1-2 uL of PLpro:inhibitor complex with 2 uL of reservoir solution containing 0.1 M MIB buffer, pH 7.2, 0.2 M (NH4)2SO4, and 24-28% PEG 4000 or 0.1 MIB buffer, pH 6.0-6.8, 0.2 M (NH4)2SO4, 13-16% PEG 3350, and 20% glycerol. Crystals grew overnight from the PEG 4000 conditions and were used to streak seed drops of PLpro:inhibitor equilibrating against the PEG 3350 conditions.

### Secondary binding analysis by Surface Plasmon Resonance (SPR)

The His-tagged SARS-CoV-2 PLpro enzyme was initially prepared in phosphate buffer and diluted to 50 µg/mL with 10 mM sodium acetate (pH 5.5) and immobilized on a CM5 sensor chip by standard amine-coupling with running buffer PBSP (10 mM phosphate, pH 7.4, 2.7 mM KCl, 137 mM NaCl, 0.05 % Tween-20). The CM5 sensor chip surface was first activated by 1-ethyl-3-(3-dimethylaminopropyl) carbodiimide hydrochloride (EDC)/N-hydroxy succinimide (NHS) mixture using a Biacore 8K instrument (Cytiva). SARS-CoV-2 PLpro enzyme was immobilized to flow channels 1 through 4 followed by ethanolamine blocking on the unoccupied surface area, and immobilization levels for all four channels were similar at ∼12,000 RU. Each flow channel has its own reference channel, and blank immobilization using EDC/NHS and ethanolamine was done for all reference channels. Compound solutions with a series of increasing concentrations (0.049 – 30 µM at 2.5-fold dilution) were applied to all active and reference channels in SPR binding buffer (10 mM HEPES, pH 7.4, 150 mM NaCl, and 0.05% Tween-20, 0.5 mM TCEP, and 2% DMSO) at a 30 µL/min flow rate at 25 °C. The data were double referenced with a reference channel and zero concentration (2% DMSO) responses, and reference subtracted sensorgrams were fitted with 1 to 1 Langmuir kinetic model using a Biacore Insight evaluation software, producing two rate constants (*k*_a_ and *k*_d_). The equilibrium dissociation constants (*K*_D_) were determined from two rate constants (*K*_D_ = *k*_d_/*k*_a_). For steady-state affinity fittings, response units at each concentration were measured during the equilibration phase, and the *K*_D_ values were determined by fitting the data to a single rectangular hyperbolic curve equation (3), where *y* is the response, *y*_*max*_ is the maximum response and *x* is the compound concentration.

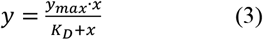

Data collection and structure refinement: The glycerol present in the crystallization solution was sufficient to cryo-protect crystals, which were flash-cooled in liquid nitrogen. Data were collected at the Life Sciences Collaborative Access Team beamlines 21-ID-D, 21-ID-G, and 21-ID-F at the Advanced Photon Source, Argonne National Laboratory. Data indexing and integration were performed using XDS.^56^ Ellipsoidal truncation and anisotropic scaling were performed by the UCLA-DOE lab’s Diffraction Anisotropy Server for the XR824 complex.^57^ Phases were determined by molecular replacement using Molrep^58^ and a SARS-CoV-2 PLpro: GRL0617 complex (PDB entry: 7JRN) as search model. Rigid body refinement followed by iterative rounds of restrained refinement and model building were performed with CCP4i modules Refmac5^59^ and Coot.^60^ The coordinates and structure factors have been deposited with PDB accession codes 7LBR (XR8-89 complex), 7LBS (XR8-24 complex), 7LLF (XR8-83 complex), 7LLZ (XR8-69 complex), and 7LOS (XR8-65 complex).

### Cell Culture and cytotoxicity

African green monkey kidney epithelial cells Vero E6 (ATCC# CRL-1586) were cultured in DMEM supplemented with 10% fetal bovine serum (Gibco), 100 units of penicillin and 100 µg/mL streptomycin (Invitrogen). Human alveolar epithelial cell line (A549) that stably express hACE2 are from BEI Resources (NR-53821). They were grown DMEM supplemented with 10% fetal bovine serum (Gibco), 100 units of penicillin and 100 µg/mL streptomycin (Invitrogen), 1% nonessential amino acids (NEAA) with 100 µg/mL Blasticidin S. HCl for selection. All cells were grown at 37 °C and 5% CO_2_. Low passage vero E6 and A549 cells (5000 cells/well) were seeded in 96-well plates and incubated at 37 °C and 5% CO_2_ for 24 hours prior to treatment. All compounds were dissolved in DMSO and final DMSO concentrations never exceeded 1%. The cytotoxicity of compounds (100 uM to 1 uM, 3-fold dilution) was examined using the CellTiter-Glo Luminescent Cell Viability Assay (Promega). Cell cytotoxicity data was normalized to DMSO control as 0% cell death.

### Antiviral activity assays

Vero E6 cells were seeded 5×10^5^ cells/well in DMEM complete into 12-well plates (1 mL/well). Cells were pretreated with 1.5 μM CP-100356 for 30 min and with both compound + 1.5 μM CP-100356 for 1-hour prior to infection. The plaque reduction assay was performed using a clinical isolate of SARS-CoV-2 (SARS-CoV-2, Isolate USA-WA1/2020) from BEI Resources. 2-fold serial dilutions of compound + CP-100356 were added to the same volume of SARS-CoV-2 (final MOI = 0.0001), the mixture was added to the monolayer of Vero E6 cells and incubated for 1 hour at 37 °C and 5% CO2. The mixture was removed, 1 mL of 1.25% (w/v) Avicel-591 in 2X MEM supplied with 4% (v/v) FBS was added onto infected cells with 10X compound + CP-100356. Plates were incubated 48 hours at 37 °C and 5% CO2. After the 48-hour incubation, the plates were fixed with 10% (v/v) formaldehyde and stained with 1% (w/v) crystal violet to visualize the plaques. All experiments were performed in a Biosafety level 3 facility. EC50 values were determined by fitting the dose-response curves with four-parameter logistic regression in Prism GraphPad (version 8.1.2). All data was normalized to virus alone. All error bars represent S.D. from three replicates.

A549-hACE2 cells were seeded 1.5×105 cells/well in DMEM complete into 24-well plates (0.5 mL/well) then incubated for 16 hours at 37 °C and 5% CO2. Cells were pretreated with compound for 1-hour prior to infection. 2-fold serial dilutions of compound added to the same volume of SARS-CoV-2 (final MOI = 0.01), the mixture was added to the monolayer cells and incubated for 1 hour at 37 °C and 5% CO2. After, the mixture was removed and replaced with 0.5 mL of infection media and incubated at 37 °C and 5% CO2. After 48 hours, supernatants were harvested and processed for RT-qPCR.

### RNA Extraction and RT-qPCR

250 µL of culture fluids were mixed with 750 µL of TRIzol™ LS Reagent (Thermo Fisher Scientific). RNA was purified following phase separation by chloroform as recommended by the manufacturer.

RNA in the aqueous phase was collected and further purified using PureLink RNA Mini Kits (Invitrogen) according to manufacturer’s protocol. Viral RNA was quantified by reverse-transcription quantitative PCR (RT-qPCR) using a 7500 Real-Time PCR System (Applied Biosystems) using TaqMan Fast Virus 1-Step Master Mix chemistry (Applied Bio-systems). SARS-CoV-2 N1 gene RNA was amplified using forward (5’-GACCCCAAAATCAGCGAAAT) and reverse (5’-TCTGGTTACTGCCAGTTGAATCTG) primers and probe (5’-FAM-ACCCCGCATTACGTTTGGTGGACC-BHQ1) designed by the United States Centers for Disease Control and Prevention (oligonucleotides produced by IDT, cat# 10006713). RNA copy numbers were determined from a standard curve produced with serial 10-fold dilutions of RNA standard material of the amplicon region from BEI Resources (NR-52358). All data was normalized to virus alone. All error bars represent S.D. from three replicates.

### Chemical synthesis and characterization

Full synthetic routes and chemical characterization are described in supporting file 2.

### Statistical analysis

GraphPad Prism 8 software package (GraphPad Software, USA) was used to perform statistical analysis. All data were presented as the mean ± SD unless otherwise noted. One-way analysis of variance (ANOVA) with appropriate post-hoc tests (3+ groups) and Student’s t-test (2 groups) were used to calculate statistical significance: \**P* < 0.05, \*\**P* < 0.01, \*\*\**P* < 0.001.

## Supporting information

Supporting information-chemistry

## ACKNOWLEDGEMENT

This study is supported by NIH grant, UL1TR002003, via UI-Centre. This research used resources of the Advanced Photon Source, a U.S. Department of Energy (DOE) Office of Science User Facility operated for the DOE Office of Science by Argonne National Laboratory under Contract No. DE-AC02-06CH11357. Use of the LS-CAT Sector 21 was supported by the Michigan Economic Development Corporation and the Michigan Technology Tri-Corridor (Grant 085P1000817). We thank Dr. Michael Flavin for his edits.

## COMPETEING INTETETS

R.X., G.T., K.R., S.Z., L.R. and L.C. are inventors of the patent application related to PLpro inhibitors.

## AUTHOR INFORMATION

These authors contributed equally: Zhengnan Shen, Kiira Ratia, and Laura Cooper.

## Author Contributions

R.X., G.T. and K.R. conceived the project. R.X., Z.S., D.K., Y.L. and S.A. synthesized the chemical library. K.R. performed structural biology and biochemical experiments. L.C. performed the antiviral assays. H.L. and Y.W. performed SPR assays. R.X., K.R., G.T. and S.Z. analyzed the data and wrote the manuscript. All authors contributed to editing the manuscript. R.X., G.T. and L. R. supervised the project.

