## Supporting data for "Potent, Novel SARS-CoV-2 PLpro Inhibitors Block Viral Replication in Monkey and Human Cell Cultures"

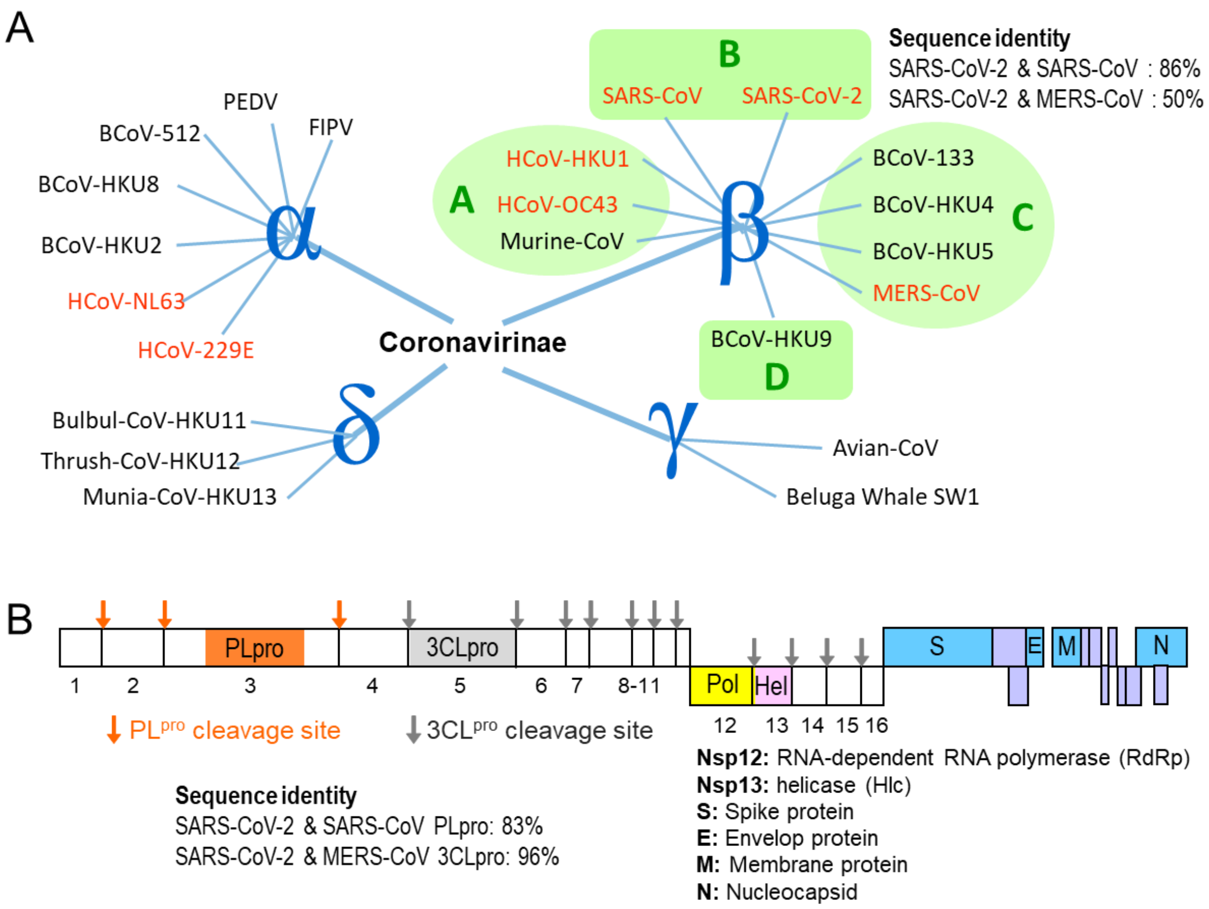


**Figure S1. A)** The phylogenetic tree of the coronavirinae family**; B)** Genetic sequence and proteins of SARS-CoV-2. PLpro cleavage sites are colored in orange; 3CLpro cleavage sites are colored in gray.

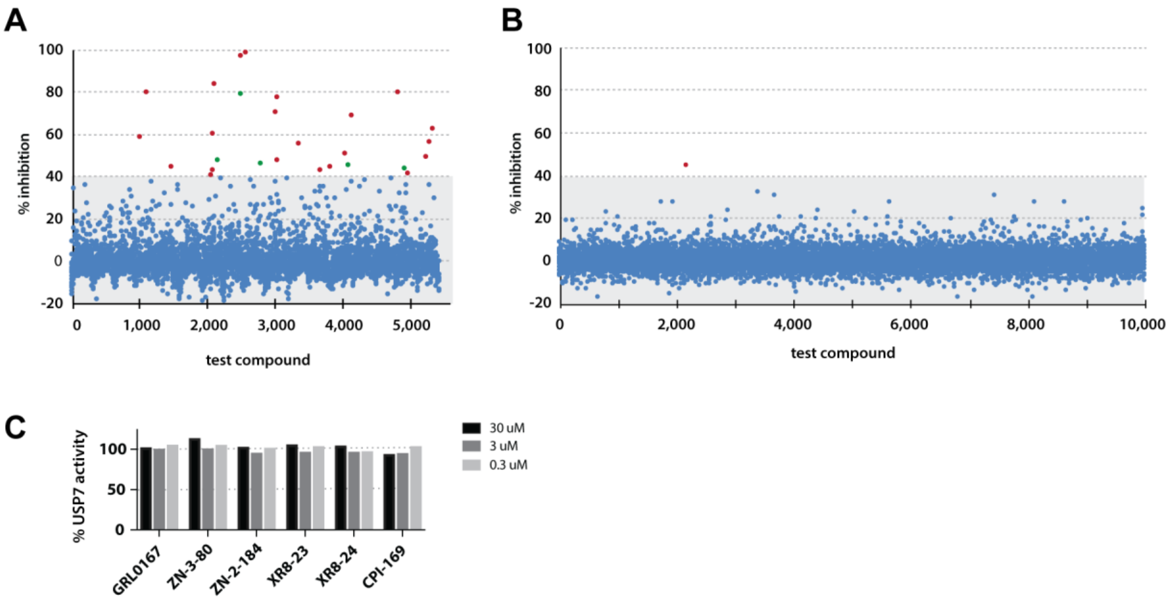


**Figure S2. High-throughput screen and counterscreen for SARS-CoV-2 PLpro inhibitors.** High-throughput screening was performed against **A)** a TargetMol Bioactives Library that contains 1,283 FDA-approved drugs and 761 drugs approved by regulatory bodies in other countries and **B)** a 10,000-compound SMART library subset from ChemDiv. Compounds producing >40% inhibition of PLpro enzymatic activity were selected for follow-up studies. **C)** Counterscreen of selected compounds against human deubiquitinating enzyme, USP7.

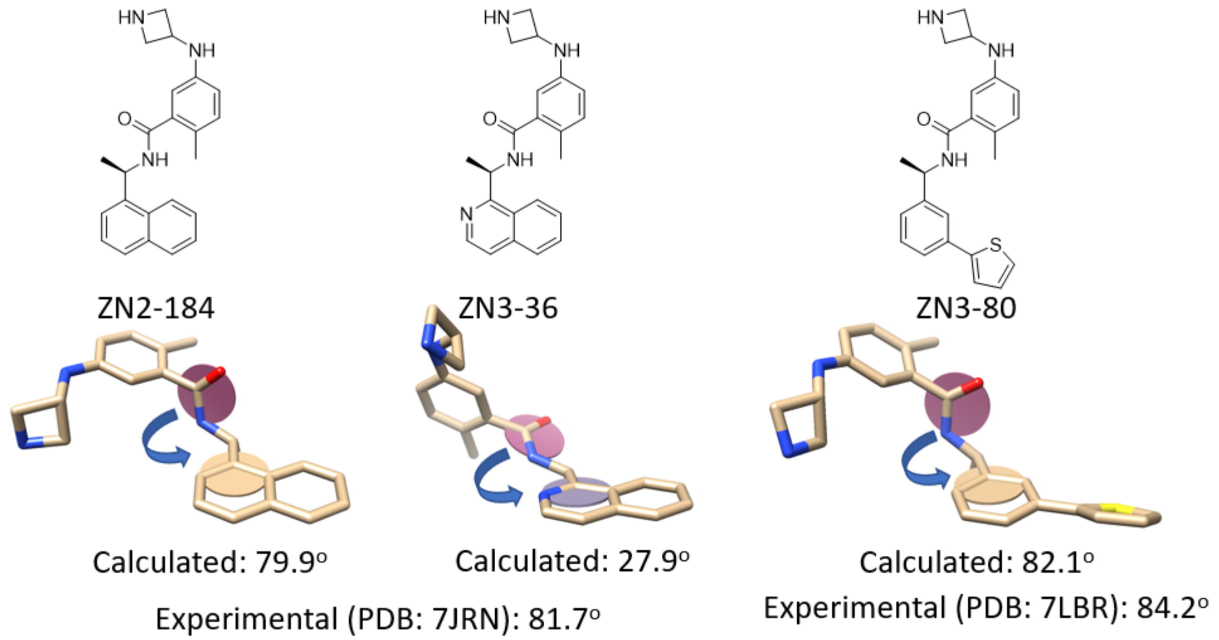


**Figure S3. Predicted angle between the amide plane and the aryl plane of GRL analogs.** Calculations use quantum mechanics (B3LYP/6-31G*) with a polarizable continuum model (PCM) as the continuum solvation method for water. The tortional angle in aniline part of the molecule was locked using experimental angles determined in PDB, 7LBR.

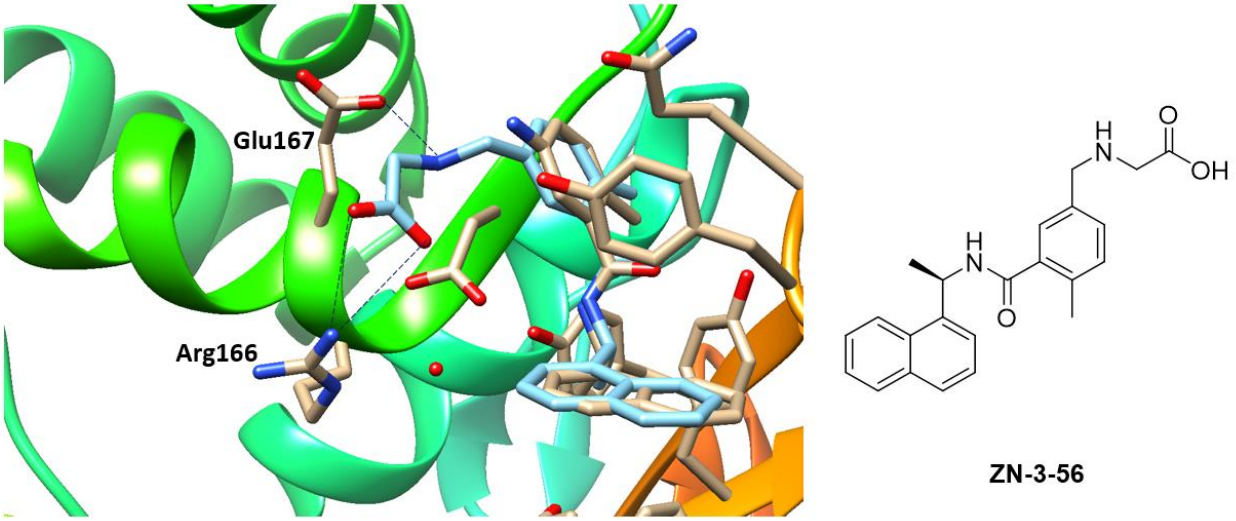


**Figure S4.** Model of ZN3-56 (light blue) bound to PLpro. The electrostatic interactions between ZN-3-56 and Arg166 and Glu167 are highlighted with dashed lines.

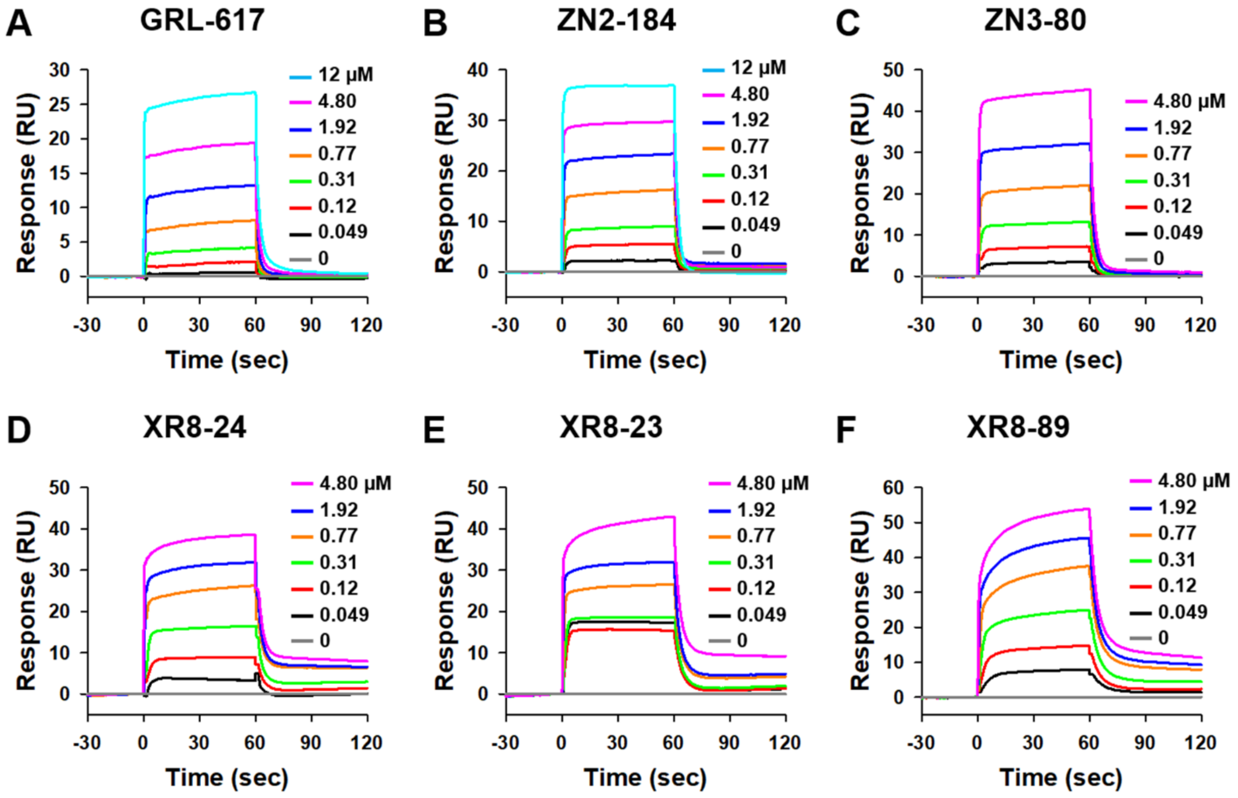


**Figure S5. SPR binding sensorgrams of GRL0617 and analogs.** The response sensorgrams were double referenced with a reference channel and zero concentration (2% DMSO) responses, and reference subtracted sensorgrams were fitted with a 1:1 Langmuir kinetic model using Biacore Insight evaluation software, producing two rate constants ($k_a$ and $k_d$). The equilibrium dissociation constants ($K_D$) were determined from the two rate constants ($K_D = k_d/k_a$).

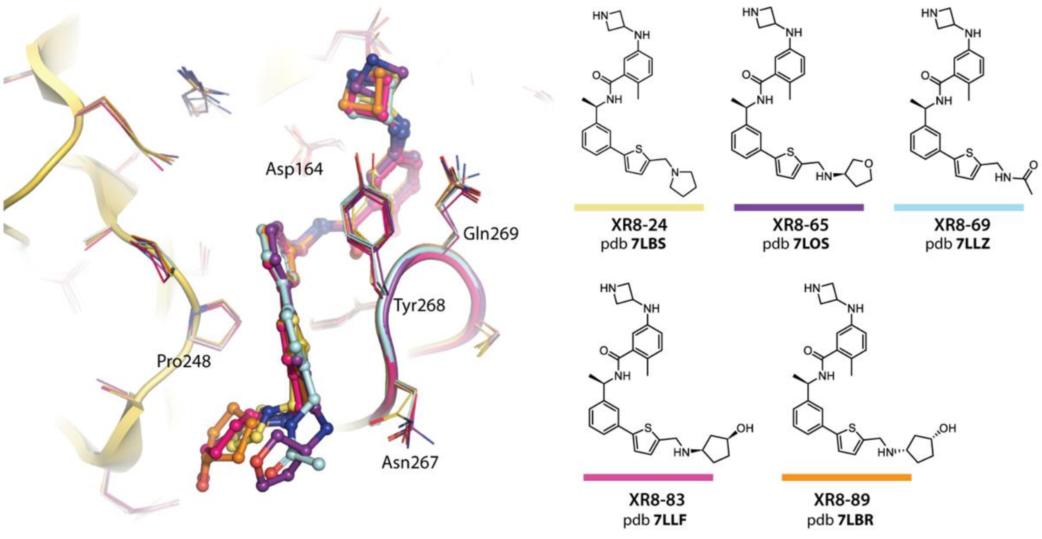


**Figure S6**. Superposition of five PLpro:XR8 inhibitor crystal structures. The chemical structures of inhibitors and their associated pdb IDs are shown at right, with colored bars corresponding to the coloring used at left.

Figure S7

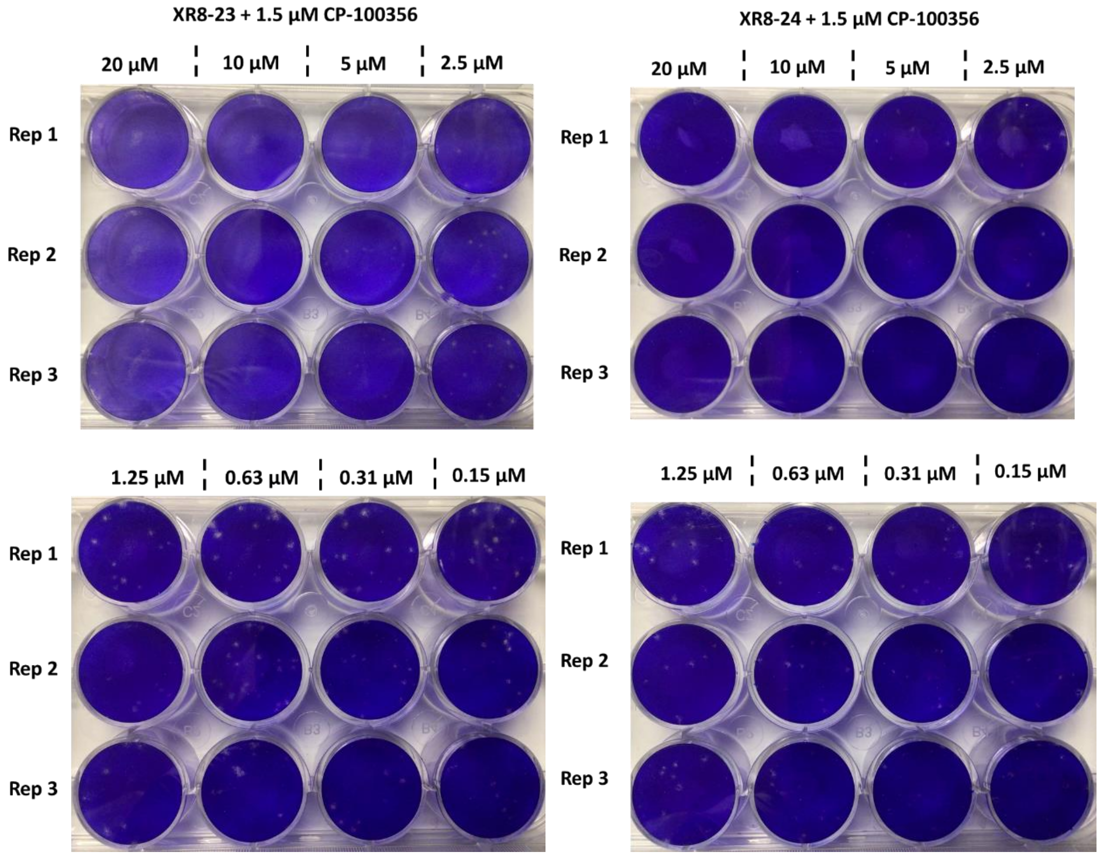


**Figure S7 Dose dependent plaque reduction of XR8-23 and XR8-24**

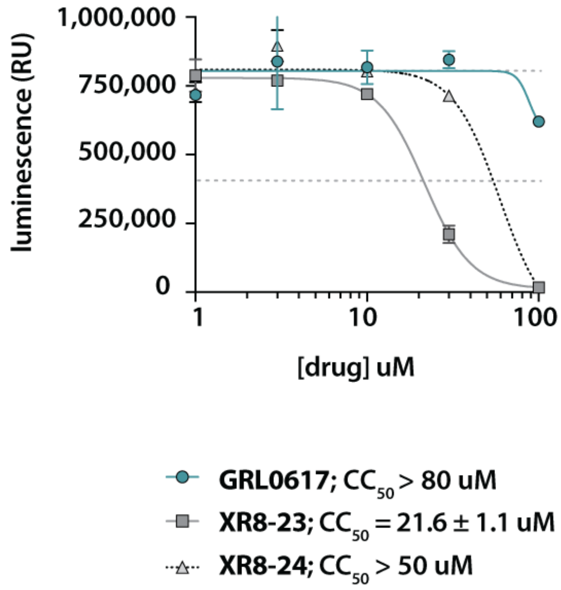


**Figure S8 Cell viability of GRL0617, XR8-23 and XR8-24 in A549-hACE2 cells**

**Table S1**.

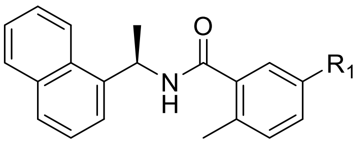

| **Compound** | $R_1$ | Enzyme inhibition $IC_{50}$ (μM) | SPR binding assays $K_D$ (μM) |
|---|---|---|---|
| **GRL0167** | 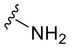 | 1.61 | 2.70 |
| **ZN-2-180** | 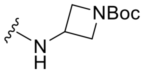 | 6.08 | 16.0 |
| **ZN-2-181** | 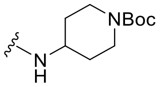 | 1.1 | 4.7 |
| **ZN-2-182** | 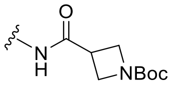 | 5.5 | 32.6 |
| **ZN-2-183** | 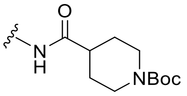 | 6.0 | 31.1 |
| **ZN-2-184** | 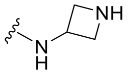 | 1.01 | 1.03 |
| **ZN-2-185** | 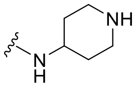 | 0.6 | 1.8 |
| **ZN-2-186** | 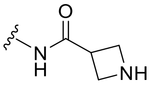 | 1.2 | 3.1 |
| **ZN-2-187** | 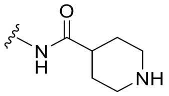 | 0.8 | 5.4 |
| **ZN-2-189** | 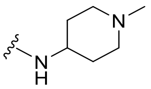 | 0.7 | 1.3 |
| **ZN-2-188-1** | 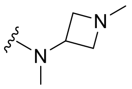 | 1.6 | 5.6 |
| **ZN-2-188-2** | 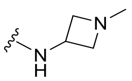 | 4.3 | 3.4 |
| **ZN-2-197** | 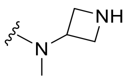 | 2.4 | 2.8 |
| **ZN-3-56** | 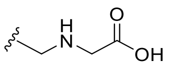 | 3.9 | 26.5 |
| **DY2-144** | 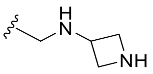 | 1.3 | 6.0 |

| | | | |
|---|---|---|---|
| **DY2-137** | 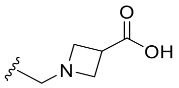 | 3.3 | 8.16 |
| **Dy2-138-2** | 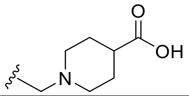 | 6.1 | NA |

**Table S2**.

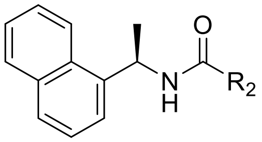


| **Compound** | R$_2$ | Enzyme inhibition IC$_{50}$ (μM) | SPR binding assays K$_D$ (μM) |
|---|---|---|---|
| **ZN-2-190** | 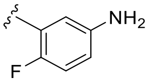 | >>100 | >1000 |
| **ZN-2-192** | 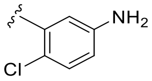 | 4.8 | 2.0 |
| **ZN-2-193** | 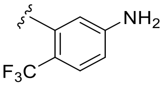 | >10 | 454 |
| **ZN-3-3** | 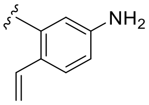 | >10 | 54.6 |
| **DY2-109** | 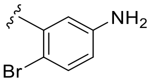 | 21 | 83.0 |
| **DY2-115** | 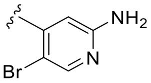 | 7 | 29.6 |
| **DY-3-8** | 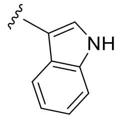 | >>100 | NA |
| **DY-3-14** | 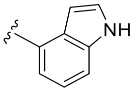 | >10 | 88.3 |
| **DY-3-15** | 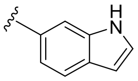 | 0.8 | 418.0 |
| **DY-3-65** | 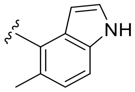 | 6.3 | 43.2 |
| **DY-3-70** | 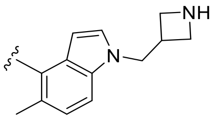 | 6.4 | 43.4 |
| **ZN-3-41** | 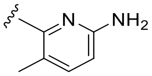 | 10-100 | >1000 |

| | | | |
|---|---|---|---|
| **ZN-3-55** | 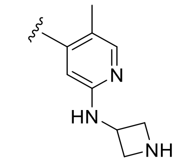 | 7.4 | 19.3 |
| **ZN-3-57** | 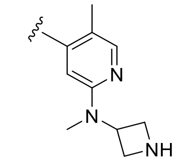 | 10-~100 | 245.5 |
| **ZN-3-66** | 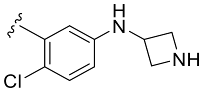 | 4.1 | 20.6 |
| **ZN-3-70** | 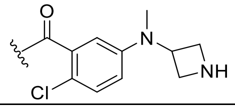 | 10.7 | 41.9 |

**Table S3**.

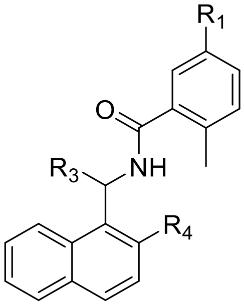


| **Compound** | R1 | R3 | R4 | Enzyme inhibition $IC_{50}$ (μM) | SPR binding assays $K_D$ (μM) |
|---|---|---|---|---|---|
| **ZN-3-13** | $NH_2$ | 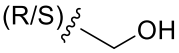 | H | ~100 | 39.1 |
| **ZN-3-19** | $NH_2$ | (R)-$CH_3$ | OH | > 100 | 184.0 |
| **DY2-97** | $NH_2$ | 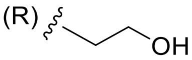 | H | ~100 | 721.5 |
| **ZN-3-61** | 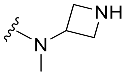 | (R)-$CH_2CH_3$ | H | >>10 | 281.5 |
| **ZN-3-32** | 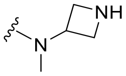 | (R)-$CH_3$ | OH | <100 | 240.2 |
| **ZN-3-33** | 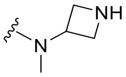 | (R)-$CH_3$ | 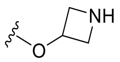 | >10 | 54.6 |
| **ZN-3-34** | 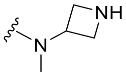 | (R)-$CH_3$ | 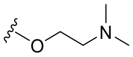 | NI | NA |
| **ZN-3-35** | 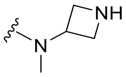 | 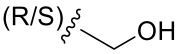 | H | >100 | 186.0 |
| **DY2-116** | 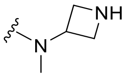 | 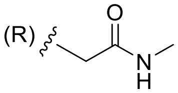 | H | NI | >1000 |
| **DY2-117** | 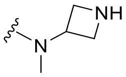 | 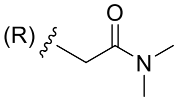 | H | NI | NA |

**Table S4**.

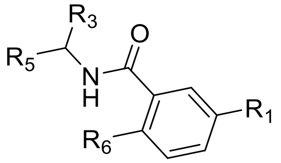


| Compound | $R_1$ | $R_3$ | $R_5$ | $R_6$ | $IC_{50}$ (μM) | $K_D$ (μM) |
|---|---|---|---|---|---|---|
| **XDY2-62** | $NH_2$ | (R)-$CH_3$ | 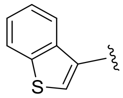 | $CH_3$ | 3.3 | 6.7 |
| **XDY2-58** | $NH_2$ | (R)-$CH_3$ | 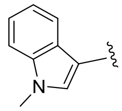 | $CH_3$ | <10 | 29.7 |
| **YF4-134** | $NO_2$ | (R/S)-$CH_3$ | 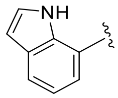 | $CH_3$ | >10 | >1000 |
| **YF4-136** | $NH_2$ | (R/S)-$CH_3$ | 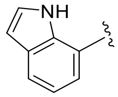 | $CH_3$ | 4.7 | 30.3 |
| **YF4-137** | $NH_2$ | (R/S)-$CH_3$ | 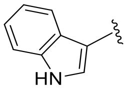 | $CH_3$ | ~100 | >1000 |
| **YF4-145** | $NH_2$ | (R/S)-$CH_3$ | 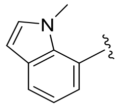 | $CH_3$ | >>100 | NA |
| **DY2-125** | 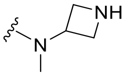 | (R)-$CH_3$ | 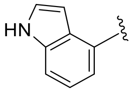 | $CH_3$ | 7 | 32.2 |
| **ZN-3-45** | 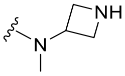 | (R/S)-$CH_3$ | 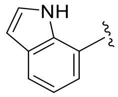 | $CH_3$ | 5.7 | 18.8 |
| **DY-3-59** | 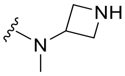 | (R)-$CH_3$ | 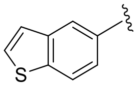 | $CH_3$ | 6.7 | 18.3 |
| **DY-3-66** | 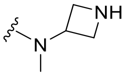 | (R)-$CH_3$ | 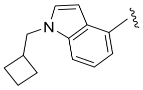 | $CH_3$ | 3.3 | 14.1 |
| **ZN-3-59** | 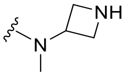 | (R)-$CH_3$ | 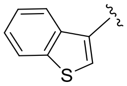 | $CH_3$ | 2.4 | 3.4 |
| **ZN-3-67** | 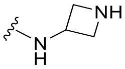 | (R)-$CH_3$ | 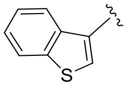 | Cl | 8.5 | 33.0 |
| **ZN-3-71** | 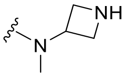 | (R)-$CH_3$ | 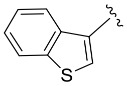 | Cl | 10.9 | 64.2 |

| | | | | | | |
|---|---|---|---|---|---|---|
| **DY2-149** | 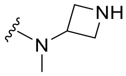 | (R)-$CH_3$ | 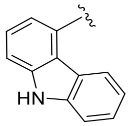 | $CH_3$ | 1.6 | 8.4 |
| **ZN-3-79** | 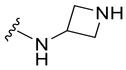 | (R)-$CH_3$ | 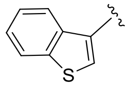 | $CH_3$ | 1.9 | 8.4 |
| **DY-2-153** | 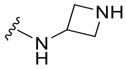 | (R)-$CH_3$ | 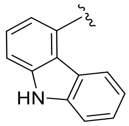 | $CH_3$ | 1.8 | 3.9 |
| **ZN-3-36** | 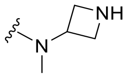 | (R)-$CH_3$ | 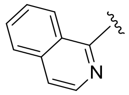 | $CH_3$ | 56 | 19.6 |
| **ZN-3-40** | 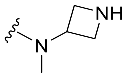 | (S)-$CH_3$ | 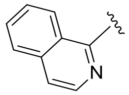 | $CH_3$ | NI | NA |
| **DY2-139** | 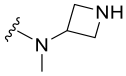 | (R)-$CH_3$ | 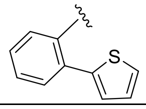 | $CH_3$ | >40 | NA |

**Table S5**.

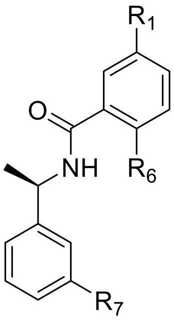


| **Compound** | $R_1$ | $R_6$ | $R_7$ | $IC_{50}$ (μM) | $K_D$ (μM) |
|---|---|---|---|---|---|
| **ZN-3-74** | 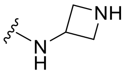 | Cl | 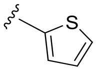 | 2.8 | NA |
| **ZN-3-80** | 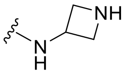 | $CH_3$ | 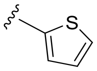 | 0.59 | 0.963 |
| **XR8-8** | 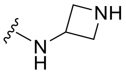 | $CH_3$ | 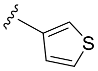 | 1.3 | 1.39 |
| **XR8-9** | 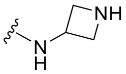 | $CH_3$ | 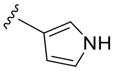 | 1.8 | 2.89 |
| **XR8-14** | 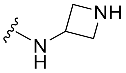 | $CH_3$ | 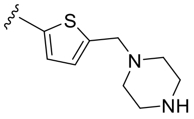 | 1.2 | 0.846 |
| **XR8-15** | 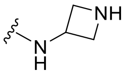 | $CH_3$ | 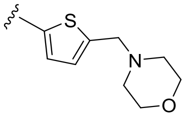 | 0.9 | 1.56 |
| **XR8-16** | 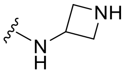 | $CH_3$ | 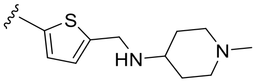 | 1.6 | 0.43 |
| **XR8-17** | 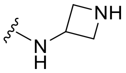 | $CH_3$ | 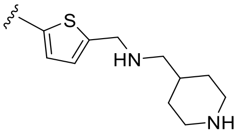 | 2.7 | 0.777 |
| **XR8-23** | 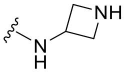 | $CH_3$ | 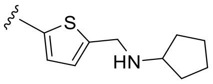 | 0.39 | 0.235 |
| **XR8-24** | 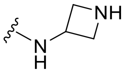 | $CH_3$ | 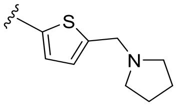 | 0.56 | 0.372 |
| **XR8-30** | 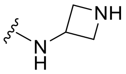 | $CH_3$ | 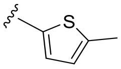 | 0.75 | 1.15 |
| **XR8-32-1** | 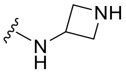 | $CH_3$ | 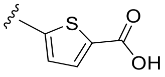 | 0.97 | 2.06 |
| **XR8-32-2** | 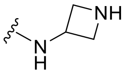 | $CH_3$ | 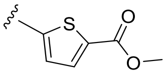 | 0.81 | 2.4 |

| | | | | | |
|---|---|---|---|---|---|
| **XR8-35** | 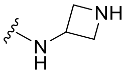 | $CH_3$ | 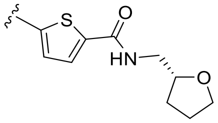 | 0.92 | 1.79 |
| **XR8-38** | 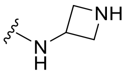 | $CH_3$ | 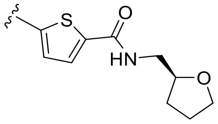 | 0.76 | 0.741 |
| **XR8-39** | 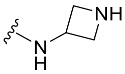 | $CH_3$ | 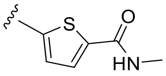 | 1.1 | 1.02 |
| **XR8-40** | 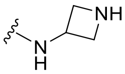 | $CH_3$ | 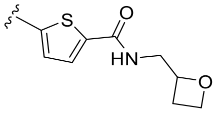 | 0.82 | 1.42 |
| **XR8-49** | 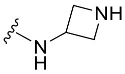 | $CH_3$ | 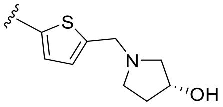 | 0.64 | 0.761 |
| **XR8-51** | 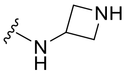 | $CH_3$ | 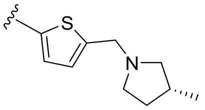 | 1.1 | 0.776 |
| **XR8-56** | 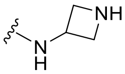 | $CH_3$ | 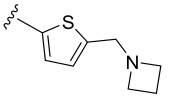 | 2.2 | 1.79 |
| **XR8-57** | 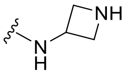 | $CH_3$ | 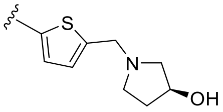 | 0.70 | 0.668 |
| **XR8-61** | 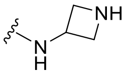 | $CH_3$ | Br | 6.5 | 3.13 |
| **XR8-65** | 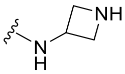 | $CH_3$ | 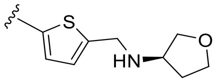 | 0.33 | 1.53 |
| **XR8-66** | 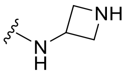 | $CH_3$ | 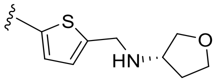 | 0.62 | 0.84 |
| **XR8-67** | 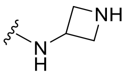 | $CH_3$ | 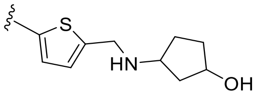 | 0.17 | 0.79 |
| **XR8-69** | 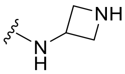 | $CH_3$ | 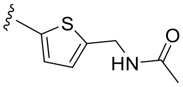 | 0.37 | NA |
| **XR8-77** | 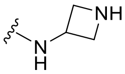 | $CH_3$ | 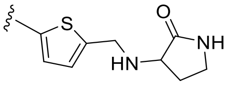 | 0.64 | 1.78 |
| **XR8-79** | 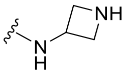 | $CH_3$ | 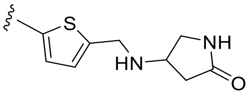 | 0.41 | 0.72 |
| **XR8-83** | 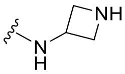 | $CH_3$ | 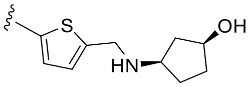 | 0.21 | 0.63 |
| **XR8-84** | 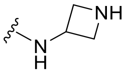 | $CH_3$ | 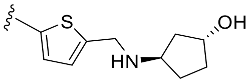 | 0.43 | 0.53 |

| | | | | | |
|---|---|---|---|---|---|
| **XR8-89** | 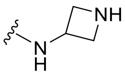 | $CH_3$ | 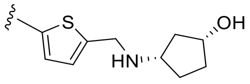 | 0.113 | 0.113 |
| **XR8-96** | 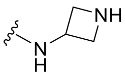 | $CH_3$ | 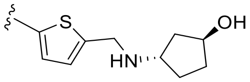 | 0.25 | 0.40 |
| **XR8-98** | $NH_2$ | $CH_3$ | 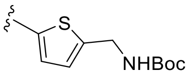 | 0.81 | 5.01 |
| **XR8-101** | $NH_2$ | $CH_3$ | 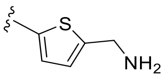 | 1.8 | 2.86 |
| **XR8-103** | $NH_2$ | $CH_3$ | 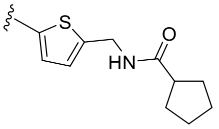 | 1.1 | 2.28 |
| **XR8-104** | NHAc | $CH_3$ | 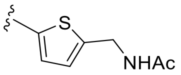 | 2.3 | 3.07 |
| **XR8-106** | $NH_2$ | $CH_3$ | 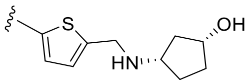 | 1.4 | 2.20 |

**Table S6. Metabolic Stability of Test Compounds in Pooled Human Liver Microsomes**

| Compound ID | Species | *In vitro* $T_{1/2}$ (min) | *In vitro* $Cl_{int}$ (μL/min/mg protein) | Scale-up $Cl_{int}$ (mL/min/kg) |
|---|---|---|---|---|
| Verapamil | Human | 13.91 | 99.66 | 124.99 |
| XR8-23 | Human | 235.50 | 5.89 | 7.38 |
| XR8-24 | Human | 144.38 | 9.60 | 12.04 |
| XR8-57 | Human | 5125.53 | 0.27 | 0.34 |
| GRL0617 | Human | 105.56 | 13.13 | 16.47 |
| ZN3-80 | Human | 245.82 | 5.64 | 7.07 |

**Table S7. Permeability Results of Test Compounds in Caco-2 Cell Line**

| Compound ID | $P_{app\ (A\text{-}B)}$ ($10^{-6}$, cm/s) | $P_{app\ (B\text{-}A)}$ ($10^{-6}$, cm/s) | Efflux Ratio | Recovery (%) AP-BL | Recovery (%) BL-AP |
|---|---|---|---|---|---|
| Propranolol | 17.92 | 18.26 | 1.02 | 58.60 | 84.44 |
| Digoxin | 0.40 | 10.45 | 26.32 | 85.20 | 92.39 |
| GRL0617 | 12.40 | 12.95 | 1.04 | 49.67 | 59.95 |
| XR8-23 | 0.16 | 1.56 | 9.94 | 68.60 | 80.29 |
| XR8-24 | 0.14 | 3.06 | 21.79 | 85.56 | 96.59 |

### Table S8 SPR data from 4 replicates

| Compounds | $k_a$1 ($M^{-1}s^{-1}$) | $k_a$2 ($M^{-1}s^{-1}$) | $k_a$3 ($M^{-1}s^{-1}$) | $k_a$4 ($M^{-1}s^{-1}$) | Ave $k_a$ ($M^{-1}s^{-1}$) | STD $k_a$ ($M^{-1}s^{-1}$) |
|---|---|---|---|---|---|---|
| **GRL0617** | 2.02E+05 | 1.73E+05 | 1.45E+05 | 1.96E+05 | 1.79E+05 | 2.61E+04 |
| **ZN2-184** | 4.26E+05 | 3.48E+05 | 2.31E+05 | 3.56E+05 | 3.40E+05 | 8.08E+04 |
| **ZN3-80** | 2.91E+05 | 3.34E+05 | 3.03E+05 | 4.77E+05 | 3.51E+05 | 8.56E+04 |
| **XR8-24** | 4.70E+05 | 2.71E+05 | 1.99E+05 | 4.17E+05 | 3.39E+05 | 1.26E+05 |
| **XR8-23** | 5.42E+05 | 5.53E+05 | 4.14E+05 | 5.79E+05 | 5.22E+05 | 7.37E+04 |
| **XR8-89** | 5.21E+05 | 4.27E+05 | | | 4.74E+05 | 6.61E+04 |

| Compounds | $k_d$1 ($s^{-1}$) | $k_d$2 ($s^{-1}$) | $k_d$3 ($s^{-1}$) | $k_d$4 ($s^{-1}$) | Ave $k_d$ ($s^{-1}$) | STD $k_d$ ($s^{-1}$) |
|---|---|---|---|---|---|---|
| **GRL0617** | 0.556 | 0.464 | 0.421 | 0.478 | 0.480 | 0.056 |
| **ZN2-184** | 0.438 | 0.407 | 0.246 | 0.305 | 0.349 | 0.089 |
| **ZN3-80** | 0.318 | 0.369 | 0.290 | 0.332 | 0.327 | 0.033 |
| **XR8-24** | 0.157 | 0.110 | 0.066 | 0.175 | 0.127 | 0.049 |
| **XR8-23** | 0.140 | 0.153 | 0.081 | 0.121 | 0.124 | 0.031 |
| **XR8-89** | 0.0529 | 0.0531 | | | 0.0530 | 0.00017 |

| Compounds | $K_D$1 (M) | $K_D$2 (M) | $K_D$3 (M) | $K_D$4 (M) | Ave $K_D$ (M) | STD $K_D$ (M) |
|---|---|---|---|---|---|---|
| **GRL0617** | 2.75E-06 | 2.69E-06 | 2.91E-06 | 2.44E-06 | 2.70E-06 | 1.94E-07 |
| **ZN2-184** | 1.03E-06 | 1.17E-06 | 1.06E-06 | 8.56E-07 | 1.03E-06 | 1.31E-07 |
| **ZN3-80** | 1.09E-06 | 1.10E-06 | 9.56E-07 | 6.97E-07 | 9.63E-07 | 1.90E-07 |
| **XR8-24** | 3.34E-07 | 4.04E-07 | 3.30E-07 | 4.20E-07 | 3.72E-07 | 4.70E-08 |
| **XR8-23** | 2.59E-07 | 2.77E-07 | 1.96E-07 | 2.09E-07 | 2.35E-07 | 3.89E-08 |
| **XR8-89** | 1.02E-07 | 1.24E-07 | | | 1.13E-07 | 1.61E-08 |
