## Supporting information-chemistry for "Potent, Novel SARS-CoV-2 PLpro Inhibitors Block Viral Replication in Monkey and Human Cell Cultures"

**General Chemical Experimental Information** Unless otherwise specified, reactions were performed under an inert atmosphere of argon and monitored by thin-layer chromatography (TLC) and/or LCMS. All reagents and solvents were purchased from commercial suppliers (Sigma-Aldrich, Fisher Scientific, Ambeed, Combi-Blocks, Enamine, and so on) and used as provided. Synthetic intermediates were purified using CombiFlash chromatography system on 230−400 mesh silica gel or Shimadzu prep-HPLC system. $^{1}$H and $^{13}$C NMR spectra were obtained using Bruker DPX-400 or AVANCE-400 spectrometer at 400 and 100 MHz, respectively. NMR chemical shifts were described in δ (ppm) using residual solvent peaks as standard (Chloroform-*d*, 7.26 ppm ($^{1}$H), 77.16 ppm ($^{13}$C); Methanol-*d*4, 3.31 ppm ($^{1}$H), 49.00 ppm ($^{13}$C); DMSO-*d*6, 2.50 ppm ($^{1}$H), 39.52 ppm ($^{13}$C); Acetone-*d*6, 2.05 ppm ($^{1}$H), 29.84 ppm ($^{13}$C) ). Data were reported in a format as follows: chemical shift, multiplicity (s = singlet, d = doublet, dd = doublet of doublet, t = triplet, q = quartet, br = broad, m = multiplet, abq = ab quartet), number of protons, and coupling constants. High resolution mass spectral data were measured in-house using a Shimadzu IT-TOF LC/MS for all final compounds. Optical rotations were measured with a Perkin-Elmer 241 polarimeter operating on the mercury lamp line (546 nm), using a 100 mm pathlength cell. All compounds submitted for biological testing were confirmed to be ≥ 95% pure by analytical HPLC. The names of the compounds were obtained using ChemDraw Ver.20.0.0.41. Synthetic methods, spectral data, and HRMS for novel compounds are described in detail below.

### Examples of synthetic routes

### Example 1

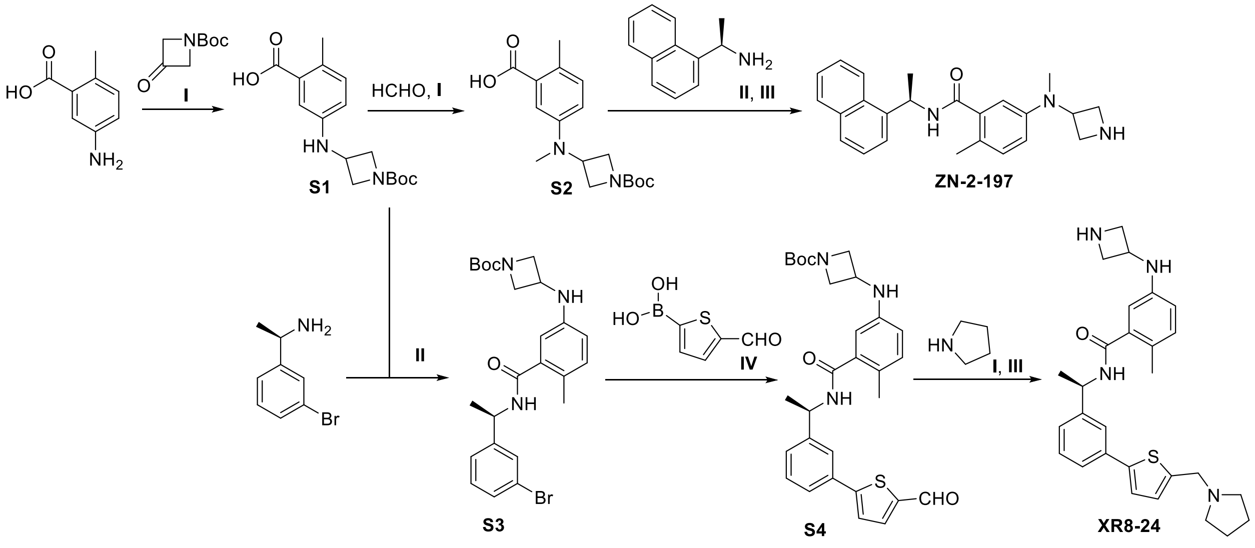


**I**. HOAc, $NaBH_3CN$, MeOH; **II**. HATU, DMAP, DMF, rt; **III**. HCl (4M in dioxane), DCM; **IV**. XPhos Pd G2, $K_3PO_4$, DMF/EtOH/$H_2O$, 95 °C

### Example 2

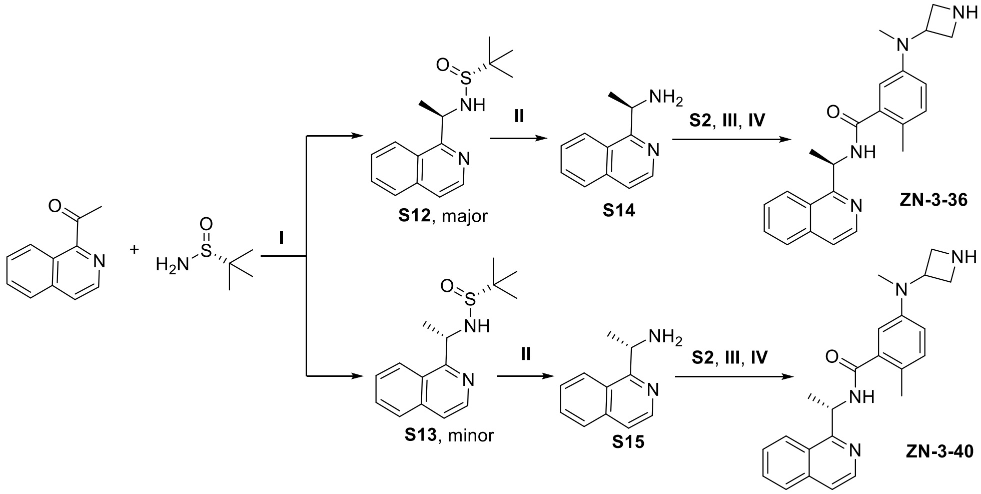


**I**.Ti(OEt)$_4$, NaBH$_4$, THF, -78 °C to rt; **II**. HCl (Con. aq.), dioxane; **III**. HATU, DMAP, DMF, rt; **IV**. HCl (4M in dioxane), DCM

### Chemical Experimental Procedures

**General Procedure for Reductive Amination**. To a solution of amine compound and ketone (or aldehyde) compound in MeOH, HOAc was added. After stirring at the indicated temperature for 2 h and cooldown, $NaBH_3CN$ was added carefully. The reaction was continued at room temperature overnight and then concentrated under vacuum. Dissolve the mixture in EA and wash with water and brine. After that, the organic layer was dried over $Na_2SO_4$, filtered, and concentrated. The residue was purified by silica gel column chromatography or Prep-HPLC to provide the amination compound.

**General Procedure for Amine Coupling**. Amine compound, acid compound, HATU, TEA (or DIPEA) and DMAP were dissolved in dry DMF or DCM and stirred at room temperature overnight. The mixture was diluted with ethyl acetate and was then washed with saturated aq. $NaHCO_3$, water, and brine, respectively. The organic layer was dried over $Na_2SO_4$, filtered, and concentrated. The residue was purified by silica gel column chromatography or Prep-HPLC to provide the desired amide.

**General Procedure for N-Boc Deprotection.** To a solution of Boc protected compound in DCM was added HCl (4M in dioxane) at 0 °C, and then warmed up to room temperature. After stirring for another 2 h, the reaction was dried under vacuum. The residue was purified by Prep-HPLC to provide the deprotected compound.

**General Procedure for Aryl Nitro Reduction**. To a solution of Aryl Nitro compound in ethanol/saturated aq. $NH_4Cl$ (4:1), Fe powder was added. The resulting solution was stirred for 2 h at 80°C.After cooling down, the mixture was extracted with ethyl acetate 3 times. The organic layer was washed with brine, dried and concentrated under vacuum. The residue was purified by silica gel column chromatography or Prep-HPLC to obtain the desired product.

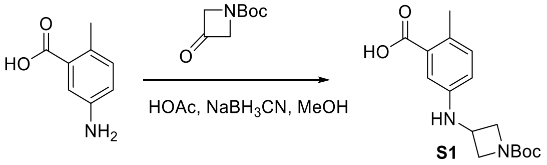

**5-((1-(tert-butoxycarbonyl)azetidin-3-yl)amino)-2-methylbenzoic acid (S1).** 5-amino-2-methylbenzoic acid (3.0 g, 19.8 mmol) and tert-butyl 3-oxoazetidine-1-carboxylate (10.1 g, 59.5 mmol) was subjected to general reductive amination procedure with MeOH (20 mL), HOAc (5 mL) at 50°C, and then $NaBH_3CN$ (6.2 g, 99.0 mmol) was added. The purification by silica gel column chromatography (Hexenes/EtOAc, 1:1) provided the amination compound (5.1 g, yield 83%) as a white solid: $^1$H NMR (400 MHz, Chloroform-*d*) δ 7.16 (d, *J* = 2.6 Hz, 1H), 7.07 (d, *J* = 8.3 Hz, 1H), 6.62 (dd, *J* = 8.3, 2.6 Hz, 1H), 4.33 – 4.27 (m, 2H), 4.24 – 4.18 (m, 1H), 3.76 – 3.70 (m, 2H), 2.49 (s, 3H), 1.43 (d, *J* = 2.1 Hz, 9H); LRMS (ESI) calcd for $C_{16}H_{23}N_2O_4$ $[M+H]^+$ 307.2, found 307.1.

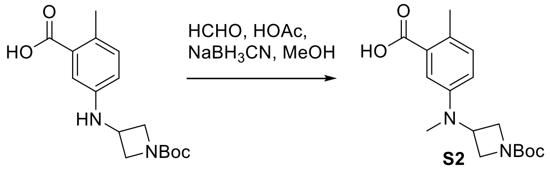


**5-((1-(tert-butoxycarbonyl)azetidin-3-yl)(methyl)amino)-2-methylbenzoic acid (S2).** 5-((1-(tert-butoxycarbonyl)azetidin-3-yl)amino)-2-methylbenzoic acid (2.3 g, 7.5 mmol) and formaldehyde (37 wt. % in $H_2O$, 2 mL) was subjected to general reductive amination procedure with MeOH (10 mL), HOAc (2 mL) at room temperature, and then $NaBH_3CN$ (2.4 g, 37.5 mmol) was added. The purification by silica gel column chromatography (Hexenes/EtOAc, 2:1) provided the amination compound (2.1 g, yield 87%) as a colorless oil: $^1$H NMR (400 MHz, Methanol-*d*$_4$) δ 7.31 (d, *J* = 2.8 Hz, 1H), 7.13 (d, *J* = 8.4 Hz, 1H), 6.88 (dd, *J* = 8.3, 2.8 Hz, 1H), 4.22 (tt, *J* = 7.0, 5.0 Hz, 1H), 4.14 (t, *J* = 8.0 Hz, 2H), 3.84 (dd, *J* = 8.9, 5.0 Hz, 2H), 2.83 (s, 3H), 2.46 (s, 3H), 1.43 (s, 9H); LRMS (ESI) calcd for $C_{17}H_{25}N_2O_4$ $[M+H]^+$ 321.2, found 321.2.

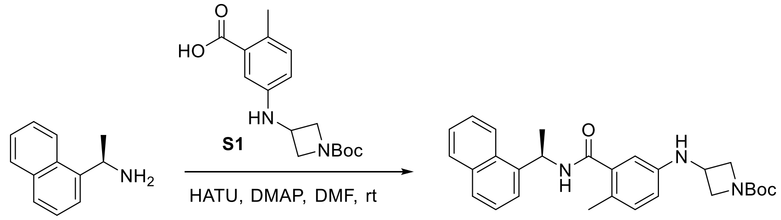


ZN-2-180

**Tert-butyl (R)-3-((4-methyl-3-((1-(naphthalen-1-yl)ethyl)carbamoyl)phenyl)amino)azetidine-1-carboxylate (ZN-2-180)**. (R)-1-(naphthalen-1-yl)ethan-1-amine (300 mg, 1.75 mmol), 5-((1-(tert-butoxycarbonyl)azetidin-3-yl)amino)-2-methylbenzoic acid (**S1**) (482 mg, 1.58 mmol), HATU (684 mg, 1.80 mmol) and DMAP (428 mg, 3.5 mmol) was subjected to general amine coupling procedure with DMF (4 mL). The purification by Prep-HPLC afforded the compound **ZN-2-180** (667 mg, yield 92%) as a white solid: $[\alpha]^{25}_{546}$ = -82.6 (c 6.7, MeOH); $^1$H NMR (400 MHz, Methanol-*d*$_4$) δ 8.69 (d, *J* = 8.2 Hz, 1H), 8.27 – 8.19 (m, 1H), 7.89 – 7.85 (m, 1H), 7.80 – 7.76 (m, 1H), 7.63 – 7.59 (m, 1H), 7.58 – 7.52 (m, 1H), 7.51 – 7.43 (m, 2H), 6.97 – 6.94 (m, 1H), 6.51 – 6.46 (m, 2H), 6.06 – 5.98 (m, 1H), 4.17 – 4.01 (m, 3H), 3.68 – 3.61 (m, 2H), 2.21 (s, 3H), 1.67 (d, *J* = 6.9 Hz, 3H), 1.43 (s, 9H). $^{13}$C NMR (100 MHz, Methanol-*d*$_4$) δ 172.40, 158.02, 146.11, 140.32, 138.66, 138.61, 135.38, 132.42, 132.33, 129.88, 128.93, 127.22, 126.68, 126.39, 125.13, 124.35, 123.77, 115.46, 112.84, 80.96, 46.35, 46.25, 44.09, 28.66, 21.37, 18.66; HRMS (ESI) calcd for $C_{28}H_{34}N_3O_3$ $[M+H]^+$ 460.2595, found 460.2590.

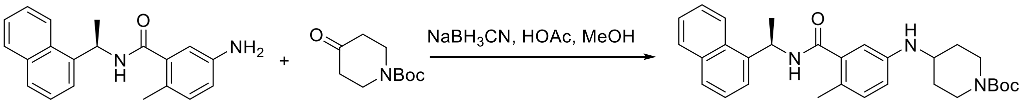


GRL0167 ZN-2-181

**Tert-butyl (R)-4-((4-methyl-3-((1-(naphthalen-1-yl)ethyl)carbamoyl)phenyl)amino)piperidine-1-carboxylate (ZN-2-181)**. (R)-5-amino-2-methyl-N-(1-(naphthalen-1-yl)ethyl)benzamide (30 mg, 0.10 mmol) and tert-butyl 4-oxopiperidine-1-carboxylate (100 mg, 0.50 mmol) was subjected to general reductive amination procedure with MeOH (1 mL), HOAc (0.2 mL) at 50 °C, and then $NaBH_3CN$ (32 mg, 0.50 mmol) was added. The purification by Prep-HPLC afforded the **ZN-2-181** (42 mg, yield 86%) as a white solid: $[\alpha]_{546}^{25}$ = -75.0 (c 0.3, MeOH); $^1H$ NMR (400 MHz, Methanol-$d_4$) δ 8.72 (d, $J$ = 8.2 Hz, 1H), 8.28 – 8.21 (m, 1H), 7.90 (dd, $J$ = 8.1, 1.4 Hz, 1H), 7.81 (d, $J$ = 8.2 Hz, 1H), 7.63 (d, $J$ = 7.1 Hz, 1H), 7.61 – 7.43 (m, 3H), 6.98 (d, $J$ = 8.3 Hz, 1H), 6.66 (dd, $J$ = 8.2, 2.5 Hz, 1H), 6.61 (d, $J$ = 2.5 Hz, 1H), 6.02 (td, $J$ = 6.8, 4.9 Hz, 1H), 4.02 – 3.93 (m, 2H), 3.41 – 3.33 (m, 1H), 2.95 – 2.83 (m, 2H), 2.21 (s, 3H), 1.96 – 1.88 (m, 2H), 1.70 (d, $J$ = 6.9 Hz, 3H), 1.46 (s, 9H), 1.33 – 1.21 (m, 2H); $^{13}C$ NMR (100 MHz, Methanol-$d_4$) δ 172.56, 156.48, 140.33, 138.70, 138.65, 135.48, 132.41, 129.91, 128.98, 127.24, 126.71, 126.41, 124.38, 123.80, 116.56, 113.79, 81.06, 51.60, 46.40, 46.31, 32.88, 28.68, 21.30, 18.57. HRMS (ESI) calcd for $C_{30}H_{38}N_3O_3$ $[M+H]^+$ 488.2908, found 488.2957.

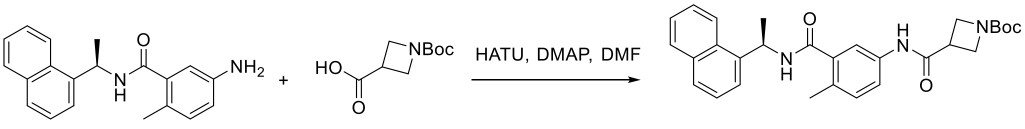


GRL0167 ZN-2-182

**Tert-butyl (R)-3-((4-methyl-3-((1-(naphthalen-1-yl)ethyl)carbamoyl)phenyl)carbamoyl)azetidine-1-carboxylate (ZN-2-182)**. **GRL0167** (30 mg, 0.10 mmol), 1-(tert-butoxycarbonyl)azetidine-3-carboxylic acid (30 mg, 0.15 mmol) , HATU (57 mg, 0.15 mmol) and DMAP (37 mg, 0.30 mmol) was subjected to general amine coupling procedure with DMF (1 mL). The purification by Prep-HPLC afforded the product **ZN-2-182** (44 mg, yield 90%) as a white solid: $[\alpha]_{546}^{25}$ = -75.0 (c 0.2, MeOH); $^1H$ NMR (400 MHz, Methanol-$d_4$) δ 8.26 (d, $J$ = 8.5 Hz, 1H), 7.93 – 7.88 (m, 1H), 7.83 – 7.78 (m, 1H), 7.66 – 7.63 (m, 1H), 7.60 – 7.46 (m, 5H), 7.18 (d, $J$ = 8.3 Hz, 1H), 6.05 (q, $J$ = 6.9 Hz, 1H), 4.08 (d, $J$ = 7.5 Hz, 4H), 3.46 (p, $J$ = 7.3 Hz, 1H), 2.30 (s, 3H), 1.71 (d, $J$ = 6.9 Hz, 3H), 1.45 (s, 9H); $^{13}C$ NMR (100 MHz, Methanol-$d_4$) δ 172.71, 171.53, 158.07, 140.30, 138.38, 137.31, 135.48, 132.61, 132.37, 132.11, 129.90, 128.98, 127.30, 126.73, 126.43, 124.29, 123.71, 122.49, 119.95, 81.27, 46.37, 34.85, 28.62, 21.40, 19.06; HRMS (ESI) calcd for $C_{30}H_{38}N_3O_3$ $[M+H]^+$ 488.2544, found 488.2534.

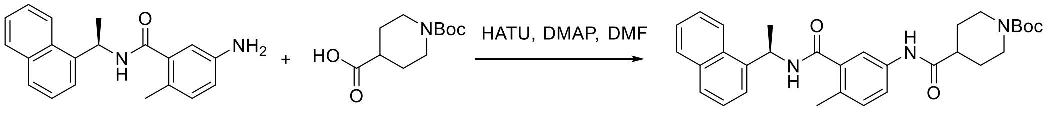


GRL0167 ZN-2-183

**Tert-butyl (R)-4-((4-methyl-3-((1-(naphthalen-1-yl)ethyl)carbamoyl)phenyl)carbamoyl)piperidine-1-carboxylate (ZN-2-183)**. **GRL0167** (30 mg, 0.10 mmol), 1-(tert-butoxycarbonyl)piperidine-4-carboxylic acid (35 mg, 0.15 mmol) , HATU (57 mg, 0.15 mmol) and DMAP (37 mg, 0.30 mmol) was subjected to general amine coupling procedure with DMF (1 mL). The purification by Prep-HPLC afforded the product **ZN-2-183** (48 mg, yield 93%) as a white solid: $[\alpha]_{546}^{25}$ = -77.0 (c 0.1, MeOH); $^1H$ NMR (400 MHz, Methanol-$d_4$)

δ 8.25 (d, *J* = 8.5 Hz, 1H), 7.92 – 7.88 (m, 1H), 7.83 – 7.79 (m, 1H), 7.65 – 7.62 (m, 1H), 7.60 – 7.44 (m, 5H), 7.17 (d, *J* = 8.3 Hz, 1H), 6.05 (q, *J* = 6.9 Hz, 1H), 4.17 – 4.09 (m, 2H), 2.89 – 2.76 (m, 2H), 2.56 – 2.47 (m, 1H), 2.30 (s, 3H), 1.84 – 1.77 (m, 2H), 1.70 (d, *J* = 6.9 Hz, 3H), 1.68 – 1.57 (m, 2H), 1.47 (s, 9H); $^{13}$C NMR (100 MHz, Methanol-*d*$_4$) δ 175.92, 171.58, 156.42, 140.30, 139.22, 138.31, 137.47, 135.47, 132.42, 132.06, 129.90, 128.98, 127.29, 126.73, 126.43, 124.29, 123.70, 122.56, 120.03, 81.17, 46.36, 44.72, 29.64, 28.68, 21.42, 19.06; HRMS (ESI) calcd for $C_{31}H_{38}N_3O_4$ [M+H]$^+$ 516.2857, found 516.2892.

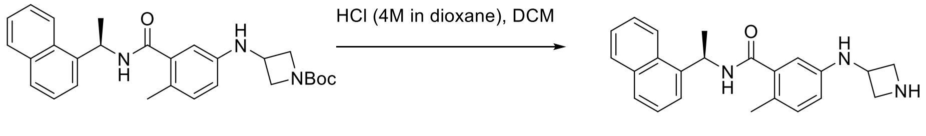


**ZN-2-180** **ZN-2-184**

**(R)-5-(azetidin-3-ylamino)-2-methyl-N-(1-(naphthalen-1-yl)ethyl)benzamide (ZN-2-184)**. General procedure for N-Boc deprotection was used with **ZN-2-180** (20 mg, 0.04 mmol) in DCM (1 mL) and HCl (4M in dioxane, 100 µL). The purification by Prep-HPLC afforded the product **ZN-2-184** (12 mg, yield 84%) as a light brown solid: $[\alpha]^{25}_{546}$ = -87.4 (c 0.8, MeOH); $^1$H NMR (400 MHz, Methanol-*d*$_4$) δ 8.53 (s, 1H), 8.28 – 8.21 (m, 1H), 7.93 – 7.88 (m, 1H), 7.84 – 7.78 (m, 1H), 7.64 – 7.61 (m, 1H), 7.60 – 7.45 (m, 3H), 7.03 – 6.99 (m, 1H), 6.57 – 6.52 (m, 1H), 6.51 – 6.48 (m, 1H), 6.07 – 6.00 (m, 1H), 4.43 (p, *J* = 7.0 Hz, 1H), 4.33 – 4.25 (m, 2H), 3.93 – 3.85 (m, 2H), 2.21 (s, 3H), 1.70 (d, *J* = 6.9 Hz, 3H); $^{13}$C NMR (100 MHz, Methanol-*d*$_4$) δ 172.26, 145.38, 140.32, 138.85, 135.47, 132.61, 132.36, 129.93, 128.98, 127.24, 126.73, 126.42, 125.82, 124.32, 123.80, 115.53, 112.83, 54.85, 46.86, 46.32, 21.35, 18.59; HRMS (ESI) calcd for $C_{23}H_{26}N_3O$ [M+H]$^+$ 360.2070, found 360.2076.

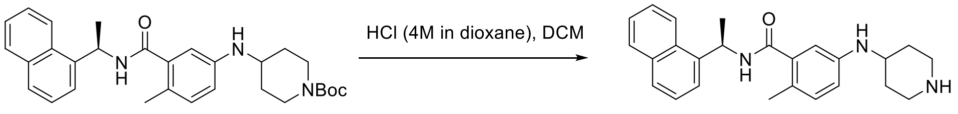


**ZN-2-181** **ZN-2-185**

**(R)-2-methyl-N-(1-(naphthalen-1-yl)ethyl)-5-(piperidin-4-ylamino)benzamide (ZN-2-185)**. General procedure for N-Boc deprotection was used with **ZN-2-181** (20 mg, 0.04 mmol) in DCM (1 mL) and HCl (4M in dioxane, 100 µL). The purification by Prep-HPLC afforded the product **ZN-2-185** (14 mg, yield 90%) as a light brown solid: $[\alpha]^{25}_{546}$ = -75.4 (c 1.2, MeOH); $^1$H NMR (400 MHz, Methanol-*d*$_4$) δ 8.48 (s, 1H), 8.27 – 8.22 (m, 1H), 7.93 – 7.87 (m, 1H), 7.84 – 7.77 (m, 1H), 7.63 (d, *J* = 7.2 Hz, 1H), 7.59 – 7.45 (m, 3H), 7.01 – 6.97 (m, 1H), 6.67 – 6.63 (m, 1H), 6.61 – 6.59 (m, 1H), 6.03 (q, *J* = 6.9 Hz, 1H), 3.51 (tt, *J* = 9.3, 3.5 Hz, 1H), 3.41 – 3.35 (m, 2H), 3.09 – 2.99 (m, 2H), 2.20 (s, 3H), 2.17 – 2.10 (m, 2H), 1.69 (d, *J* = 7.0 Hz, 3H), 1.66 – 1.54 (m, 2H); $^{13}$C NMR (100 MHz, Methanol-*d*$_4$) δ 172.58, 146.22, 140.37, 138.72, 135.46, 132.46, 132.36, 129.92, 128.96, 127.24, 126.71, 126.44, 124.62, 124.36, 123.80, 115.92, 113.18, 48.32, 46.32, 43.92, 29.92, 21.37, 18.57; HRMS (ESI) calcd for $C_{25}H_{30}N_3O$ [M+H]$^+$ 388.2383, found 388.2389.

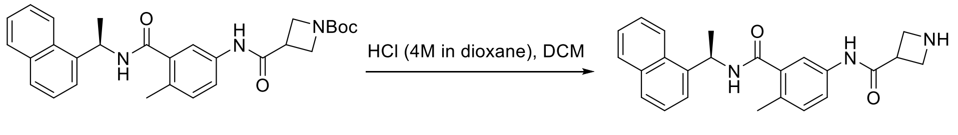


**ZN-2-182** **ZN-2-186**

**(R)-N-(4-methyl-3-((1-(naphthalen-1-yl)ethyl)carbamoyl)phenyl)azetidine-3-carboxamide (ZN-2-186)**. General procedure for N-Boc deprotection was used with **ZN-2-182** (20 mg, 0.04

mmol) in DCM (1 mL) and HCl (4M in dioxane, 100 µL). The purification by Prep-HPLC afforded the product **ZN-2-186** (14 mg, yield 88%) as a white solid: $[\alpha]^{25}_{546}$ = -94.9 (c 0.5, MeOH); $^1$H NMR (400 MHz, Methanol-*d*$_4$) δ 8.34 (s, 1H), 8.26 (d, *J* = 8.5 Hz, 1H), 7.90 (dd, *J* = 8.2, 1.5 Hz, 1H), 7.81 (d, *J* = 8.1 Hz, 1H), 7.67 – 7.42 (m, 6H), 7.19 (td, *J* = 7.6, 7.0, 4.8 Hz, 1H), 6.11 – 6.01 (m, 1H), 4.25 (t, *J* = 6.6 Hz, 2H), 3.99 – 3.73 (m, 2H), 3.43 (dd, *J* = 14.0, 9.1 Hz, 1H), 2.31 (s, 3H), 1.71 (d, *J* = 6.9 Hz, 3H); $^{13}$C NMR (100 MHz, Methanol-*d*$_4$) δ 171.49, 169.74, 140.26, 138.14, 137.02, 135.46, 132.81, 132.35, 132.10, 129.91, 129.00, 127.31, 126.74, 126.42, 124.27, 123.71, 122.51, 119.95, 47.60, 46.36, 39.88, 21.43, 19.09; HRMS (ESI) calcd for $C_{24}H_{26}N_3O_2$ $[M+H]^+$ 388.2020, found 388.2028.

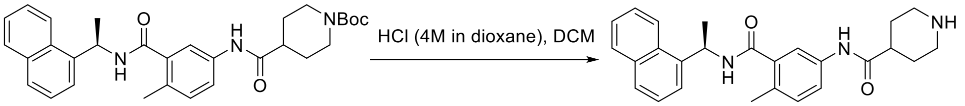


ZN-2-183 ZN-2-187

**(R)-N-(4-methyl-3-((1-(naphthalen-1-yl)ethyl)carbamoyl)phenyl)piperidine-4-carboxamide (ZN-2-187)**. General procedure for N-Boc deprotection was used with **ZN-2-183** (20 mg, 0.04 mmol) in DCM (1 mL) and HCl (4M in dioxane, 100 µL). The purification by Prep-HPLC afforded the product **ZN-2-187** (14 mg, yield 84%) as a white solid: $[\alpha]^{25}_{546}$ = -96.1 (c 0.6, MeOH); $^1$H NMR (400 MHz, Methanol-*d*$_4$) δ 8.52 (s, 1H), 8.28 – 8.22 (m, 1H), 7.93 – 7.88 (m, 1H), 7.83 – 7.78 (m, 1H), 7.66 – 7.62 (m, 1H), 7.60 – 7.45 (m, 5H), 7.20 – 7.16 (m, 1H), 6.05 (q, *J* = 6.9 Hz, 1H), 3.45 (dt, *J* = 13.0, 3.8 Hz, 2H), 3.04 (td, *J* = 12.5, 3.4 Hz, 2H), 2.67 (tt, *J* = 10.8, 4.0 Hz, 1H), 2.31 (s, 3H), 2.09 – 1.89 (m, 4H), 1.70 (d, *J* = 6.9 Hz, 3H); $^{13}$C NMR (100 MHz, Methanol-*d*$_4$) δ 174.20, 171.52, 140.28, 138.41, 137.28, 135.47, 132.55, 132.36, 132.11, 129.91, 128.99, 127.30, 126.74, 126.42, 124.28, 123.71, 122.52, 119.99, 46.35, 44.24, 41.59, 26.65, 21.41, 19.07; HRMS (ESI) calcd for $C_{26}H_{30}N_3O_2$ $[M+H]^+$ 416.2333, found 416.2340.

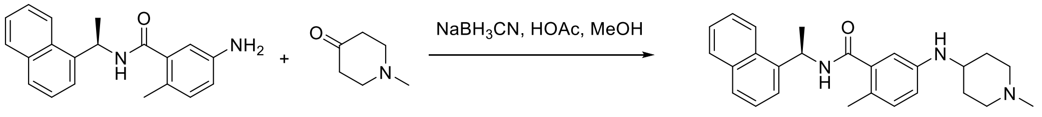


GRL0167 ZN-2-189

**(R)-2-methyl-5-((1-methylpiperidin-4-yl)amino)-N-(1-(naphthalen-1-yl)ethyl)benzamide (ZN-2-189)**. **GRL0167** (30 mg, 0.10 mmol) and 1-methylpiperidin-4-one (60 mg, 0.50 mmol) was subjected to general reductive amination procedure with MeOH (1 mL), HOAc (0.2 mL) at 50 °C, and then $NaBH_3CN$ (32 mg, 0.50 mmol) was added. The purification by Prep-HPLC afforded the product **ZN-2-189** (34 mg, yield 85%) as a white solid: $[\alpha]^{25}_{546}$ = -76.9 (c 3.0, MeOH); $^1$H NMR (400 MHz, Methanol-*d*$_4$) δ 8.54 – 8.48 (m, 1H), 8.24 (d, *J* = 8.4 Hz, 1H), 7.93 – 7.87 (m, 1H), 7.83 – 7.76 (m, 1H), 7.65 – 7.62 (m, 1H), 7.59 – 7.45 (m, 3H), 7.00 – 6.97 (m, 1H), 6.66 – 6.59 (m, 2H), 6.02 (q, *J* = 6.9 Hz, 1H), 3.51 – 3.44 (m, 1H), 3.39 – 3.33 (m, 2H), 3.07 – 2.95 (m, 2H), 2.75 (s, 3H), 2.21 (s, 3H), 2.15 – 2.06 (m, 2H), 1.71 – 1.58 (m, 5H); $^{13}$C NMR (100 MHz, Methanol-*d*$_4$) δ 169.70, 146.22, 140.48, 138.70, 135.40, 132.49, 132.28, 129.93, 128.92, 127.25, 126.73, 126.49, 124.60, 124.31, 123.75, 115.87, 113.21, 53.95, 47.46, 46.36, 43.58, 30.21, 21.48, 18.60; HRMS (ESI) calcd for $C_{26}H_{32}N_3O$ $[M+H]^+$ 402.2540, found 416.2545.

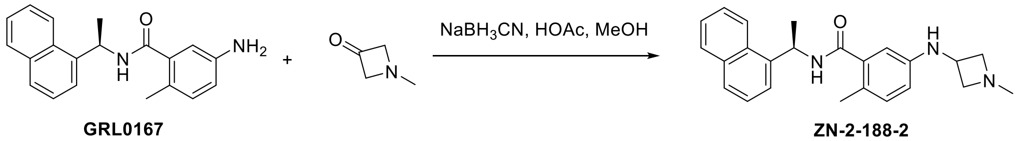


GRL0167 ZN-2-188-2

**(R)-2-methyl-5-((1-methylazetidin-3-yl)amino)-N-(1-(naphthalen-1-yl)ethyl)benzamide (ZN-2-188-2)**. **GRL0167** (100 mg, 0.31 mmol) and 1-methylazetidin-3-one (40 mg, 0.49 mmol) was subjected to general reductive amination procedure with MeOH (4 mL), HOAc (1 mL) at 50 $^{o}$C, and then $NaBH_3CN$ (98 mg, 1.55 mmol) was added. The purification by Prep-HPLC afforded the product **ZN-2-188-2** (110 mg, yield 95%) as a white solid: $[\alpha]_{546}^{25}$ = -82.1 (c 1.3, MeOH); $^{1}$H NMR (400 MHz, Methanol-$d_4$) δ 8.46 (s, 1H), 8.24 (d, $J$ = 8.4 Hz, 1H), 7.93 – 7.87 (m, 1H), 7.81 (d, $J$ = 8.2 Hz, 1H), 7.65 – 7.45 (m, 4H), 7.01 (d, $J$ = 8.2 Hz, 1H), 6.58 – 6.48 (m, 2H), 6.03 (q, $J$ = 6.9 Hz, 1H), 4.44 – 4.32 (m, 3H), 3.94 – 3.84 (m, 2H), 2.90 (s, 3H), 2.21 (s, 3H), 1.69 (d, $J$ = 6.9 Hz, 3H); $^{13}$C NMR (100 MHz, Methanol-$d_4$) δ 172.22, 145.34, 140.35, 138.84, 135.45, 132.62, 132.34, 129.93, 128.97, 127.25, 126.73, 126.44, 125.94, 124.32, 123.79, 115.57, 112.94, 63.95, 46.34, 44.05, 42.70, 21.38, 18.61; HRMS (ESI) calcd for $C_{24}H_{28}N_3O$ $[M+H]^+$ 374.2227, found 388.2230.

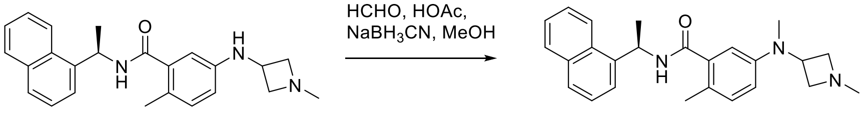


**ZN-2-188-2** **ZN-2-188-1**

**(R)-2-methyl-5-(methyl(1-methylazetidin-3-yl)amino)-N-(1-(naphthalen-1-yl)ethyl)benzamide (ZN-2-188-1)**. **ZN-2-188-2** (16 mg, 0.04 mmol) and formaldehyde (37 wt. % in $H_2O$, 200 µL) was subjected to general reductive amination procedure with MeOH (1 mL), HOAc (50 µL) at room temperature, and then $NaBH_3CN$ (14 mg, 0.21 mmol) was added. The purification by Prep-HPLC afforded the product **ZN-2-188-1** (110 mg, yield 95%) as a white solid: $[\alpha]_{546}^{25}$ = -80.9 (c 0.8, MeOH); $^{1}$H NMR (400 MHz, Methanol-$d_4$) δ 8.31 (s, 1H), 8.25 (d, $J$ = 8.5 Hz, 1H), 7.93 – 7.89 (m, 1H), 7.82 (d, $J$ = 8.2 Hz, 1H), 7.64 – 7.46 (m, 4H), 7.14 – 7.10 (m, 1H), 6.80 (dd, $J$ = 8.3, 2.7 Hz, 1H), 6.72 – 6.70 (m, 1H), 6.04 (q, $J$ = 6.9 Hz, 1H), 4.37 – 4.24 (m, 3H), 4.09 – 4.00 (m, 2H), 2.91 (s, 3H), 2.80 (s, 3H), 2.25 (s, 3H), 1.71 (d, $J$ = 6.9 Hz, 3H); $^{13}$C NMR (100 MHz, Methanol-$d_4$) δ 171.90, 148.35, 140.26, 138.88, 135.48, 132.64, 132.36, 129.97, 129.04, 128.96, 127.25, 126.75, 126.45, 124.33, 123.86, 120.07, 117.43, 61.16, 50.84, 46.44, 42.60, 37.95, 21.31, 18.70; HRMS (ESI) calcd for $C_{25}H_{30}N_3O$ $[M+H]^+$ 388.2383, found 388.2389.

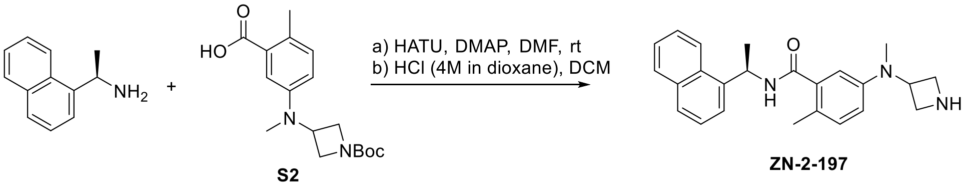


**S2** **ZN-2-197**

**(R)-5-(azetidin-3-yl(methyl)amino)-2-methyl-N-(1-(naphthalen-1-yl)ethyl)benzamide (ZN-2-197)**. (R)-1-(naphthalen-1-yl)ethan-1-amine (100 mg, 0.58 mmol), 5-((1-(tert-butoxycarbonyl)azetidin-3-yl)(methyl)amino)-2-methylbenzoic acid (**S2**) (189 mg, 0.58 mmol), HATU (221 mg, 0.58 mmol) and DMAP (213 mg, 1.74 mmol) was subjected to general amine coupling procedure with DMF (4 mL). After purification by Prep-HPLC, the product was subjected to general N-Boc deprotection procedure with HCl (4M in dioxane, 1.4 mL) and DCM (5 mL). The purification by Prep-HPLC afforded the **ZN-2-197** (150 mg, yield 69% for 2 steps) as a white solid: $[\alpha]_{546}^{25}$ = -78.9 (c 1.1, MeOH); $^{1}$H NMR (400 MHz, Methanol-$d_4$) δ 8.47 (s, 1H), 8.28 – 8.22 (m, 1H), 7.94 – 7.87 (m, 1H), 7.81 (d, $J$ = 8.2 Hz, 1H), 7.66 – 7.44 (m, 4H), 7.11 (d, $J$ = 8.3 Hz, 1H), 6.80 (dd, $J$ = 8.4, 2.7 Hz, 1H), 6.72 (d, $J$ = 2.7 Hz, 1H), 6.04 (q, $J$ = 6.9 Hz, 1H), 4.53 – 4.41 (m, 1H), 4.23 – 4.14 (m, 2H), 4.09 – 4.00 (m, 2H), 2.82 (s, 3H), 2.24 (s, 3H), 1.71 (d,

$J$ = 6.9 Hz, 3H); $^{13}$C NMR (100 MHz, Methanol-$d_4$) δ 171.98, 148.37, 140.27, 138.83, 135.48, 132.61, 132.34, 129.96, 129.02, 128.56, 127.24, 126.74, 126.44, 124.31, 123.84, 119.66, 116.96, 53.31, 51.99, 46.45, 37.03, 21.34, 18.68. HRMS (ESI) calcd for $C_{24}H_{28}N_3O$ $[M+H]^+$ 374.2227, found 374.2231.

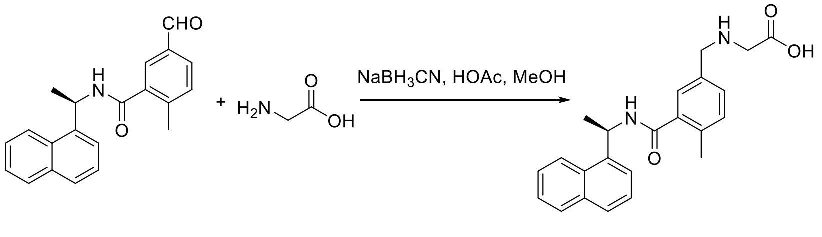


ZN-3-56

**(R)-(4-methyl-3-((1-(naphthalen-1-yl)ethyl)carbamoyl)benzyl)glycine (ZN-3-56)**. (R)-5-formyl-2-methyl-N-(1-(naphthalen-1-yl)ethyl)benzamide (30 mg, 0.10 mmol) and glycine (11 mg, 0.15 mmol) was subjected to general reductive amination procedure with MeOH (1 mL), HOAc (200 µL) at 50°C, and then $NaBH_3CN$ (19 mg, 0.30 mmol) was added. The purification by Prep-HPLC provided the amination compound **ZN-3-56** (20 mg, yield 53%) as a white solid: $[\alpha]_{546}^{25}$ = -64.4 (c 0.7, MeOH); $^1$H NMR (400 MHz, Methanol-$d_4$) δ 8.27 (d, $J$ = 8.5 Hz, 1H), 7.91 (d, $J$ = 8.1 Hz, 1H), 7.82 (d, $J$ = 8.2 Hz, 1H), 7.66 (d, $J$ = 7.3 Hz, 1H), 7.60 – 7.43 (m, 5H), 7.34 (d, $J$ = 8.1 Hz, 1H), 6.09 (q, $J$ = 6.9 Hz, 1H), 4.24 (s, 2H), 3.89 (s, 2H), 2.38 (s, 3H), 1.73 (d, $J$ = 6.9 Hz, 3H); $^{13}$C NMR (100 MHz, Methanol-$d_4$) δ 171.08, 168.15, 140.05, 138.86, 138.59, 135.47, 132.62, 132.43, 132.33, 129.94, 129.68, 129.52, 129.08, 127.34, 126.78, 126.47, 124.23, 123.85, 68.13, 47.53, 46.41, 21.38, 19.46; HRMS (ESI) calcd for $C_{23}H_{25}N_2O_3$ $[M+H]^+$ 377.1860, found 377.1857.

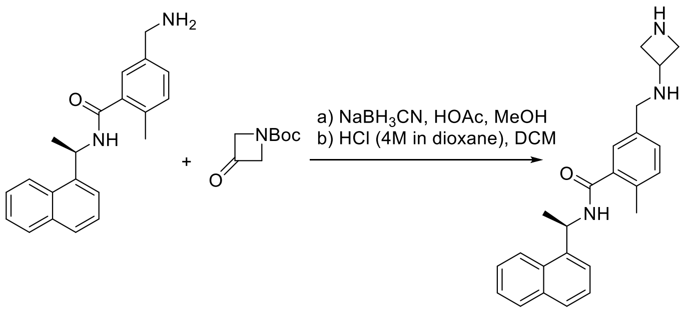


DY2-144

**(R)-5-((azetidin-3-ylamino)methyl)-2-methyl-N-(1-(naphthalen-1-yl)ethyl)benzamide (DY2-144)**. (R)-5-(aminomethyl)-2-methyl-N-(1-(naphthalen-1-yl)ethyl)benzamide[1] (30 mg, 0.09 mmol) and tert-butyl 3-oxoazetidine-1-carboxylate (31 mg, 0.18 mmol) was subjected to general reductive amination procedure with MeOH (2 mL), HOAc (500 µL) at 50 °C, and then $NaBH_3CN$ (17 mg, 0.27 mmol) was added. After purification by Prep-HPLC, the product was subjected to general N-Boc deprotection procedure with HCl (4M in dioxane, 100 µL) and DCM (2 mL). The purification by Prep-HPLC afforded the **DY2-144** (26 mg, yield 77% for 2 steps) as a white solid: $^1$H NMR (400 MHz, Methanol-$d_4$) δ 8.31 (s, 1H), 8.27 (d, $J$ = 8.6 Hz, 1H), 7.93 – 7.89 (m, 1H), 7.82 (d, $J$ = 8.2 Hz, 1H), 7.64 (d, $J$ = 7.1 Hz, 1H), 7.60 – 7.46 (m, 3H), 7.31 – 7.26 (m, 2H), 7.20 (d, $J$ = 7.7 Hz, 1H), 6.07 (q, $J$ = 6.8 Hz, 1H), 4.05 – 3.98 (m, 2H), 3.86 – 3.73 (m, 3H), 3.71 (s, 2H), 2.33 (s, 3H), 1.71 (d, $J$ = 7.0 Hz, 3H); $^{13}$C NMR (100 MHz, Methanol-$d_4$) δ 171.85, 140.23, 138.40, 135.86, 135.49, 132.38, 131.95, 130.78, 129.94, 129.04, 128.01, 127.29, 126.76, 126.45,

124.31, 123.81, 54.80, 51.33, 51.02, 46.33, 21.38, 19.32; HRMS (ESI) calcd for $C_{24}H_{28}N_3O$ $[M+H]^+$ 374.2227, found 374.2231.

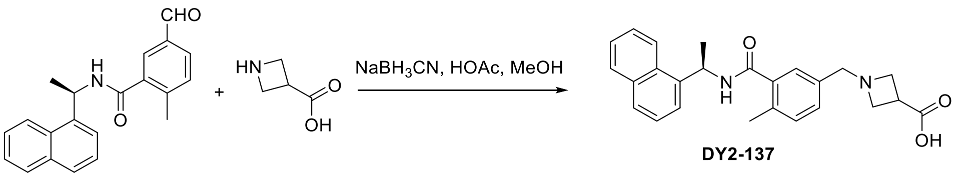


**(R)-1-(4-methyl-3-((1-(naphthalen-1-yl)ethyl)carbamoyl)benzyl)azetidine-3-carboxylic acid (DY2-137)**. (R)-5-formyl-2-methyl-N-(1-(naphthalen-1-yl)ethyl)benzamide (30 mg, 0.10 mmol) and azetidine-3-carboxylic acid (15 mg, 0.15 mmol) was subjected to general reductive amination procedure with MeOH (1 mL), HOAc (200 µL) at 50°C, and then $NaBH_3CN$ (19 mg, 0.30 mmol) was added. The purification by Prep-HPLC provided the amination compound **DY2-137** (18 mg, yield 45%) as a white solid: $^1H$ NMR (400 MHz, Acetone-$d_6$) δ 8.48 (d, $J$ = 8.1 Hz, 1H), 8.28 (d, $J$ = 8.5 Hz, 1H), 7.94 – 7.87 (m, 1H), 7.78 (d, $J$ = 8.1 Hz, 1H), 7.71 – 7.63 (m, 2H), 7.57 – 7.42 (m, 3H), 7.32 (d, $J$ = 7.7 Hz, 1H), 7.15 (d, $J$ = 7.6 Hz, 1H), 6.05 (p, $J$ = 7.0 Hz, 1H), 4.12 (s, 2H), 4.00 – 3.88 (m, 4H), 3.22 – 3.15 (m, 1H), 2.35 (s, 3H), 1.63 (d, $J$ = 6.9 Hz, 3H); $^{13}C$ NMR (100 MHz, Acetone-$d_6$) δ 176.13, 164.02, 141.08, 138.09, 137.92, 134.86, 131.98, 131.78, 131.60, 130.18, 129.62, 128.37, 126.97, 126.44, 126.35, 124.34, 123.52, 67.58, 58.97, 57.17, 45.49, 35.10, 21.78, 19.85; HRMS (ESI) calcd for $C_{25}H_{27}N_2O_3$ $[M+H]^+$ 403.2016, found 403.2019.

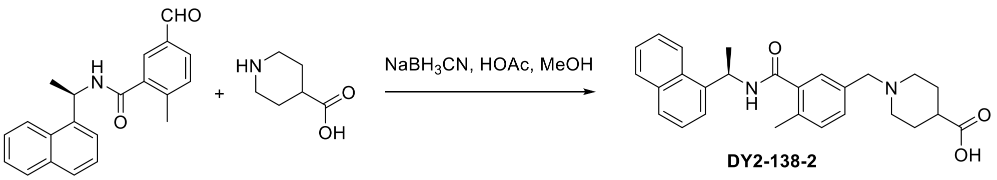


**(R)-1-(4-methyl-3-((1-(naphthalen-1-yl)ethyl)carbamoyl)benzyl)piperidine-4-carboxylic acid (DY2-138-2)**. (R)-5-formyl-2-methyl-N-(1-(naphthalen-1-yl)ethyl)benzamide (30 mg, 0.10 mmol) and piperidine-4-carboxylic acid (19 mg, 0.15 mmol) was subjected to general reductive amination procedure with MeOH (1 mL), HOAc (200 µL) at 50°C, and then $NaBH_3CN$ (19 mg, 0.30 mmol) was added. The purification by Prep-HPLC provided the amination compound **DY2-138-2** (19 mg, yield 44%) as a white solid: $^1H$ NMR (400 MHz, Acetone-$d_6$) δ 8.33 (d, $J$ = 8.5 Hz, 1H), 8.00 (d, $J$ = 8.1 Hz, 1H), 7.93 (dd, $J$ = 8.2, 1.5 Hz, 1H), 7.83 (d, $J$ = 8.2 Hz, 1H), 7.71 (d, $J$ = 7.1 Hz, 1H), 7.63 – 7.57 (m, 1H), 7.55 – 7.44 (m, 3H), 7.28 (dd, $J$ = 7.8, 1.8 Hz, 1H), 7.16 (d, $J$ = 7.7 Hz, 1H), 6.11 (p, $J$ = 7.2 Hz, 1H), 2.90 (d, $J$ = 11.2 Hz, 2H), 2.37 (s, 3H), 2.34 – 2.18 (m, 3H), 2.07 – 2.07 (m, 2H), 1.92 – 1.84 (m, 2H), 1.79 – 1.67 (m, 5H); $^{13}C$ NMR (100 MHz, Acetone-$d_6$) δ 176.16, 163.15, 140.80, 138.02, 137.99, 136.02, 134.92, 132.10, 131.34, 131.16, 129.61, 128.94, 128.48, 126.99, 126.48, 126.29, 124.40, 123.61, 62.35, 53.03, 45.43, 45.33, 40.81, 28.43, 21.57, 19.63; HRMS (ESI) calcd for $C_{27}H_{31}N_2O_3$ $[M+H]^+$ 431.2329, found 431.2333.

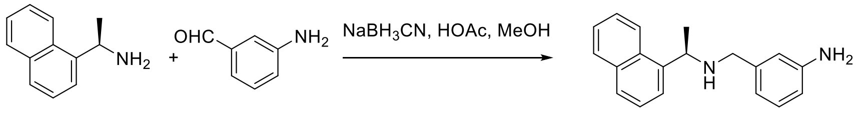


**(R)-3-(((1-(naphthalen-1-yl)ethyl)amino)methyl)aniline (XDY2-64)**. (R)-1-(naphthalen-1-yl)ethan-1-amine (172 mg, 1.00 mmol) and 3-aminobenzaldehyde (61 mg, 0.50 mmol) was subjected to general reductive amination procedure with MeOH (5 mL), HOAc (500 µL) at 50°C, and then $NaBH_3CN$ (189 mg, 3.00 mmol) was added. The purification by Prep-HPLC provided the amination compound **XDY2-64** (134 mg, yield 48%) as a white solid: $^1$H NMR (400 MHz, Methanol-$d_4$) δ 8.40 – 8.31 (m, 2H), 8.14 – 8.06 (m, 1H), 8.04 – 7.95 (m, 2H), 7.83 – 7.77 (m, 1H), 7.68 – 7.57 (m, 3H), 6.93 (d, *J* = 7.9 Hz, 1H), 6.74 – 6.63 (m, 2H), 5.44 (q, *J* = 6.8 Hz, 1H), 3.98 (dd, *J* = 91.8, 13.1 Hz, 2H), 1.83 (d, *J* = 6.7 Hz, 3H); $^{13}$C NMR (100 MHz, Methanol-$d_4$) δ 147.53, 135.57, 134.31, 132.86, 132.38, 131.22, 131.11, 130.42, 128.56, 127.60, 127.33, 126.63, 125.07, 122.97, 118.09, 118.00, 53.94, 20.21, 17.95; HRMS (ESI) calcd for $C_{19}H_{21}N_2$ $[M+H]^+$ 277.1699, found 277.1709.

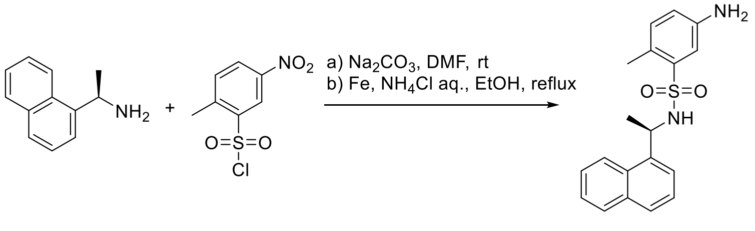


**DY-3-63**

**(R)-5-amino-2-methyl-N-(1-(naphthalen-1-yl)ethyl)benzenesulfonamide (DY-3-63)**. To a solution of (R)-1-(naphthalen-1-yl)ethan-1-amine (86 mg, 0.50 mmol) and 2-methyl-5-nitrobenzenesulfonyl chloride (141 mg, 0.60 mmol) in DMF (5 mL), $Na_2CO_3$ (159 mg, 1.5 mmol) was added. The reaction was stirred at room temperature overnight and then was diluted with ethyl acetate, washed with saturated aq. $NaHCO_3$, water, and brine, respectively. The organic layer was dried over $Na_2SO_4$, filtered, and concentrated. After purification by silica gel column chromatography, the product was applied to the general Aryl Nitro reduction procedure with ethanol/ saturated aq. $NH_4Cl$ (10 mL/2 mL) and Iron Powder (140 mg, 2.50 mmol). The purification by Prep-HPLC afforded the desired product **DY-3-63** (124 mg, yield 73%) as a light yellow solid: $^1$H NMR (400 MHz, DMSO-$d_6$) δ 8.14 (d, *J* = 8.2 Hz, 1H), 7.99 – 7.88 (m, 2H), 7.79 (d, *J* = 8.1 Hz, 1H), 7.63 – 7.59 (m, 1H), 7.53 – 7.43 (m, 3H), 7.15 (d, *J* = 2.5 Hz, 1H), 6.92 (dd, *J* = 8.1, 0.8 Hz, 1H), 6.61 (dd, *J* = 8.0, 2.5 Hz, 1H), 5.05 (p, *J* = 7.0 Hz, 1H), 2.35 (s, 3H), 1.35 (d, *J* = 6.9 Hz, 3H); $^{13}$C NMR (100 MHz, DMSO-$d_6$) δ 146.75, 139.46, 139.16, 133.19, 132.75, 129.56, 128.62, 127.30, 126.09, 125.43, 125.30, 123.36, 122.59, 122.10, 117.25, 113.96, 48.54, 22.73, 18.75; HRMS (ESI) calcd for $C_{19}H_{21}N_2O_2S$ $[M+H]^+$ 341.1318, found 341.1311.

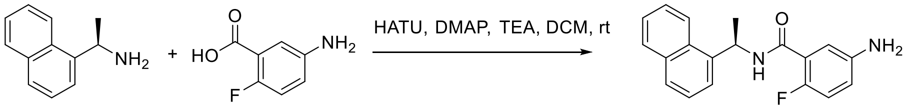


**ZN-2-190**

**(R)-5-amino-2-fluoro-N-(1-(naphthalen-1-yl)ethyl)benzamide (ZN-2-190)**. (R)-1-(naphthalen-1-yl)ethan-1-amine (221 mg, 1.29 mmol), 5-amino-2-fluorobenzoic acid (100 mg, 0.64 mmol), HATU (269 mg, 0.71 mmol), TEA (200 µL), and DMAP (8 mg, 0.06 mmol) was subjected to general amine coupling procedure with DCM (4 mL). The purification by Prep-HPLC afforded the product **ZN-2-190** (167 mg, 85%) as a white solid: $[\alpha]^{25}_{546}$ = -72.2 (c 13.3, MeOH); $^1$H NMR (400 MHz, Methanol-$d_4$) δ 8.19 – 8.15 (m, 1H), 7.82 (dd, *J* = 8.2, 1.5 Hz, 1H), 7.73 (d, *J* = 8.2 Hz, 1H), 7.60 (d, *J* = 7.1 Hz, 1H), 7.53 – 7.38 (m, 3H), 7.00 (dd, *J* = 6.0, 2.9 Hz, 1H), 6.89 – 6.83 (m, 1H), 6.76 – 6.71 (m, 1H), 6.03 (q, *J* = 6.8 Hz, 1H), 1.63 (d, *J* = 6.9 Hz, 3H); $^{13}$C NMR (100 MHz, Methanol-$d_4$) δ 166.07, 154.04 (d, *J* = 237.9 Hz), 145.48, 140.27, 135.21, 132.01, 129.81, 128.83, 127.20, 126.61, 126.41, 124.21 (d, *J* = 14.8 Hz), 124.06, 123.49, 119.76 (d, *J* = 8.0 Hz), 117.36 (d,

$J$ = 24.2 Hz), 116.43, 46.71, 38.78, 21.70; HRMS (ESI) calcd for $C_{19}H_{18}FN_2O$ $[M+H]^+$ 309.1398, found 309.1404.

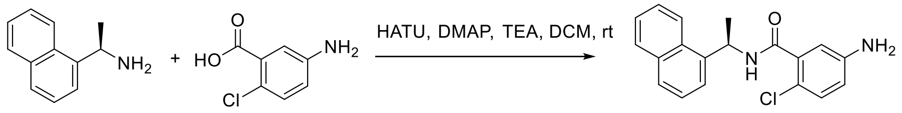


**ZN-2-192**

**(R)-5-amino-2-chloro-N-(1-(naphthalen-1-yl)ethyl)benzamide (ZN-2-192)**. (R)-1-(naphthalen-1-yl)ethan-1-amine (200 mg, 1.16 mmol), 5-amino-2-chlorobenzoic acid (100 mg, 0.58 mmol), HATU (243 mg, 0.64 mmol), TEA (200 µL), and DMAP (8 mg, 0.06 mmol) was subjected to general amine coupling procedure with DCM (4 mL). The purification by Prep-HPLC afforded the **ZN-2-192** (147 mg, 78%) as a white solid: $[\alpha]_{546}^{25}$ = -75.6 (c 12.8, MeOH); $^1$H NMR (400 MHz, Methanol-$d_4$) δ 8.22 (d, $J$ = 8.5 Hz, 1H), 7.88 (dd, $J$ = 8.1, 1.5 Hz, 1H), 7.79 (d, $J$ = 8.2 Hz, 1H), 7.64 (d, $J$ = 7.2 Hz, 1H), 7.60 – 7.42 (m, 3H), 7.11 (dd, $J$ = 8.0, 1.0 Hz, 1H), 6.72 (d, $J$ = 8.0 Hz, 2H), 6.01 (q, $J$ = 6.9 Hz, 1H), 1.68 (d, $J$ = 6.9 Hz, 3H); $^{13}$C NMR (100 MHz, Methanol-$d_4$) δ 169.43, 147.36, 140.07, 137.96, 135.39, 132.28, 131.34, 129.83, 128.92, 127.24, 126.66, 126.39, 124.32, 123.77, 119.88, 118.69, 116.13, 46.58, 21.41; HRMS (ESI) calcd for $C_{19}H_{18}ClN_2O$ $[M+H]^+$ 325.1102, found 325.1101.

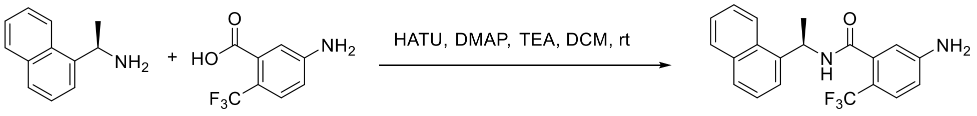


**ZN-2-193**

**(R)-5-amino-N-(1-(naphthalen-1-yl)ethyl)-2-(trifluoromethyl)benzamide (ZN-2-193)**. (R)-1-(naphthalen-1-yl)ethan-1-amine (200 mg, 1.16 mmol), 5-amino-2-(trifluoromethyl)benzoic acid (119 mg, 0.58 mmol), HATU (243 mg, 0.64 mmol), TEA (200 µL), and DMAP (8 mg, 0.06 mmol) was subjected to general amine coupling procedure with DCM (4 mL). The purification by Prep-HPLC afforded the **ZN-2-193** (154 mg, 74%) as a white solid: $[\alpha]_{546}^{25}$ = -66.2 (c 3.4, MeOH); $^1$H NMR (400 MHz, Methanol-$d_4$) δ 8.25 – 8.19 (m, 1H), 7.88 (dd, $J$ = 8.2, 1.4 Hz, 1H), 7.79 (d, $J$ = 8.2 Hz, 1H), 7.64 – 7.61 (m, 1H), 7.59 – 7.54 (m, 1H), 7.52 – 7.44 (m, 2H), 7.37 (d, $J$ = 8.6 Hz, 1H), 6.70 (dd, $J$ = 8.8, 2.4 Hz, 1H), 6.64 – 6.62 (m, 1H), 6.02 (q, $J$ = 6.9 Hz, 1H), 1.66 (d, $J$ = 6.9 Hz, 3H); $^{13}$C NMR (100 MHz, Methanol-$d_4$) δ 170.40, 152.88, 140.00, 138.39, 135.38, 132.32, 129.82, 128.92, 128.70 (q, $J$ = 4.8 Hz), 127.29, 126.67, 126.39, 126.10 (q, $J$ = 270.7 Hz), 124.23, 123.74, 114.94, 114.17, 46.39, 21.22; HRMS (ESI) calcd for $C_{20}H_{18}F_3N_2O$ $[M+H]^+$ 359.1366, found 359.1371.

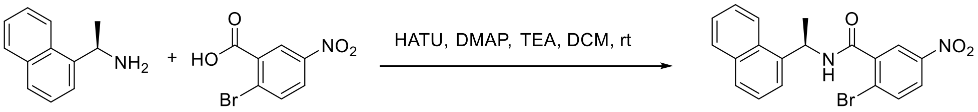


**S5**

**(R)-2-bromo-N-(1-(naphthalen-1-yl)ethyl)-5-nitrobenzamide**. (R)-1-(naphthalen-1-yl)ethan-1-amine (200 mg, 1.16 mmol), 2-bromo-5-nitrobenzoic acid (285 mg, 1.16 mmol), HATU (456 mg, 1.20 mmol), TEA (500 µL), and DMAP (13 mg, 0.1 mmol) was subjected to general amine coupling procedure with DCM (10 mL). The purification by silica gel column chromatography (Hexenes/EtOAc, 3:1) provided the amination compound (431 mg, yield 93%) as a white solid: $^1$H NMR (400 MHz, Chloroform-$d$) δ 8.31 (d, $J$ = 2.7 Hz, 1H), 8.22 (d, $J$ = 8.5 Hz, 1H), 8.07 (dd, $J$ = 8.7, 2.7 Hz, 1H), 7.91 – 7.88 (m, 1H), 7.84 (d, $J$ = 8.2 Hz, 1H), 7.75 (d, $J$ = 8.7 Hz, 1H), 7.64 –

7.46 (m, 5H), 6.22 – 6.11 (m, 1H), 1.86 (d, *J* = 6.4 Hz, 3H); LRMS (ESI) calcd for $C_{20}H_{16}BrN_2O_3$ $[M+H]^+$ 399.0, found 399.1.

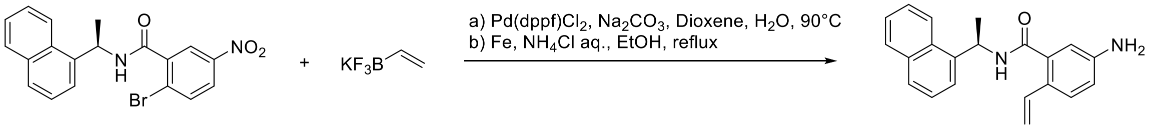


S5 ZN-3-3

**(R)-5-amino-N-(1-(naphthalen-1-yl)ethyl)-2-vinylbenzamide (ZN-3-3)**. A flask fitted with a rubber septum was charged with (R)-2-bromo-N-(1-(naphthalen-1-yl)ethyl)-5-nitrobenzamide (100 mg, 0.25 mmol), Potassium vinyltrifluoroborate (67 mg, 0.50 mmol), Pd(dppf)$Cl_2$ (25 mg, 0.03 mmol), $Na_2CO_3$ (63 mg, 0.75 mmol), Dioxene/$H_2O$ (4 mL/ 0.4 mL) and then purged with argon. The mixture was stirred at 90 °C overnight. The reaction mixture was then cooled to room temperature, diluted with ethyl acetate (50 mL), filtered through celite and concentrated in vacuo. After purification by Prep-HPLC, the product was applied to the general Aryl Nitro reduction procedure with ethanol/ saturated aq. $NH_4Cl$ (4 mL/1 mL) and Iron Powder (42 mg, 0.75 mmol). The purification by Prep-HPLC afforded the desired product **ZN-3-3** (53 mg, yield 67%) as a light brown solid: $[\alpha]^{25}_{546}$ = -69.8 (c 0.2, MeOH); $^1H$ NMR (400 MHz, Methanol-$d_4$) δ 8.25 (d, *J* = 8.5 Hz, 1H), 7.93 – 7.87 (m, 1H), 7.81 (d, *J* = 8.2 Hz, 1H), 7.64 – 7.44 (m, 4H), 7.40 (d, *J* = 8.4 Hz, 1H), 6.80 – 6.70 (m, 2H), 6.64 – 6.62 (m, 1H), 6.03 (q, *J* = 6.9 Hz, 1H), 5.50 (dd, *J* = 17.5, 1.3 Hz, 1H), 4.96 (dd, *J* = 11.1, 1.3 Hz, 1H), 1.68 (d, *J* = 7.0 Hz, 3H); $^{13}C$ NMR (100 MHz, Methanol-$d_4$) δ 172.09, 148.86, 140.39, 138.15, 135.46, 135.07, 132.34, 129.88, 128.93, 127.39, 127.27, 126.70, 126.41, 125.93, 124.30, 123.73, 117.56, 114.00, 111.86, 46.39, 21.43; HRMS (ESI) calcd for $C_{21}H_{21}N_2O$ $[M+H]^+$ 317.1648, found 374.2231.

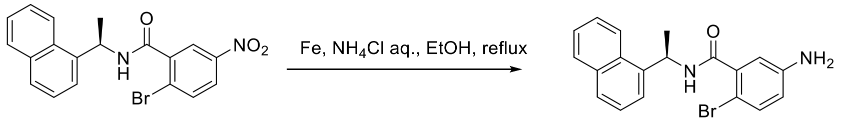


S5 DY2-109

**(R)-5-amino-2-bromo-N-(1-(naphthalen-1-yl)ethyl)benzamide** (**DY2-109**). (R)-2-bromo-N-(1-(naphthalen-1-yl)ethyl)-5-nitrobenzamide (100 mg, 0.25 mmol) was applied to the general Aryl Nitro reduction procedure with ethanol/ saturated aq. $NH_4Cl$ (4 mL/1 mL) and Iron Powder (42 mg, 0.75 mmol). The purification by Prep-HPLC afforded the desired product **DY2-109** (84 mg, yield 91%) as a light brown solid: Light yellow solid (yield 82%): $^1H$ NMR (400 MHz, Methanol-$d_4$) δ 8.24 (d, *J* = 8.5 Hz, 1H), 7.93 – 7.86 (m, 1H), 7.80 (d, *J* = 8.2 Hz, 1H), 7.65 (d, *J* = 7.2 Hz, 1H), 7.59 – 7.44 (m, 3H), 7.24 (d, *J* = 8.6 Hz, 1H), 6.67 (d, *J* = 2.8 Hz, 1H), 6.62 (dd, *J* = 8.5, 2.8 Hz, 1H), 6.01 (q, *J* = 6.9 Hz, 1H), 1.69 (d, *J* = 6.9 Hz, 3H); LRMS (ESI) calcd for $C_{19}H_{18}BrN_2O$ $[M+H]^+$ 369.1, found 369.0.

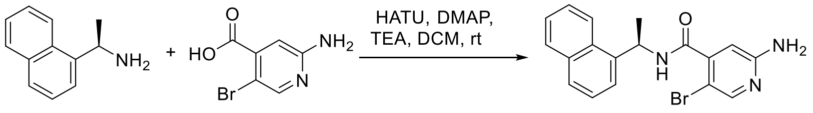


DY2-115

**(R)-2-amino-5-bromo-N-(1-(naphthalen-1-yl)ethyl)isonicotinamide (DY2-115)**. (R)-1-(naphthalen-1-yl)ethan-1-amine (41 mg, 0.24 mmol), 2-amino-5-bromoisonicotinic acid (25 mg, 0.12 mmol), HATU (46 mg, 0.12 mmol), TEA (100 µL), and DMAP (3 mg, 0.02 mmol) was subjected to general amine coupling procedure with DCM (2 mL). The purification by Prep-HPLC afforded the product **DY2-115 (34)** (31 mg, yield 71%) as a brown solid: $[\alpha]^{25}_{546}$ = -37.4 (c 0.5, MeOH); $^{1}H$ NMR (400 MHz, Methanol-$d_4$) δ 8.22 (d, *J* = 8.7 Hz, 1H), 8.00 (s, 1H), 7.91 – 7.88 (m, 1H), 7.81 (d, *J* = 8.3 Hz, 1H), 7.64 (d, *J* = 7.1 Hz, 1H), 7.60 – 7.45 (m, 3H), 6.52 (s, 1H), 6.01 (q, *J* = 6.9 Hz, 1H), 1.70 (d, *J* = 7.0 Hz, 3H); $^{13}C$ NMR (100 MHz, Methanol-$d_4$) δ 168.04, 160.22, 150.46, 148.32, 139.70, 135.44, 132.31, 129.88, 129.11, 127.37, 126.76, 126.39, 124.31, 123.89, 109.01, 103.71, 46.50, 21.28; HRMS (ESI) calcd for $C_{18}H_{17}BrN_3O$ $[M+H]^+$ 370.0550, found 370.0556.

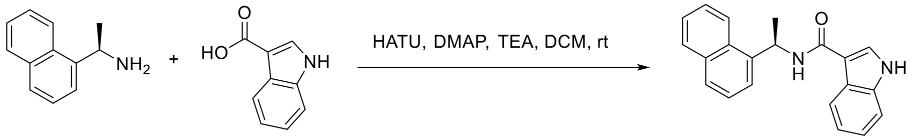


**DY-3-8**

**(R)-N-(1-(naphthalen-1-yl)ethyl)-1H-indole-3-carboxamide (DY-3-8)**. (R)-1-(naphthalen-1-yl)ethan-1-amine (233 mg, 1.36 mmol), 1H-indole-3-carboxylic acid (200 mg, 1.24 mmol), HATU (708 mg, 1.86 mmol) and DIPEA (0.6 mL, 3.72 mmol) was subjected to general amine coupling procedure with DMF (5 mL). The purification by Prep-HPLC afforded the product **DY-3-8** (334 mg, yield 86%) as a white solid: $[\alpha]^{25}_{546}$ = -14.8 (c 0.5, MeOH); $^{1}H$ NMR (400 MHz, Chloroform-*d*) δ 9.06 (s, 1H), 8.25 (d, *J* = 8.4 Hz, 1H), 7.90 – 7.84 (m, 2H), 7.80 (d, *J* = 8.2 Hz, 1H), 7.69 (d, *J* = 2.9 Hz, 1H), 7.63 (d, *J* = 6.4 Hz, 1H), 7.55 – 7.44 (m, 3H), 7.39 (dd, *J* = 6.8, 1.6 Hz, 1H), 7.20 (pd, *J* = 7.1, 1.4 Hz, 2H), 6.23 – 6.16 (m, 1H), 1.81 (d, *J* = 6.5 Hz, 2H); $^{13}C$ NMR (100 MHz, Chloroform-*d*) δ 164.35, 138.82, 136.40, 134.00, 131.21, 128.72, 128.29, 127.92, 126.56, 125.83, 125.25, 124.75, 123.64, 122.78, 122.59, 121.49, 119.94, 111.93, 44.75, 21.22; HRMS (ESI) calcd for $C_{21}H_{19}N_2O$ $[M+H]^+$ 315.1492, found 315.1498.

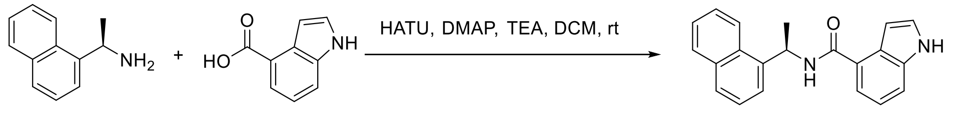


**DY-3-14**

**(R)-N-(1-(naphthalen-1-yl)ethyl)-1H-indole-4-carboxamide (DY-3-14)**. (R)-1-(naphthalen-1-yl)ethan-1-amine (94 mg, 0.55 mmol), 1H-indole-4-carboxylic acid (80 mg, 0.50 mmol), HATU (283 mg, 0.74 mmol) and DIPEA (0.26 mL, 1.5 mmol) was subjected to general amine coupling procedure with DMF (5 mL). The purification by Prep-HPLC afforded the **DY-3-14** (117 mg, yield 75%) as a white solid: $^{1}H$ NMR (400 MHz, DMSO-$d_6$) δ 11.24 (s, 1H), 8.76 (d, *J* = 7.9 Hz, 1H), 8.29 (d, *J* = 8.4 Hz, 1H), 7.95 (dd, *J* = 8.0, 1.6 Hz, 1H), 7.83 (d, *J* = 8.1 Hz, 1H), 7.69 (d, *J* = 6.7 Hz, 1H), 7.59 (ddd, *J* = 8.5, 6.8, 1.6 Hz, 1H), 7.56 – 7.47 (m, 4H), 7.39 (t, *J* = 2.8 Hz, 1H), 7.16 – 7.11 (m, 1H), 6.80 – 6.69 (m, 1H), 6.03 (p, *J* = 7.1 Hz, 1H), 1.63 (d, *J* = 6.9 Hz, 3H); $^{13}C$ NMR (100 MHz, DMSO-$d_6$) δ 167.02, 140.83, 136.46, 133.36, 130.45, 128.61, 127.06, 126.66, 126.31, 126.06, 126.00, 125.49, 123.25, 122.55, 120.02, 118.53, 114.02, 101.74, 44.43, 39.52, 21.61; HRMS (ESI) calcd for $C_{21}H_{19}N_2O$ $[M+H]^+$ 315.1492, found 315.1491.

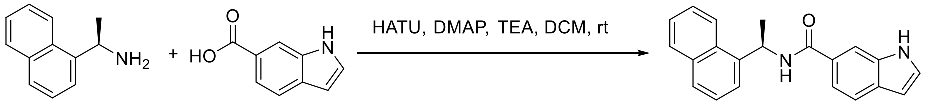


**DY-3-15**

**(R)-N-(1-(naphthalen-1-yl)ethyl)-1H-indole-6-carboxamide (DY-3-15)**. (R)-1-(naphthalen-1-yl)ethan-1-amine (58 mg, 0.34 mmol), 1H-indole-6-carboxylic acid (50 mg, 0.31 mmol), HATU (171 mg, 0.46 mmol) and DIPEA (0.15 mL, 0.93 mmol) was subjected to general amine coupling procedure with DMF (5 mL). The purification by Prep-HPLC afforded the **DY-3-15** (69 mg, yield 71%) as a white solid: $[\alpha]^{25}_{546}$ = -5.0 (c 2.5, MeOH); $^{1}$H NMR (400 MHz, DMSO-$d_6$) δ 11.34 (s, 1H), 8.86 (d, *J* = 7.8 Hz, 1H), 8.25 (d, *J* = 8.4 Hz, 1H), 8.00 (s, 1H), 7.99 – 7.92 (m, 1H), 6.49 (td, *J* = 2.0, 0.9 Hz, 1H), 6.01 (p, *J* = 7.1 Hz, 1H), 1.64 (d, *J* = 6.9 Hz, 3H); $^{13}$C NMR (100 MHz, DMSO-$d_6$) δ 166.92, 141.31, 135.66, 133.84, 130.95, 130.25, 129.09, 128.39, 127.82, 127.56, 126.57, 125.98, 125.93, 123.69, 123.04, 119.75, 118.69, 111.88, 101.66, 45.12, 22.06; HRMS (ESI) calcd for $C_{21}H_{19}N_2O$ $[M+H]^+$ 315.1492, found 315.1498.

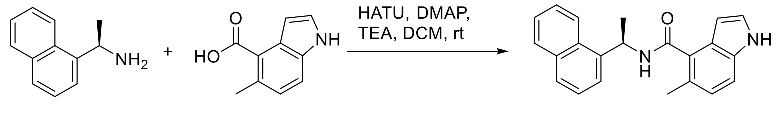


DY-3-65

**(R)-5-methyl-N-(1-(naphthalen-1-yl)ethyl)-1H-indole-4-carboxamide (DY-3-65)**. (R)-1-(naphthalen-1-yl)ethan-1-amine (150 mg, 0.88 mmol), 5-methyl-1H-indole-4-carboxylic acid (128 mg, 0.73 mmol), HATU (333 mg, 0.88 mmol), and TEA (400 µL) was subjected to general amine coupling procedure with DMF (5 mL). The purification by Prep-HPLC afforded the **DY-3-65** (209 mg, yield 87%) as a brown solid: $^{1}$H NMR (400 MHz, Methanol-$d_4$) *δ* 7.91 (d, J = 8.1 Hz, 1H), 7.82 (d, J = 8.2 Hz, 1H), 7.66 (d, J = 7.1 Hz, 1H), 7.60 (ddd, J = 8.4, 6.9, 1.3 Hz, 1H), 7.56 – 7.45 (m, 2H), 7.30 (d, J = 8.3 Hz, 1H), 7.16 (d, J = 3.2 Hz, 1H), 6.96 (d, J = 8.3 Hz, 1H), 6.28 – 6.24 (m, 1H), 6.20 (q, J = 6.9 Hz, 1H), 2.38 (s, 3H), 1.76 (d, J = 6.9 Hz, 3H). $^{13}$C NMR (100 MHz, Methanol-$d_4$) *δ* 140.30, 135.47, 132.46, 129.87, 128.97, 127.25, 126.73, 126.40, 125.99, 124.79, 124.60, 123.97, 113.18, 100.86, 49.00, 46.16, 21.46, 19.13; HRMS (ESI) calcd for $C_{22}H_{22}N_2O$ $[M+H]^+$ 329.1648, found 329.1649.

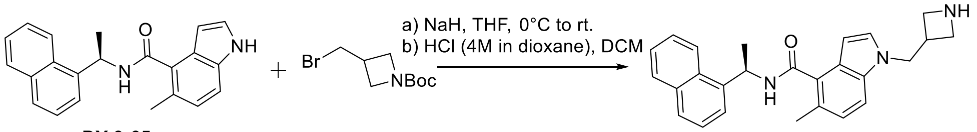


DY-3-65 DY-3-70

**(R)-1-(azetidin-3-ylmethyl)-5-methyl-N-(1-(naphthalen-1-yl)ethyl)-1H-indole-4-carboxamide (DY-3-70)**. To a solution of **DY-3-65** (40 mg, 0.12 mmol) in dry THF (5 mL), sodium hydride (12 mg 0.25 mmol) was added at 0 °C. After 15 minutes, the mixture was added tert-butyl 3-(bromomethyl)azetidine-1-carboxylate (30 mg, 0.16 mmol) and stirred at 25 °C for another 16 hours. Quench the reaction with methanol and remove the solvent. The residue was purified by preparative HPLC system to obtain the desired product.

The product from previous step was subjected to general N-Boc deprotection procedure with HCl (4M in dioxane, 200 µL) and DCM (4 mL). The purification by Prep-HPLC afforded the product **DY-3-70** (39 mg, yield 82% for 2 steps) as a white solid: $[\alpha]^{25}_{546}$ = -63.9 (c 1.1, MeOH); $^{1}$H NMR (400 MHz, Methanol-$d_4$) δ 8.53 (s, 1H), 7.33 (d, J = 8.1 Hz, 1H), 7.25 (d, J = 3.2 Hz, 1H), 7.18 – 7.07 (m, 3H), 6.79 (dd, J = 8.3, 2.7 Hz, 1H), 6.73 (d, J = 2.6 Hz, 1H), 6.66 (d, J = 3.7 Hz, 1H), 5.62 (q, J = 7.0 Hz, 1H), 4.46 (p, J = 7.2 Hz, 1H), 4.19 (dd, J = 12.3, 7.4 Hz, 4H), 4.05 (dd, J = 10.3, 7.4 Hz, 2H), 2.83 (s, 3H), 2.23 (s, 3H), 2.06 – 1.96 (m, 2H), 1.93 – 1.79 (m, 4H), 1.65 (d, J = 7.0 Hz, 3H). $^{13}$C NMR (100 MHz, Methanol-$d_4$) δ 148.36, 139.08, 137.99, 136.30, 132.56, 129.04, 128.62, 127.88, 122.25, 119.64, 117.05, 116.55, 111.39, 109.89, 99.99, 53.41, 52.28, 52.02, 49.00, 37.58, 37.12, 27.10, 21.24, 18.98, 18.65; HRMS (ESI) calcd for $C_{26}H_{28}N_3O$ $[M+H]^+$ 398.2227, found 398.2230.

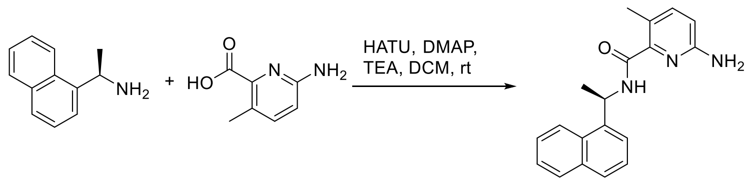


ZN-3-41

**(R)-6-amino-3-methyl-N-(1-(naphthalen-1-yl)ethyl)picolinamide (ZN-3-41)**. (R)-1-(naphthalen-1-yl)ethan-1-amine (55 mg, 0.32 mmol), 6-amino-3-methylpicolinic acid (25 mg, 0.16 mmol), HATU (61 mg, 0.16 mmol), TEA (100 µL), and DMAP (3 mg, 0.02 mmol) was subjected to general amine coupling procedure with DCM (2 mL). The purification by Prep-HPLC afforded the product **ZN-3-41** (43 mg, yield 87%) as a brown solid: $[\alpha]^{25}_{546}$ = -116.1 (c 0.6, MeOH); $^{1}$H NMR (400 MHz, Methanol-$d_4$) δ 8.21 – 8.16 (m, 1H), 7.89 – 7.86 (m, 1H), 7.80 – 7.77 (m, 1H), 7.65 – 7.62 (m, 1H), 7.56 – 7.43 (m, 4H), 7.31 (d, *J* = 8.4 Hz, 1H), 6.60 (d, *J* = 8.3 Hz, 1H), 5.97 (q, *J* = 6.9 Hz, 1H), 2.39 (s, 3H), 1.70 (d, *J* = 6.9 Hz, 3H); $^{13}$C NMR (100 MHz, Methanol-$d_4$) δ 168.28, 162.59, 158.18, 146.77, 143.24, 140.42, 135.45, 132.25, 129.86, 128.93, 127.25, 126.68, 126.45, 124.20, 123.54, 112.72, 46.02, 21.73, 18.70; HRMS (ESI) calcd for $C_{19}H_{20}N_3O$ $[M+H]^+$ 306.1601, found 306.1602.

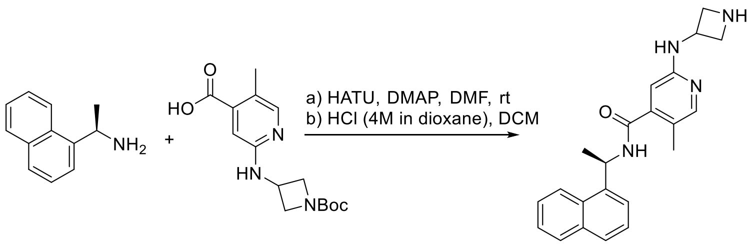


ZN-3-55

**(R)-2-(azetidin-3-ylamino)-5-methyl-N-(1-(naphthalen-1-yl)ethyl)isonicotinamide (ZN-3-55)**. (R)-1-(naphthalen-1-yl)ethan-1-amine (35 mg, 0.20 mmol), 2-((1-(tert-butoxycarbonyl)azetidin-3-yl)amino)-5-methylisonicotinic acid (30 mg, 0.10 mmol), HATU (46 mg, 0.12 mmol) and DMAP (37 mg, 0.30 mmol) was subjected to general amine coupling procedure with DMF (1.5 mL). After purification by Prep-HPLC, the product was subjected to general N-Boc deprotection procedure with HCl (4M in dioxane, 100 µL) and DCM (2 mL). The purification by Prep-HPLC afforded the product **ZN-3-55** (31 mg, yield 86% for 2 steps) as a white solid: $[\alpha]^{25}_{546}$ = -88.9 (c 1.0, MeOH); $^{1}$H NMR (400 MHz, Methanol-$d_4$) δ 8.45 (s, 1H), 8.24 – 8.17 (m, 1H), 7.99 – 7.79 (m, 3H), 7.66 – 7.46 (m, 4H), 6.95 (d, *J* = 5.8 Hz, 1H), 6.06 – 5.99 (m, 1H), 4.82 – 4.47 (m, 3H), 4.36 – 4.05 (m, 1H), 3.24 – 3.12 (m, 1H), 2.23 – 2.15 (m, 3H), 1.77 – 1.72 (m, 3H); $^{13}$C NMR (100 MHz, Methanol-$d_4$) δ 169.57, 166.46, 155.84, 154.27, 149.40, 139.32, 137.43, 135.49, 132.26, 130.04, 129.38, 127.47, 126.89, 126.43, 124.03, 123.93, 108.11, 55.83, 55.40, 54.39, 46.47, 21.02, 15.45; HRMS (ESI) calcd for $C_{22}H_{25}N_4O$ $[M+H]^+$ 361.2023, found 361.2024.

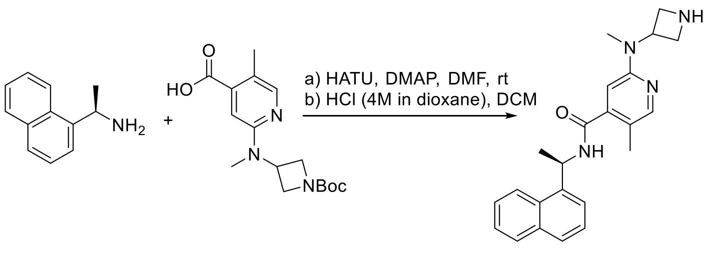


**ZN-3-57**

**(R)-2-(azetidin-3-yl(methyl)amino)-5-methyl-N-(1-(naphthalen-1-yl)ethyl)isonicotinamide (ZN-3-57)**. (R)-1-(naphthalen-1-yl)ethan-1-amine (42 mg, 0.24 mmol), 2-((1-(tert-butoxycarbonyl)azetidin-3-yl)(methyl)amino)-5-methylisonicotinic acid (50 mg, 0.16 mmol), HATU (73 mg, 0.19 mmol) and DMAP (39 mg, 0.32 mmol) was subjected to general amine coupling procedure with DMF (1 mL). After purification by Prep-HPLC, the product was subjected to general N-Boc deprotection procedure with HCl (4M in dioxane, 200 µL) and DCM (4 mL). The purification by Prep-HPLC afforded the product **ZN-3-57** (44 mg, yield 73% for 2 steps) as a white solid: $[\alpha]^{25}_{546}$ = -109.3 (c 0.2, MeOH); $^{1}$H NMR (400 MHz, Methanol-$d_4$) δ 8.42 (s, 1H), 8.22 (d, *J* = 8.5 Hz, 1H), 7.98 (s, 1H), 7.92 (d, *J* = 8.1 Hz, 1H), 7.85 (d, *J* = 8.2 Hz, 1H), 7.68 – 7.63 (m, 1H), 7.62 – 7.48 (m, 3H), 7.05 (d, *J* = 4.7 Hz, 1H), 6.06 (q, *J* = 6.9 Hz, 1H), 4.82 – 4.75 (m, 1H), 4.63 – 4.54 (m, 1H), 4.32 (d, *J* = 6.1 Hz, 1H), 3.23 – 3.15 (m, 1H), 3.13 (s, 3H), 3.08 – 3.02 (m, 1H), 2.17 (d, *J* = 4.8 Hz, 3H), 1.76 (d, *J* = 6.9 Hz, 3H); $^{13}$C NMR (100 MHz, Methanol-$d_4$) δ 169.57, 165.77, 161.93, 144.52, 137.97, 135.38, 132.30, 130.06, 129.41, 127.48, 126.92, 126.45, 124.09, 123.99, 122.19, 106.54, 54.03, 46.52, 41.31, 31.14, 21.11, 15.38; HRMS (ESI) calcd for $C_{23}H_{27}N_4O$ $[M+H]^+$ 375.2179, found 375.2185.

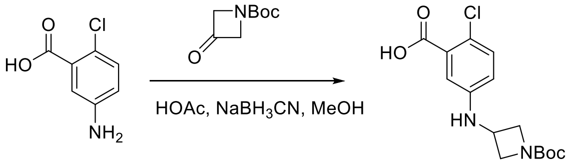


**S6**

**5-((1-(tert-butoxycarbonyl)azetidin-3-yl)amino)-2-chlorobenzoic acid (S6)**. 5-amino-2-chlorobenzoic acid (3.0 g, 17.5 mmol) and tert-butyl 3-oxoazetidine-1-carboxylate (3.6 g, 21.0 mmol) was subjected to general reductive amination procedure with MeOH (20 mL), HOAc (4 mL) at 50$^{o}$C, and then $NaBH_3CN$ (3.3 g, 52.5 mmol) was added. The purification by silica gel column chromatography (Hexenes/EtOAc, 1:1) provided the amination compound **S6** (4.8 g, yield 84%) as a white solid: $^{1}$H NMR (400 MHz, Methanol-$d_4$) δ 7.20 (d, *J* = 8.7 Hz, 1H), 6.96 (d, *J* = 2.9 Hz, 1H), 6.64 (dd, *J* = 8.7, 2.9 Hz, 1H), 4.30 – 4.16 (m, 3H), 3.72 (dd, *J* = 8.7, 4.5 Hz, 2H), 1.44 (s, 9H); LRMS (ESI) calcd for $C_{15}H_{20}ClN_2O_4$ $[M+H]^+$ 327.1, found 327.2.

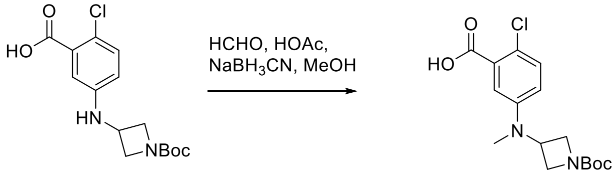


**S7**

**5-((1-(tert-butoxycarbonyl)azetidin-3-yl)(methyl)amino)-2-chlorobenzoic acid (S7)**. 5-((1-(tert-butoxycarbonyl)azetidin-3-yl)amino)-2-chlorobenzoic acid (1.8 g, 5.5 mmol) and formaldehyde (37 wt. % in $H_2O$, 2 mL) was subjected to general reductive amination procedure with MeOH (10 mL), HOAc (2 mL) at room temperature, and then $NaBH_3CN$ (1.1 g, 16.5 mmol) was added. The purification by silica gel column chromatography (Hexenes/EtOAc, 2:1) provided the amination compound **S7** (1.7 g, yield 91%) as a white solid: $^1H$ NMR (400 MHz, Methanol-*d*$_4$) δ 7.25 (d, *J* = 8.8 Hz, 1H), 7.15 (d, *J* = 3.1 Hz, 1H), 6.83 (dd, *J* = 8.9, 3.1 Hz, 1H), 4.34 (tt, *J* = 7.5, 5.3 Hz, 1H), 4.19 – 4.13 (m, 2H), 3.88 (dd, *J* = 9.1, 5.3 Hz, 2H), 2.88 (s, 3H), 1.43 (s, 9H); $^{13}C$ NMR (100 MHz, Methanol-*d*$_4$) δ 169.12, 157.88, 149.62, 132.36, 123.42, 120.40, 118.49, 81.17, 54.89, 50.22, 35.75, 28.63; LRMS (ESI) calcd for $C_{16}H_{22}ClN_2O_4$ $[M+H]^+$ 341.1, found 341.1.

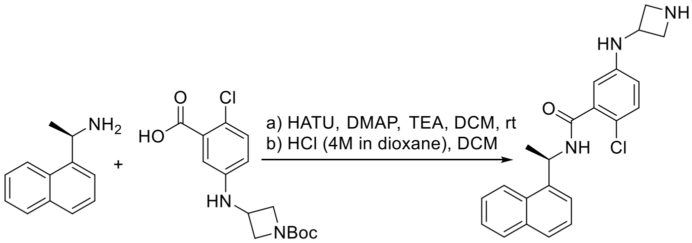


S6

ZN-3-66

**(R)-5-(azetidin-3-ylamino)-2-chloro-N-(1-(naphthalen-1-yl)ethyl)benzamide (ZN-3-66)**. (R)-1-(naphthalen-1-yl)ethan-1-amine (39 mg, 0.23 mmol), 5-((1-(tert-butoxycarbonyl)azetidin-3-yl)amino)-2-chlorobenzoic acid (**S6**) (50 mg, 0.15 mmol), HATU (70 mg, 0.18 mmol) and DMAP (56 mg, 0.46 mmol) was subjected to general amine coupling procedure with DMF (2 mL). After purification by Prep-HPLC, the product was subjected to general N-Boc deprotection procedure with HCl (4M in dioxane, 200 µL) and DCM (4 mL). The purification by Prep-HPLC afforded the product **ZN-3-66** (43 mg, yield 75% for 2 steps) as a white solid: $[\alpha]^{25}_{546}$ = -54.0 (c 2.3, MeOH); $^1H$ NMR (400 MHz, Methanol-*d*$_4$) δ 8.51 (s, 1H), 8.25 – 8.20 (m, 1H), 7.92 – 7.87 (m, 1H), 7.80 (d, *J* = 8.2 Hz, 1H), 7.67 – 7.63 (m, 1H), 7.59 – 7.44 (m, 3H), 7.18 (d, *J* = 8.7 Hz, 1H), 6.62 – 6.53 (m, 2H), 6.02 (q, *J* = 6.9 Hz, 1H), 4.48 – 4.39 (m, 1H), 4.34 – 4.26 (m, 2H), 3.94 – 3.87 (m, 2H), 1.69 (d, *J* = 7.0 Hz, 3H); $^{13}C$ NMR (100 MHz, Methanol-*d*$_4$) δ 169.24, 146.53, 140.08, 138.22, 135.42, 132.27, 131.68, 129.89, 128.98, 127.24, 126.70, 126.42, 124.31, 123.87, 120.16, 116.24, 113.82, 54.65, 46.71, 46.49, 21.41; HRMS (ESI) calcd for $C_{22}H_{23}ClN_3O$ $[M+H]^+$ 380.1524, found 388.1532.

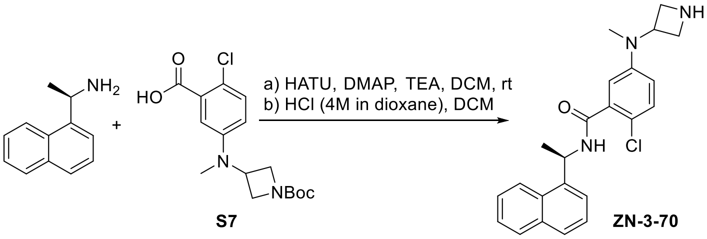


S7

ZN-3-70

**(R)-5-(azetidin-3-yl(methyl)amino)-2-chloro-N-(1-(naphthalen-1-yl)ethyl)benzamide (ZN-3-70)**. (R)-1-(naphthalen-1-yl)ethan-1-amine (39 mg, 0.23 mmol), 5-((1-(tert-butoxycarbonyl)azetidin-3-yl)(methyl)amino)-2-chlorobenzoic acid (**S7**) (51 mg, 0.15 mmol), HATU (70 mg, 0.18 mmol) and DMAP (56 mg, 0.46 mmol) was subjected to general amine coupling procedure with DMF (2 mL). After purification by Prep-HPLC, the product was subjected to general N-Boc deprotection procedure with HCl (4M in dioxane, 200 µL) and DCM (4 mL). The purification by Prep-HPLC afforded the **ZN-3-70** (49 mg, yield 83% for 2 steps) as a white solid: $[\alpha]^{25}_{546}$ = -56.6 (c 1.3, MeOH); $^1H$ NMR (400 MHz, Methanol-*d*$_4$) δ 8.50 (s, 1H), 8.26 – 8.22 (m, 1H), 7.92 – 7.88 (m, 1H), 7.81 (d, *J* = 8.2 Hz, 1H), 7.67 – 7.64 (m, 1H), 7.60 – 7.45 (m,

3H), 7.29 (d, $J$ = 8.8 Hz, 1H), 6.85 (dd, $J$ = 8.8, 3.1 Hz, 1H), 6.76 (d, $J$ = 3.0 Hz, 1H), 6.02 (q, $J$ = 6.9 Hz, 1H), 4.65 – 4.56 (m, 1H), 4.24 – 4.17 (m, 2H), 4.14 – 4.09 (m, 2H), 2.89 (s, 3H), 1.71 (d, $J$ = 6.9 Hz, 3H); $^{13}$C NMR (100 MHz, Methanol-$d_4$) δ 169.63, 149.30, 140.02, 138.14, 135.45, 132.28, 131.65, 129.91, 129.02, 127.24, 126.72, 126.43, 124.33, 123.93, 122.34, 119.54, 117.04, 52.73, 51.96, 46.81, 35.70, 21.39; HRMS (ESI) calcd for $C_{23}H_{25}ClN_3O$ [M+H]$^+$ 394.1681, found 394.1677.

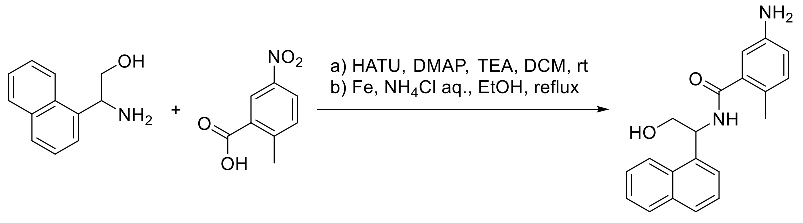


**ZN-3-13**

**5-amino-N-(2-hydroxy-1-(naphthalen-1-yl)ethyl)-2-methylbenzamide (ZN-3-13)**. 2-amino-2-(naphthalen-1-yl)ethan-1-ol (100 mg, 0.53 mmol), 2-methyl-5-nitrobenzoic acid (87 mg, 0.48 mmol), HATU (202 mg, 0.53 mmol), TEA (200 µL), and DMAP (6 mg, 0.05 mmol) was subjected to general amine coupling procedure with DCM (4 mL). After purification by Prep-HPLC, the product was applied to the general Aryl Nitro reduction procedure with ethanol/ saturated aq. $NH_4Cl$ (4 mL/1 mL) and Iron Powder (148 mg, 2.65 mmol). The purification by Prep-HPLC gave the **ZN-3-13** (92 mg, yield 60% for 2 steps) as a white solid: $^1$H NMR (400 MHz, Methanol-$d_4$) δ 8.34 – 8.27 (m, 1H), 7.92 – 7.85 (m, 1H), 7.84 – 7.77 (m, 1H), 7.67 – 7.42 (m, 5H), 7.25 – 6.95 (m, 1H), 6.80 – 6.67 (m, 1H), 6.10 – 6.02 (m, 1H), 4.08 – 3.99 (m, 1H), 3.94 – 3.83 (m, 1H), 2.27 (s, 3H); $^{13}$C NMR (100 MHz, Methanol-$d_4$) δ 173.30, 146.27, 138.45, 136.75, 135.38, 132.31, 129.91, 129.10, 127.35, 126.72, 126.34, 124.77, 124.06, 122.34, 119.79, 118.10, 115.19, 65.35, 53.20, 18.74; HRMS (ESI) calcd for $C_{20}H_{21}N_2O_2$ [M+H]$^+$ 321.1598, found 308.1589.

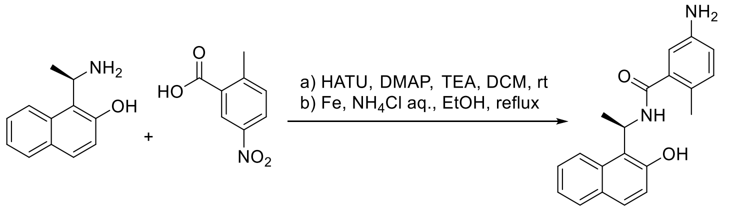


**ZN-3-19**

**(R)-5-amino-N-(1-(2-hydroxynaphthalen-1-yl)ethyl)-2-methylbenzamide (ZN-3-19)**. (R)-1-(1-aminoethyl)naphthalen-2-ol (100 mg, 0.53 mmol), 2-methyl-5-nitrobenzoic acid (87 mg, 0.48 mmol), HATU (202 mg, 0.53 mmol), TEA (200 µL), and DMAP (6 mg, 0.05 mmol) was subjected to general amine coupling procedure with DCM (4 mL). After purification by Prep-HPLC, the product was applied to the general Aryl Nitro reduction procedure with ethanol/ saturated aq. $NH_4Cl$ (4 mL/1 mL) and Iron Powder (148 mg, 2.65 mmol). The purification by Prep-HPLC gave the **ZN-3-19** (87 mg, yield 57% for 2 steps) as a light brown solid: $[\alpha]^{25}_{546}$ = -62.4 (c 0.3, MeOH); $^1$H NMR (400 MHz, Methanol-$d_4$) δ 8.15 (d, $J$ = 8.8 Hz, 1H), 7.77 (dd, $J$ = 8.2, 1.4 Hz, 1H), 7.69 (d, $J$ = 8.9 Hz, 1H), 7.50 (ddd, $J$ = 8.5, 6.8, 1.4 Hz, 1H), 7.35 – 7.26 (m, 1H), 7.12 (d, $J$ = 8.8 Hz, 1H), 6.96 (d, $J$ = 8.1 Hz, 1H), 6.77 – 6.67 (m, 2H), 6.19 (q, $J$ = 7.0 Hz, 1H), 2.21 (s, 3H), 1.65 (d, $J$ = 7.0 Hz, 3H); $^{13}$C NMR (100 MHz, Methanol-$d_4$) δ 171.98, 153.95, 146.47, 138.44, 133.26, 132.57, 130.42, 130.13, 129.73, 127.76, 125.65, 123.92, 122.98, 120.89, 119.26, 118.30, 114.94, 44.76, 20.68, 18.67; HRMS (ESI) calcd for $C_{20}H_{21}N_2O_2$ [M+H]$^+$ 321.1598, found 321.1599.

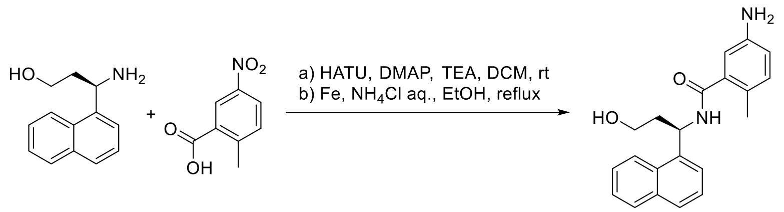


DY2-97

**(R)-5-amino-N-(3-hydroxy-1-(naphthalen-1-yl)propyl)-2-methylbenzamide (DY2-97)**. (R)-3-amino-3-(naphthalen-1-yl)propan-1-ol (100 mg, 0.50 mmol), 2-methyl-5-nitrobenzoic acid (60 mg, 0.33 mmol), HATU (124 mg, 0.33 mmol), TEA (100 µL), and DMAP (5 mg, 0.04 mmol) was subjected to general amine coupling procedure with DCM (4 mL). After purification by Prep-HPLC, the product was applied to the general Aryl Nitro reduction procedure with ethanol/ saturated aq. $NH_4Cl$ (4 mL/1 mL) and Iron Powder (92 mg, 1.65 mmol). The purification by Prep-HPLC gave the **DY2-97** (62 mg, yield 56% for 2 steps) as a white solid: $^1$H NMR (400 MHz, Methanol-$d_4$) δ 8.33 (dd, $J$ = 8.7, 4.8 Hz, 1H), 7.91 – 7.86 (m, 1H), 7.81 (d, $J$ = 8.2 Hz, 1H), 7.64 – 7.45 (m, 5H), 6.98 – 6.95 (m, 1H), 6.73 – 6.69 (m, 2H), 6.15 – 6.07 (m, 1H), 3.85 – 3.71 (m, 2H), 2.29 – 2.12 (m, 5H); $^{13}$C NMR (100 MHz, Methanol-$d_4$) δ 173.07, 146.00, 139.67, 138.40, 135.43, 132.38, 129.87, 128.90, 127.28, 126.75, 126.39, 125.88, 124.30, 124.10, 118.30, 115.25, 60.12, 47.73, 39.41, 18.73; LRMS (ESI) calcd for $C_{21}H_{23}N_2O_2$ [M+H]$^+$ 335.2, found 335.2.

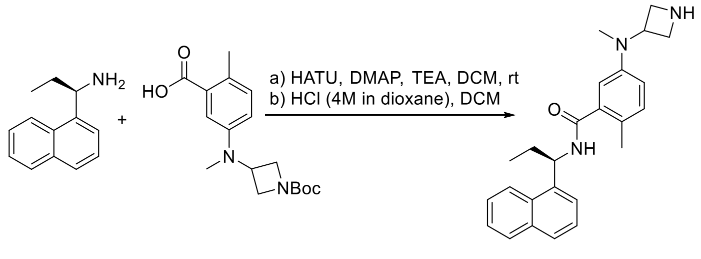


ZN-3-61

**(R)-5-(azetidin-3-yl(methyl)amino)-2-methyl-N-(1-(naphthalen-1-yl)propyl)benzamide (ZN-3-61)**. (R)-1-(naphthalen-1-yl)propan-1-amine (17 mg, 0.09 mmol), 5-((1-(tert-butoxycarbonyl)azetidin-3-yl)(methyl)amino)-2-methylbenzoic acid (20 mg, 0.06 mmol), HATU (24 mg, 0.06 mmol) and DMAP (15 mg, 0.13 mmol) was subjected to general amine coupling procedure with DMF (1 mL). After purification by Prep-HPLC, the product was subjected to general N-Boc deprotection procedure with HCl (4M in dioxane, 100 µL) and DCM (2 mL). The purification by Prep-HPLC afforded the product **ZN-3-61** (18 mg, yield 77% for 2 steps) as a white solid: $[\alpha]^{25}_{546}$= -65.4 (c 0.9, MeOH); $^1$H NMR (400 MHz, Methanol-$d_4$) δ 8.40 (s, 1H), 8.30 (d, $J$ = 8.5 Hz, 1H), 7.92 – 7.89 (m, 1H), 7.81 (d, $J$ = 8.2 Hz, 1H), 7.60 – 7.46 (m, 4H), 7.12 (d, $J$ = 8.4 Hz, 1H), 6.81 (dd, $J$ = 8.4, 2.7 Hz, 1H), 6.69 (d, $J$ = 2.7 Hz, 1H), 5.84 (dd, $J$ = 9.0, 5.7 Hz, 1H), 4.51 – 4.41 (m, 1H), 4.22 – 4.15 (m, 2H), 4.09 – 4.01 (m, 2H), 2.83 (s, 3H), 2.23 (s, 3H), 2.15 – 1.95 (m, 2H), 1.13 (t, $J$ = 7.3 Hz, 3H); $^{13}$C NMR (100 MHz, Methanol-$d_4$) δ 172.56, 148.38, 139.84, 139.05, 135.48, 132.60, 129.96, 128.89, 128.49, 127.22, 126.72, 126.41, 124.30, 124.26, 119.66, 116.94, 53.33, 52.51, 51.97, 37.10, 29.84, 18.73, 11.92; HRMS (ESI) calcd for $C_{25}H_{30}N_3O$ [M+H]$^+$ 388.2383, found 388.2389.

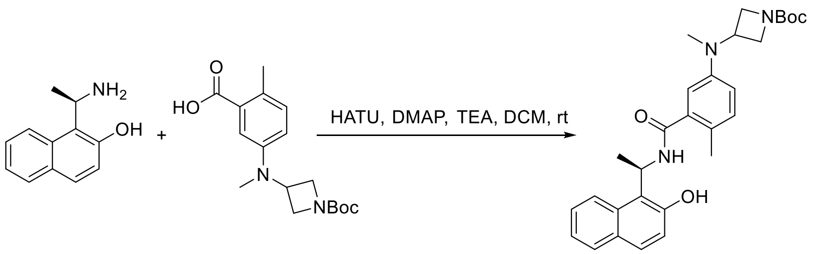


S8

**Tert-butyl (R)-3-((3-((1-(2-hydroxynaphthalen-1-yl)ethyl)carbamoyl)-4-methylphenyl)(methyl)amino)azetidine-1-carboxylate (S8)**. (R)-1-(1-aminoethyl)naphthalen-2-ol (178 mg, 0.95 mmol), 5-((1-(tert-butoxycarbonyl)azetidin-3-yl)(methyl)amino)-2-methylbenzoic acid (254 mg, 0.79 mmol), HATU (300 mg, 0.79 mmol), TEA (500 µL), and DMAP (10 mg, 0.08 mmol) was subjected to general amine coupling procedure with DCM (4 mL). The purification by Prep-HPLC gave the desired compound **S8** (325 mg, yield 84%) as a white solid: $^{1}$H NMR (400 MHz, Chloroform-*d*) δ 9.07 (s, 1H), 8.15 (d, *J* = 8.7 Hz, 1H), 7.81 – 7.72 (m, 1H), 7.61 (d, *J* = 8.8 Hz, 1H), 7.49 – 7.44 (m, 1H), 7.34 – 7.30 (m, 1H), 7.18 (d, *J* = 8.8 Hz, 1H), 7.01 (d, *J* = 8.3 Hz, 1H), 6.75 – 6.56 (m, 2H), 6.31 – 6.19 (m, 1H), 4.17 – 4.04 (m, 3H), 3.84 (td, *J* = 8.1, 7.6, 5.3 Hz, 2H), 2.70 (s, 3H), 2.32 (s, 3H), 1.65 (d, *J* = 6.9 Hz, 3H), 1.45 (s, 9H); LRMS (ESI) calcd for $C_{29}H_{36}N_{3}O_{4}$ $[M+H]^{+}$ 490.3, found 490.3.

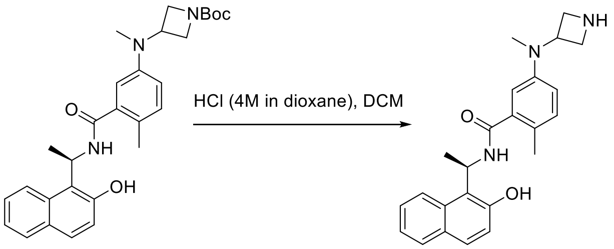


S8 ZN-3-32

**(R)-5-(azetidin-3-yl(methyl)amino)-N-(1-(2-hydroxynaphthalen-1-yl)ethyl)-2-methylbenzamide (ZN-3-32)**. General procedure for N-Boc deprotection was used with tert-butyl (R)-3-((3-((1-(2-hydroxynaphthalen-1-yl)ethyl)carbamoyl)-4-methylphenyl)(methyl)amino)azetidine-1-carboxylate (**S8**) (40 mg, 0.08 mmol) in DCM (2 mL) and HCl (4M in dioxane, 200 µL). The purification by Prep-HPLC afforded the product **ZN-3-32** (23 mg, yield 72%) as a light brown solid: $[\alpha]_{546}^{25}$ = -70.4 (c 0.4, MeOH); $^{1}$H NMR (400 MHz, Methanol-$d_4$) δ 8.55 (s, 1H), 8.17 (d, *J* = 8.7 Hz, 1H), 7.81 – 7.75 (m, 1H), 7.70 (d, *J* = 8.8 Hz, 1H), 7.53 – 7.48 (m, 1H), 7.33 – 7.29 (m, 1H), 7.16 – 7.12 (m, 2H), 6.83 (dd, *J* = 8.4, 2.7 Hz, 1H), 6.78 – 6.76 (m, 1H), 6.20 (q, *J* = 6.9 Hz, 1H), 4.47 (p, *J* = 7.2 Hz, 1H), 4.23 – 4.16 (m, 2H), 4.07 – 4.01 (m, 2H), 2.85 (s, 3H), 2.27 (s, 3H), 1.67 (d, *J* = 7.0 Hz, 3H); $^{13}$C NMR (100 MHz, Methanol-$d_4$) δ 169.52, 153.96, 148.60, 138.71, 133.28, 132.89, 130.43, 130.19, 129.78, 128.42, 127.74, 123.92, 123.01, 120.93, 119.94, 119.25, 116.70, 53.46, 52.12, 44.89, 37.05, 20.51, 18.74; HRMS (ESI) calcd for $C_{24}H_{28}N_{3}O_{2}$ $[M+H]^{+}$ 390.2176, found 390.2185.

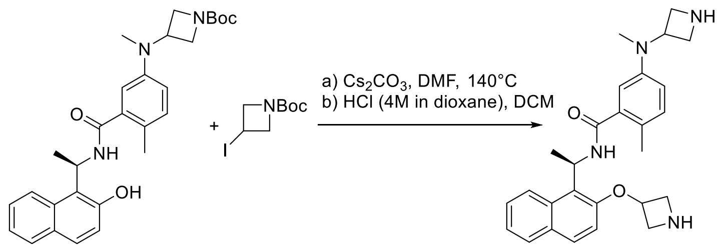


**S8** **ZN-3-33**

**(R)-5-(azetidin-3-yl(methyl)amino)-N-(1-(2-(azetidin-3-yloxy)naphthalen-1-yl)ethyl)-2-methylbenzamide (ZN-3-33)**. To a solution of tert-butyl (R)-3-((3-((1-(2-hydroxynaphthalen-1-yl)ethyl)carbamoyl)-4-methylphenyl)(methyl)amino)azetidine-1-carboxylate (**S8**) (20 mg, 0.04 mmol) and tert-butyl 3-iodoazetidine-1-carboxylate (14 mg, 0.05 mol) in DMF (1 mL), $Cs_2CO_3$ (26 mg, 0.08 mmol) was added. The reaction was stirred at 140 °C for 5 h. After cooling down, quench the reaction with MeOH (1 mL). The mixture was diluted with Ethyl Acetate and was then washed with saturated aq. $NaHCO_3$, water, and brine, respectively. The organic layer was dried over $Na_2SO_4$, filtered, and concentrated. After purification by Prep-HPLC, the product was applied to the general procedure for N-Boc deprotection with HCl (4M in dioxane, 50 µL) and DCM (1 mL). The purification by Prep-HPLC afforded the product **ZN-3-33** (8 mg, yield 45% for 2 steps) as a light brown solid: $[\alpha]^{25}_{546}$ = -87 (c 0.6, MeOH); $^1$H NMR (400 MHz, Methanol-$d_4$) δ 8.53 (s, 1H), 8.41 (d, $J$ = 8.7 Hz, 1H), 7.91 – 7.82 (m, 2H), 7.57 – 7.52 (m, 1H), 7.46 – 7.06 (m, 3H), 6.80 – 6.75 (m, 1H), 6.65 – 6.62 (m, 1H), 6.24 (dq, $J$ = 9.8, 7.4 Hz, 1H), 5.44 – 5.09 (m, 1H), 4.59 – 4.28 (m, 3H), 4.22 – 4.12 (m, 2H), 4.07 – 3.97 (m, 3H), 3.56 – 3.43 (m, 1H), 2.84 (s, 3H), 2.10 (d, $J$ = 9.0 Hz, 3H), 1.77 (d, $J$ = 7.4 Hz, 3H); $^{13}$C NMR (100 MHz, Methanol-$d_4$) δ 172.23, 152.57, 148.36, 146.49, 138.90, 138.51, 132.55, 131.04, 130.42, 130.28, 127.61, 125.15, 125.05, 119.45, 116.92, 115.78, 114.98, 70.75, 54.93, 54.77, 53.39, 52.07, 44.41, 36.58, 20.06, 18.47; HRMS (ESI) calcd for $C_{27}H_{33}N_4O_2$ $[M+H]^+$ 445.2598, found 445.2587.

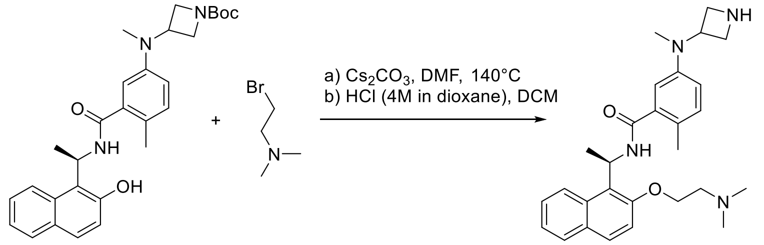


**S8** **ZN-3-34**

**(R)-5-(azetidin-3-yl(methyl)amino)-N-(1-(2-(2-(dimethylamino)ethoxy)naphthalen-1-yl)ethyl)-2-methylbenzamide (ZN-3-34)**. To a solution of tert-butyl (R)-3-((3-((1-(2-hydroxynaphthalen-1-yl)ethyl)carbamoyl)-4-methylphenyl)(methyl)amino)azetidine-1-carboxylate (**S8**) (20 mg, 0.04 mmol) and 2-bromo-N,N-dimethylethan-1-amine (12 mg, 0.05 mol) in DMF (1 mL), $Cs_2CO_3$ (26 mg, 0.08 mmol) was added. The reaction was stirred at 140 °C for 5 h. After cooling down, quench the reaction with MeOH (1 mL). The mixture was diluted with Ethyl Acetate and was then washed with saturated aq. $NaHCO_3$, water, and brine, respectively. The organic layer was dried over $Na_2SO_4$, filtered, and concentrated. After purification by Prep-HPLC, the product was applied to the general procedure for N-Boc deprotection with HCl (4M in dioxane, 50 µL) and DCM (1 mL). The purification by Prep-HPLC afforded the product **ZN-3-34** (9 mg, yield 49% for 2 steps) as a light brown solid: $[\alpha]^{25}_{546}$ = -57.8 (c 0.5, MeOH); $^1$H NMR (400 MHz, Methanol-$d_4$) δ 8.53 (s, 1H), 8.31 (d, $J$ = 8.9 Hz, 1H), 7.88 – 7.83 (m, 2H), 7.58 – 7.52 (m,

1H), 7.44 – 7.36 (m, 2H), 7.12 (d, *J* = 8.4 Hz, 1H), 6.80 (dd, *J* = 8.4, 2.7 Hz, 1H), 6.65 (d, *J* = 2.7 Hz, 1H), 6.36 (q, *J* = 7.3 Hz, 1H), 4.51 – 3.99 (m, 9H), 2.83 (s, 3H), 2.26 – 2.03 (m, 9H), 1.72 (d, *J* = 7.1 Hz, 3H); $^{13}$C NMR (100 MHz, Methanol-$d_4$) δ 172.40, 148.53, 132.90, 132.54, 130.75, 130.01, 128.07, 127.85, 124.74, 120.95, 119.30, 116.70, 115.42, 66.20, 59.41, 53.35, 52.08, 45.01, 44.06, 36.94, 20.61, 18.43; HRMS (ESI) calcd for $C_{28}H_{37}N_4O_2$ [M+H]$^+$ 461.2911, found 461.2918.

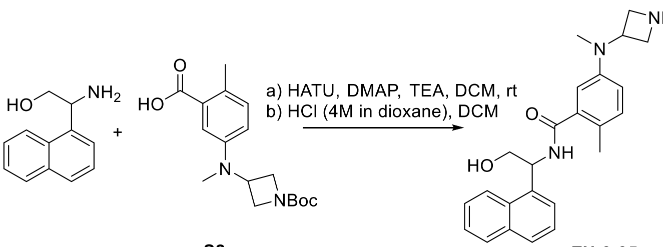


S2 ZN-3-35

**5-(azetidin-3-yl(methyl)amino)-N-(2-hydroxy-1-(naphthalen-1-yl)ethyl)-2-methylbenzamide (ZN-3-35)**. 2-amino-2-(naphthalen-1-yl)ethan-1-ol (100 mg, 0.53 mmol), 5-((1-(tert-butoxycarbonyl)azetidin-3-yl)(methyl)amino)-2-methylbenzoic acid (**S2**) (170 mg, 0.53 mmol), HATU (201 mg, 0.53 mmol), DMAP (6 mg, 0.05 mmol) and TEA (200 µL) was subjected to general amine coupling procedure with DCM (4 mL). After purification by Prep-HPLC, the product was subjected to general N-Boc deprotection procedure with HCl (4M in dioxane, 200 µL) and DCM (4 mL). The purification by Prep-HPLC afforded the product **ZN-2-35** (151 mg, yield 73% for 2 steps) as a white solid: $^1$H NMR (400 MHz, Methanol-$d_4$) δ 8.62 – 8.59 (m, 1H), 8.34 – 8.29 (m, 1H), 7.94 – 7.89 (m, 1H), 7.85 – 7.81 (m, 1H), 7.63 – 7.45 (m, 4H), 7.09 (d, *J* = 8.1 Hz, 1H), 6.85 – 6.79 (m, 2H), 6.09 – 6.03 (m, 1H), 4.53 – 4.47 (m, 1H), 4.29 – 4.21 (m, 2H), 4.11 – 4.02 (m, 3H), 3.90 (dd, *J* = 11.5, 8.3 Hz, 1H), 2.87 (s, 3H), 2.26 (s, 3H); $^{13}$C NMR (100 MHz, Methanol-$d_4$) δ 162.40, 148.38, 138.93, 138.53, 135.46, 132.64, 132.19, 129.99, 129.07, 128.88, 127.41, 126.82, 126.34, 124.29, 124.05, 119.96, 117.17, 65.42, 53.43, 52.22, 46.99, 37.30, 18.75; HRMS (ESI) calcd for $C_{28}H_{37}N_4O_2$ [M+H]$^+$ 461.2911, found 461.2918.

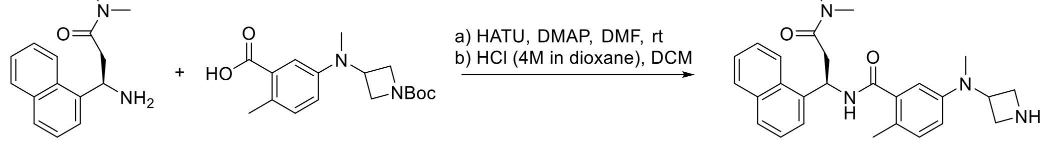


S2 DY2-117

**(R)-5-(azetidin-3-yl(methyl)amino)-N-(3-(dimethylamino)-1-(naphthalen-1-yl)-3-oxopropyl)-2-methylbenzamide (DY2-117)**. (R)-3-amino-N,N-dimethyl-3-(naphthalen-1-yl)propanamide (36 mg, 0.15 mmol), 5-((1-(tert-butoxycarbonyl)azetidin-3-yl)(methyl)amino)-2-methylbenzoic acid (**S2**) (48 mg, 0.15 mmol), HATU (70 mg, 0.18 mmol) and DMAP (56 mg, 0.46 mmol) was subjected to general amine coupling procedure with DMF (2 mL). After purification by Prep-HPLC, the product was subjected to general N-Boc deprotection procedure with HCl (4M in dioxane, 200 µL) and DCM (4 mL). The purification by Prep-HPLC afforded the product **DY2-117** (57 mg, yield 86% for 2 steps) as a white solid: $^1$H NMR (400 MHz, Methanol-$d_4$) δ 8.55 (s, 1H), 8.31 (d, *J* = 8.5 Hz, 1H), 7.97 – 7.82 (m, 2H), 7.63 – 7.45 (m, 4H), 7.12 (d, *J* = 8.3 Hz, 1H), 6.82 (dd, *J* = 8.3, 2.6 Hz, 1H), 6.76 (d, *J* = 2.6 Hz, 1H), 6.44 (dd, *J* = 8.6, 5.5 Hz, 1H), 4.49 – 4.37 (m, 1H), 4.21 – 4.12 (m, 2H), 4.06 – 3.95 (m, 2H), 3.20 – 3.05 (m, 2H), 2.99 (s, 3H), 2.92 (s, 3H), 2.84 (s, 3H), 2.24 (s, 3H); $^{13}$C NMR (100 MHz, Methanol-$d_4$) δ 172.49, 172.13, 135.54, 132.63, 132.19, 130.05, 129.30, 127.52, 126.93, 126.38, 124.30, 124.15, 119.78, 116.85,

53.80, 52.25, 39.61, 37.90, 37.17, 35.92, 18.68; HRMS (ESI) calcd for $C_{27}H_{33}N_4O_2$ $[M+H]^+$ 445.2598, found 445.2590.

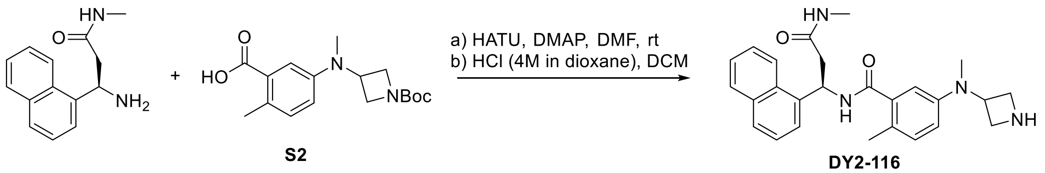


**(R)-3-(azetidin-3-yl(methyl)amino)-N-(3-(methylamino)-1-(naphthalen-1-yl)-3-oxopropyl)benzamide** (**DY2-116**). (R)-3-amino-N-methyl-3-(naphthalen-1-yl)propanamide (34 mg, 0.15 mmol), 5-((1-(tert-butoxycarbonyl)azetidin-3-yl)(methyl)amino)-2-methylbenzoic acid (**S2**) (48 mg, 0.15 mmol), HATU (70 mg, 0.18 mmol) and DMAP (56 mg, 0.46 mmol) was subjected to general amine coupling procedure with DMF (2 mL). After purification by Prep-HPLC, the product was subjected to general N-Boc deprotection procedure with HCl (4M in dioxane, 200 µL) and DCM (4 mL). The purification by Prep-HPLC afforded the product **DY2-116** (45 mg, yield 70% for 2 steps) as a white solid: $[\alpha]^{25}_{546}$= -17.9 (c 0.5, MeOH); $^1$H NMR (400 MHz, Methanol-$d_4$) δ 8.55 (s, 1H), 8.37 (d, $J$ = 8.1 Hz, 1H), 7.92 (d, $J$ = 7.8 Hz, 1H), 7.83 (d, $J$ = 8.8 Hz, 1H), 7.63 – 7.45 (m, 4H), 7.12 (d, $J$ = 8.1 Hz, 1H), 6.84 – 6.80 (m, 1H), 6.76 – 6.74 (m, 1H), 6.43 – 6.38 (m, 1H), 4.48 (p, $J$ = 7.3 Hz, 1H), 4.20 – 4.15 (m, 2H), 4.06 – 3.99 (m, 2H), 2.98 – 2.81 (m, 5H), 2.71 (s, 3H), 2.23 (s, 3H); $^{13}$C NMR (100 MHz, Methanol-$d_4$) δ 173.41, 172.04, 148.51, 138.64, 138.55, 135.51, 132.67, 132.15, 129.98, 129.30, 128.52, 127.48, 126.90, 126.35, 124.25, 119.81, 116.82, 53.68, 52.24, 42.52, 37.09, 26.45, 18.69; HRMS (ESI) calcd for $C_{26}H_{31}N_4O_2$ $[M+H]^+$ 431.2442, found 431.2441.

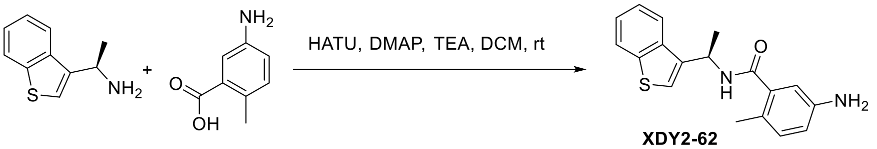


**(R)-5-amino-N-(1-(benzo[b]thiophen-3-yl)ethyl)-2-methylbenzamide (XDY2-62)**. (R)-1-(benzo[b]thiophen-3-yl)ethan-1-amine (57 mg, 0.32 mmol), 5-amino-2-methylbenzoic acid (24 mg, 0.16 mmol), HATU (61 mg, 0.16 mmol), TEA (100 µL), and DMAP (3 mg, 0.02 mmol) was subjected to general amine coupling procedure with DCM (2 mL). The purification by Prep-HPLC afforded the product **XDY2-62** (41 mg, yield 83%) as a light brown solid: $[\alpha]^{25}_{546}$= -34.7 (c 1.4, MeOH); $^1$H NMR (400 MHz, Chloroform-$d$) δ 7.97 – 7.92 (m, 1H), 7.88 – 7.83 (m, 1H), 7.45 – 7.30 (m, 3H), 6.96 – 6.91 (m, 1H), 6.60 – 6.55 (m, 2H), 5.80 – 5.69 (m, 1H), 2.29 (s, 3H), 1.75 (d, $J$ = 6.7 Hz, 3H); $^{13}$C NMR (100 MHz, Chloroform-$d$) δ 172.91, 144.23, 140.71, 137.90, 131.99, 125.47, 124.89, 124.53, 122.98, 122.46, 122.35, 116.86, 116.82, 113.47, 113.44, 43.15, 20.44, 18.87; HRMS (ESI) calcd for $C_{18}H_{18}N_3O_3$ $[M+H]^+$ 324.1343, found 324.1341.

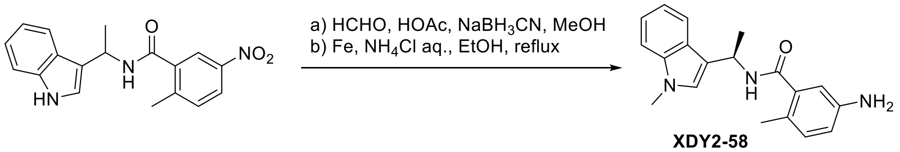


**(R)-5-amino-2-methyl-N-(1-(1-methyl-1H-indol-3-yl)ethyl)benzamide** (**XDY2-58**). N-(1-(1H-indol-3-yl)ethyl)-2-methyl-5-nitrobenzamide (50 mg, 0.15 mmol) and formaldehyde (37 wt. % in $H_2O$, 100 µL) was subjected to general reductive amination procedure with MeOH (2 mL), HOAc (50 µL) at room temperature, and then $NaBH_3CN$ (47 mg, 0.75 mmol) was added. After purification by Prep-HPLC, the product was applied to the general Aryl Nitro reduction procedure with ethanol/ saturated aq. $NH_4Cl$ (4 mL/1 mL) and Iron Powder (42 mg, 0.75 mmol). The purification by Prep-HPLC afforded the desired product **XDY2-58** (33 mg, yield 72%) as a light brown solid: $^1$H NMR (400 MHz, Chloroform-*d*) δ 7.73 (dt, *J* = 7.9, 1.0 Hz, 1H), 7.34 – 7.23 (m, 2H), 7.17 – 7.11 (m, 1H), 7.05 – 7.00 (m, 1H), 6.94 (d, *J* = 8.1 Hz, 1H), 6.64 (d, *J* = 2.6 Hz, 1H), 6.59 (dd, *J* = 8.1, 2.5 Hz, 1H), 5.92 (d, *J* = 8.2 Hz, 1H), 5.64 (tt, *J* = 7.4, 6.4 Hz, 1H), 3.77 (s, 3H), 2.32 (s, 3H), 1.73 (d, *J* = 6.7 Hz, 3H); LRMS (ESI) calcd for $C_{19}H_{22}N_3O$ $[M+H]^+$ 308.2, found 308.1.

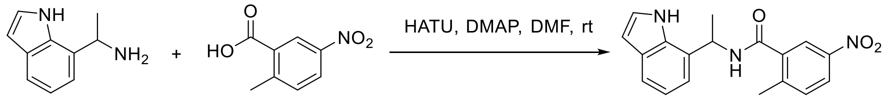


YF4-134

**N-(1-(1H-indol-7-yl)ethyl)-2-methyl-5-nitrobenzamide (YF4-134)**. 1-(1H-indol-7-yl)ethan-1-amine (100 mg, 0.62 mmol), 2-methyl-5-nitrobenzoic acid (101 mg, 0.56 mmol), HATU (213 mg, 0.56 mmol) and DMAP (205 mg, 1.68 mmol) was subjected to general amine coupling procedure with DMF (3 mL). The purification by Prep-HPLC afforded the **YF4-134** (142 mg, yield 78%) as a brown solid: $^1$H NMR (400 MHz, Methanol-*d*$_4$) δ 8.16 – 8.12 (m, 2H), 7.51 – 7.48 (m, 1H), 7.44 – 7.40 (m, 1H), 7.28 (d, *J* = 3.2 Hz, 1H), 7.18 – 7.15 (m, 1H), 7.05 – 7.00 (m, 1H), 6.49 (d, *J* = 3.2 Hz, 1H), 5.71 (q, *J* = 7.0 Hz, 1H), 2.37 (s, 3H), 1.70 (d, *J* = 7.0 Hz, 3H); $^{13}$C NMR (100 MHz, Methanol-*d*$_4$) δ 169.94, 147.22, 144.92, 139.02, 135.22, 132.90, 129.95, 127.31, 125.58, 125.21, 122.99, 120.69, 120.25, 118.45, 103.00, 46.58, 20.38, 19.70; HRMS (ESI) calcd for $C_{18}H_{19}N_2OS$ $[M+H]^+$ 311.1213, found 311.1215.

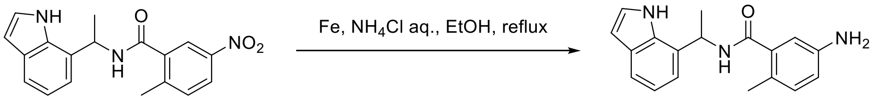


YF4-134 YF4-136

**N-(1-(1H-indol-7-yl)ethyl)-5-amino-2-methylbenzamide (YF4-136)**. To a solution of N-(1-(1H-indol-7-yl)ethyl)-2-methyl-5-nitrobenzamide (**YF4-134**) (80 mg, 0.25 mmol) in ethanol/ saturated aq. $NH_4Cl$ (4/1 mL), Fe powder (67 mg, 1.2 mmol) was added. The resulting solution was stirred for 2 h at 80°C and then concentrated under vacuum. The residue was extracted with 3×10 mL of ethyl acetate and the organic layers combined. The organic layer was washed with 5 mL of brine, dried and concentrated under vacuum to remove the solvent. The residue was purified by Prep-HPLC to obtain the desired product **YF4-136** (62 mg, yield 85%) as a light brown solid: $^1$H NMR (400 MHz, Methanol-*d*$_4$) δ 7.50 – 7.46 (m, 1H), 7.25 (d, *J* = 3.1 Hz, 1H), 7.18 – 7.14 (m, 1H), 7.05 – 6.98 (m, 1H), 6.94 – 6.91 (m, 1H), 6.66 (d, *J* = 7.2 Hz, 2H), 6.48 (d, *J* = 3.2 Hz, 1H), 5.67 (q, *J* = 7.0 Hz, 1H), 2.15 (s, 3H), 1.67 (d, *J* = 7.0 Hz, 3H); $^{13}$C NMR (100 MHz, Methanol-*d*$_4$) δ 173.02, 146.38, 138.36, 135.29, 132.29, 129.83, 127.59, 125.60, 125.44, 120.54, 120.19, 118.61, 118.02, 115.03, 102.98, 102.91, 46.29, 20.26, 18.49; HRMS (ESI) calcd for $C_{18}H_{20}N_3O$ $[M+H]^+$ 294.1601, found 294.1606.

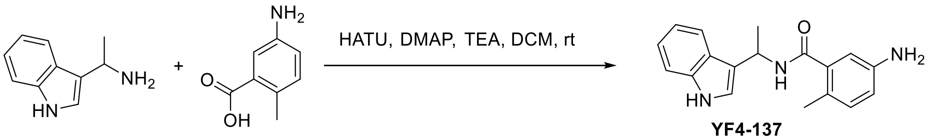


YF4-137

**N-(1-(1H-indol-3-yl)ethyl)-5-amino-2-methylbenzamide (YF4-137)**. 1-(1H-indol-3-yl)ethan-1-amine (75 mg, 0.47 mmol), 5-amino-2-methylbenzoic acid (47 mg, 0.31 mmol), HATU (118 mg, 0.31 mmol), TEA (100 µL), and DMAP (4 mg, 0.03 mmol) was subjected to general amine coupling procedure with DCM (2 mL). The purification by Prep-HPLC afforded the product **YF4-137** (55 mg, yield 60%) as a brown solid: $^{1}$H NMR (400 MHz, Methanol-$d_4$) δ 7.72 – 7.67 (m, 1H), 7.36 – 7.33 (m, 1H), 7.23 – 7.21 (m, 1H), 7.17 – 6.98 (m, 3H), 6.95 – 6.89 (m, 1H), 6.67 – 6.62 (m, 1H), 5.66 – 5.55 (m, 1H), 2.22 (s, 3H), 1.68 (d, *J* = 6.8 Hz, 3H); $^{13}$C NMR (100 MHz, Methanol-$d_4$) δ 172.38, 161.60, 146.27, 138.70, 138.35, 132.25, 125.56, 122.73, 122.62, 119.97, 119.93, 119.89, 117.95, 115.12, 112.34, 43.03, 20.85, 18.66; HRMS (ESI) calcd for $C_{18}H_{20}N_3O$ $[M+H]^+$ 294.1601, found 294.1649.

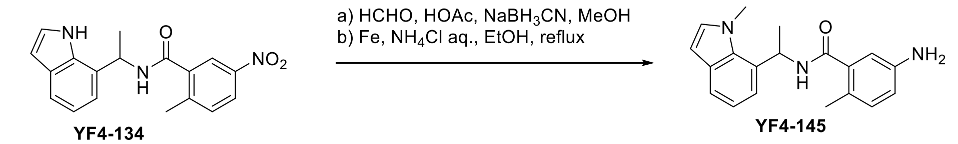


YF4-134 YF4-145

**5-amino-2-methyl-N-(1-(1-methyl-1H-indol-7-yl)ethyl)benzamide (YF4-145)**. N-(1-(1H-indol-7-yl)ethyl)-2-methyl-5-nitrobenzamide (50 mg, 0.15 mmol) and formaldehyde (37 wt. % in $H_2O$, 100 µL) was subjected to general reductive amination procedure with MeOH (2 mL), HOAc (50 µL) at room temperature, and then $NaBH_3CN$ (47 mg, 0.75 mmol) was added. After purification by Prep-HPLC, the product was dissolved in ethanol/ saturated aq. $NH_4Cl$ (4 mL/1 mL), and then Fe powder (42 mg, 0.75 mmol) was added. The resulting solution was stirred for 2 h at 80°C and then concentrated under vacuum. The residue was extracted with 3×10 mL of ethyl acetate and the organic layers were combined. The organic mixture was washed with brine, dried and concentrated under vacuum. The residue was purified by Prep-HPLC to obtain the desired product **YF4-145** (31 mg, yield 67%) as a brown solid: $^{1}$H NMR (400 MHz, Methanol-$d_4$) δ 7.58 – 7.51 (m, 1H), 7.33 (d, *J* = 7.4 Hz, 1H), 7.14 – 6.93 (m, 3H), 6.71 – 6.62 (m, 2H), 6.51 – 6.42 (m, 2H), 4.08 (s, 3H), 2.43 (s, 3H), 1.74 (d, *J* = 6.8 Hz, 3H); $^{13}$C NMR (100 MHz, Methanol-$d_4$) δ 174.63, 147.31, 137.70, 132.89, 132.32, 130.95, 125.77, 122.91, 122.38, 121.54, 120.09, 119.60, 117.35, 102.21, 46.05, 32.01, 17.89, 15.94; HRMS (ESI) calcd for $C_{19}H_{22}N_3O$ $[M+H]^+$ 308.1757, found 308.1709.

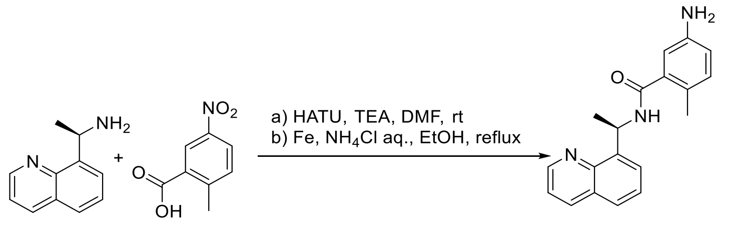


DY3-49

**(R)-5-amino-2-methyl-N-(1-(quinolin-8-yl)ethyl)benzamide (DY3-49)**. (R)-1-(quinolin-8-yl)ethan-1-amine (60 mg, 0.29 mmol), 2-methyl-5-nitrobenzoic acid (68 mg, 0.37 mmol), HATU (215 mg, 0.58 mmol), TEA (100 µL), and DMAP (5 mg, 0.04 mmol) was subjected to general amine coupling procedure with DMF (5 mL). After purification by Prep-HPLC, the product was applied to the general Aryl Nitro reduction procedure with ethanol/ saturated aq. $NH_4Cl$ (4 mL/1 mL) and Iron Powder (67 mg, 1.20 mmol). The purification by Prep-HPLC gave the **DY-3-49** (55 mg, 63% for 2 steps) as a white solid: $^{1}$H NMR (400 MHz, Methanol-d$_4$) $\delta$ 8.91 (dd, J = 4.1, 2.0 Hz, 1H), 8.37 – 8.22 (m, 1H), 7.91 (d, J = 2.4 Hz, 1H), 7.86 – 7.81 (m, 1H), 7.77 (d, J = 7.1 Hz,

1H), 7.62 – 7.55 (m, 1H), 7.50 (dt, J = 8.1, 4.0 Hz, 1H), 7.16 (d, J = 2.5 Hz, 1H), 6.24 (q, J = 7.9, 6.9 Hz, 1H), 2.38 (s, 3H), 1.66 (d, J = 7.0 Hz, 3H); $^{13}$C NMR (100 MHz, Methanol-*d*$_4$) $\delta$ 162.01, 150.68, 150.62, 146.69, 144.42, 143.26, 142.50, 141.69, 138.06, 136.20, 136.09, 130.14, 128.57, 127.54, 127.19, 122.74, 122.43, 49.00, 47.98, 22.64, 20.47; HRMS (ESI) calcd for $C_{19}H_{20}N_3O$ $[M+H]^+$ 306.1601, found 306.1600.

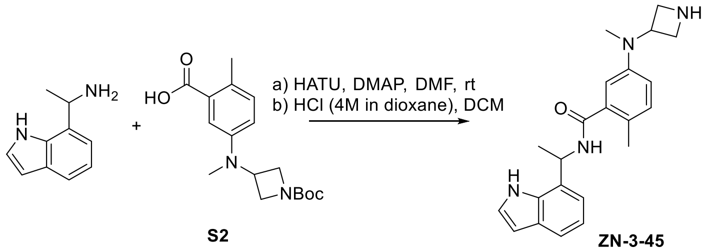


**N-(1-(1H-indol-7-yl)ethyl)-5-(azetidin-3-yl(methyl)amino)-2-methylbenzamide (ZN-3-45)**. 1-(1H-indol-7-yl)ethan-1-amine (24 mg, 0.15 mmol), 5-((1-(tert-butoxycarbonyl)azetidin-3-yl)(methyl)amino)-2-methylbenzoic acid (**S2**) (40 mg, 0.13 mmol), HATU (48 mg, 0.13 mmol) and DMAP (31 mg, 0.25 mmol) was subjected to general amine coupling procedure with DMF (1 mL). After purification by Prep-HPLC, the product was subjected to general N-Boc deprotection procedure with HCl (4M in dioxane, 100 µL) and DCM (2 mL). The purification by Prep-HPLC afforded the product **ZN-3-45** (33 mg, yield 70% for 2 steps) as a white solid: $^1$H NMR (400 MHz, Methanol-*d*$_4$) δ 8.38 (s, 1H), 7.49 (dd, *J* = 7.9, 1.1 Hz, 1H), 7.28 (d, *J* = 3.2 Hz, 1H), 7.14 (dd, *J* = 18.8, 7.8 Hz, 2H), 7.05 – 6.99 (m, 1H), 6.81 (dd, *J* = 8.3, 2.7 Hz, 1H), 6.72 (d, *J* = 2.7 Hz, 1H), 6.49 (d, *J* = 3.1 Hz, 1H), 5.69 (q, *J* = 6.9 Hz, 1H), 4.52 – 4.43 (m, 1H), 4.23 – 4.17 (m, 2H), 4.09 – 4.03 (m, 2H), 2.84 (s, 3H), 2.20 (s, 3H), 1.69 (d, *J* = 7.0 Hz, 3H); $^{13}$C NMR (100 MHz, Methanol-*d*$_4$) δ 172.48, 138.84, 135.24, 132.63, 129.91, 128.54, 127.61, 125.49, 120.59, 120.22, 119.76, 118.60, 116.90, 102.97, 53.34, 52.04, 46.50, 37.07, 20.40, 18.53; HRMS (ESI) calcd for $C_{22}H_{27}N_4O$ $[M+H]^+$ 363.2179, found 363.2179.

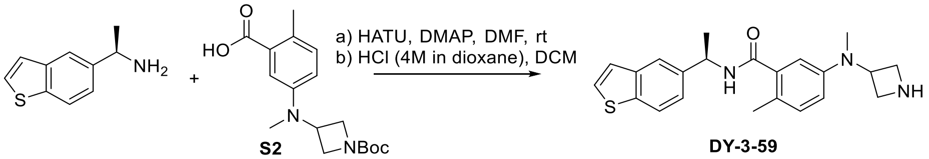


**(R)-5-(azetidin-3-yl(methyl)amino)-N-(1-(benzo[b]thiophen-5-yl)ethyl)-2-methylbenzamide (DY-3-59)**. (R)-1-(benzo[b]thiophen-5-yl)ethan-1-amine (21 mg, 0.12 mmol), 5-((1-(tert-butoxycarbonyl)azetidin-3-yl)(methyl)amino)-2-methylbenzoic acid (**S2**) (32 mg, 0.10 mmol), HATU (76 mg, 0.20 mmol) and DMAP (44 mg, 0.36 mmol) was subjected to general amine coupling procedure with DMF (1 mL). After purification by Prep-HPLC, the product was subjected to general N-Boc deprotection procedure with HCl (4M in dioxane, 100 µL) and DCM (2 mL). The purification by Prep-HPLC afforded the product **DY-3-59** (30 mg, yield 79% for 2 steps) as a white solid: $^1$H NMR (400 MHz, Methanol-*d*$_4$) $\delta$ 8.54 (s, 1H), 7.92 – 7.85 (m, 2H), 7.57 (d, J = 5.4 Hz, 1H), 7.45 – 7.39 (m, 1H), 7.36 (d, J = 5.4 Hz, 1H), 7.13 (d, J = 8.4 Hz, 1H), 6.81 (dd, J = 8.3, 2.6 Hz, 1H), 6.76 (d, J = 2.5 Hz, 1H), 5.33 (q, J = 7.0 Hz, 1H), 4.51 (p, J = 7.3 Hz, 1H), 4.22 (dd, J = 11.3, 7.6 Hz, 2H), 4.13 – 4.02 (m, 2H), 2.86 (s, 3H), 2.24 (s, 3H), 1.61 (s, 3H); $^{13}$C NMR (100 MHz, Methanol-*d*$_4$) $\delta$ 172.16, 148.44, 141.53, 141.35, 139.91, 138.96, 132.60, 128.45, 128.07, 124.88, 124.00, 123.48, 122.08, 119.65, 116.87, 53.35, 52.01, 50.66, 49.00, 37.03, 22.50, 18.64; HRMS (ESI) calcd for $C_{22}H_{26}N_3OS$ $[M+H]^+$ 380.1791, found 380.1798.

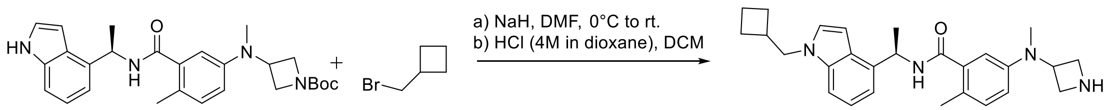


DY-3-66

**(R)-5-(azetidin-3-yl(methyl)amino)-N-(1-(1-(cyclobutylmethyl)-1H-indol-4-yl)ethyl)-2-methylbenzamide (DY-3-66)**. To a solution of tert-butyl (R)-3-((3-((1-(1H-indol-4-yl)ethyl)carbamoyl)-4-methylphenyl)(methyl)amino)azetidine-1-carboxylate (25 mg, 0.05 mmol) in dry DMF (2 mL), sodium hydride (4 mg, 0.08mmol) was added at 0 $^{o}$C. After 15 minutes, the mixture was added (bromomethyl)cyclobutane (9 mg, 0.06 mmol) and stirred at 25 $^{o}$C for another 16 hours. Quench the reaction with methanol and remove the solvent. The residue was purified by preparative HPLC system to obtain the desired product.

The product from previous step was subjected to general N-Boc deprotection procedure with HCl (4M in dioxane, 200 µL) and DCM (4 mL). The purification by Prep-HPLC afforded the product **DY-3-66** (17 mg, yield 80% for 2 steps) as a white solid: $^{1}$H NMR (400 MHz, Methanol-$d_4$) δ 8.53 (s, 1H), 7.33 (d, $J$ = 8.1 Hz, 1H), 7.25 (d, $J$ = 3.2 Hz, 1H), 7.18 – 7.07 (m, 3H), 6.79 (dd, $J$ = 8.3, 2.7 Hz, 1H), 6.73 (d, $J$ = 2.6 Hz, 1H), 6.66 (d, $J$ = 3.7 Hz, 1H), 5.62 (q, $J$ = 7.0 Hz, 1H), 4.46 (p, $J$ = 7.2 Hz, 1H), 4.19 (dd, $J$ = 12.3, 7.4 Hz, 4H), 4.05 (dd, $J$ = 10.3, 7.4 Hz, 2H), 2.83 (s, 3H), 2.23 (s, 3H), 2.06 – 1.96 (m, 2H), 1.93 – 1.79 (m, 4H), 1.65 (d, $J$ = 7.0 Hz, 3H). $^{13}$C NMR (100 MHz, Methanol-$d_4$) δ 148.36, 139.08, 137.99, 136.30, 132.56, 129.04, 128.62, 127.88, 122.25, 119.64, 117.05, 116.55, 111.39, 109.89, 99.99, 53.41, 52.28, 52.02, 49.00, 37.58, 37.12, 27.10, 21.24, 18.98, 18.65; HRMS (ESI) calcd for $C_{27}H_{35}N_4O$ $[M+H]^+$ 431.2805, found 431.2811.

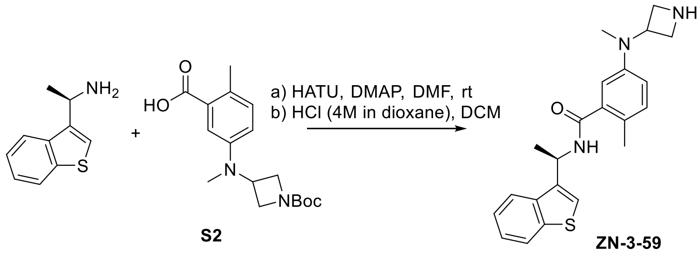


S2

ZN-3-59

**(R)-5-(azetidin-3-yl(methyl)amino)-N-(1-(benzo[b]thiophen-3-yl)ethyl)-2-methylbenzamide (ZN-3-59)**. (R)-1-(benzo[b]thiophen-3-yl)ethan-1-amine (30 mg, 0.14 mmol), 5-((1-(tert-butoxycarbonyl)azetidin-3-yl)(methyl)amino)-2-methylbenzoic acid (**S2**) (49 mg, 0.15 mmol), HATU (57 mg, 0.15 mmol) and DMAP (34 mg, 0.28 mmol) was subjected to general amine coupling procedure with DMF (1 mL). After purification by Prep-HPLC, the product was subjected to general N-Boc deprotection procedure with HCl (4M in dioxane, 100 µL) and DCM (2 mL). The purification by Prep-HPLC afforded the product **ZN-3-59** (45 mg, yield 85% for 2 steps) as a white solid: $[\alpha]^{25}_{546}$= -29.2 (c 0.5, MeOH); $^{1}$H NMR (400 MHz, Methanol-$d_4$) δ 8.41 (s, 1H), 7.98 (dt, $J$ = 8.1, 0.9 Hz, 1H), 7.94 – 7.86 (m, 1H), 7.53 (d, $J$ = 1.0 Hz, 1H), 7.47 – 7.33 (m, 2H), 7.11 (d, $J$ = 8.4 Hz, 1H), 6.80 (dd, $J$ = 8.4, 2.7 Hz, 1H), 6.70 (d, $J$ = 2.7 Hz, 1H), 5.73 – 5.63 (m, 1H), 4.46 (p, $J$ = 7.2 Hz, 1H), 4.18 (dd, $J$ = 10.9, 7.7 Hz, 2H), 4.05 (dd, $J$ = 11.1, 7.0 Hz, 2H), 2.82 (s, 3H), 2.24 (s, 3H), 1.72 (d, $J$ = 6.9 Hz, 3H); $^{13}$C NMR (100 MHz, Methanol-$d_4$) δ 172.08, 148.38, 142.09, 139.24, 139.19, 138.76, 132.63, 128.53, 125.62, 125.13, 123.91, 123.51, 123.09, 119.68, 116.92, 53.30, 51.96, 44.61, 37.00, 20.55, 18.67; HRMS (ESI) calcd for $C_{22}H_{26}N_3OS$ $[M+H]^+$ 380.1791, found 380.1791.

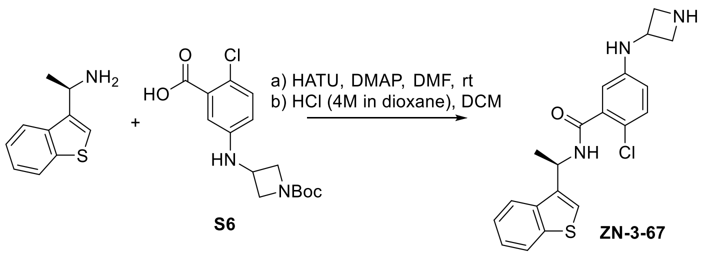


**(R)-5-(azetidin-3-ylamino)-N-(1-(benzo[b]thiophen-3-yl)ethyl)-2-chlorobenzamide (ZN-3-67)**. (R)-1-(benzo[b]thiophen-3-yl)ethan-1-amine (41 mg, 0.23 mmol), 5-((1-(tert-butoxycarbonyl)azetidin-3-yl)amino)-2-chlorobenzoic acid (**S6**) (50 mg, 0.15 mmol), HATU (70 mg, 0.18 mmol) and DMAP (56 mg, 0.46 mmol) was subjected to general amine coupling procedure with DMF (2 mL). After purification by Prep-HPLC, the product was subjected to general N-Boc deprotection procedure with HCl (4M in dioxane, 200 µL) and DCM (4 mL). The purification by Prep-HPLC afforded the product **ZN-3-67** (39 mg, yield 68% for 2 steps) as a white solid: $[\alpha]_{546}^{25}$ = -24.6 (c 1.7, MeOH); $^{1}$H NMR (400 MHz, Methanol-$d_4$) δ 8.49 (s, 1H), 8.00 – 7.95 (m, 1H), 7.90 – 7.85 (m, 1H), 7.54 (d, *J* = 1.1 Hz, 1H), 7.44 – 7.34 (m, 2H), 7.18 (d, *J* = 8.7 Hz, 1H), 6.60 (dd, *J* = 8.7, 2.8 Hz, 1H), 6.54 (d, *J* = 2.8 Hz, 1H), 5.68 – 5.61 (m, 1H), 4.48 – 4.40 (m, 1H), 4.36 – 4.27 (m, 2H), 3.95 – 3.86 (m, 2H), 1.71 (d, *J* = 6.9 Hz, 3H); $^{13}$C NMR (100 MHz, Methanol-$d_4$) δ 169.63, 146.53, 142.03, 139.13, 139.02, 138.17, 131.68, 125.58, 125.13, 123.80, 123.48, 123.17, 120.13, 116.28, 113.73, 54.66, 46.49, 44.88, 20.54; HRMS (ESI) calcd for $C_{20}H_{21}ClN_3OS$ $[M+H]^+$ 386.1088, found 386.1094.

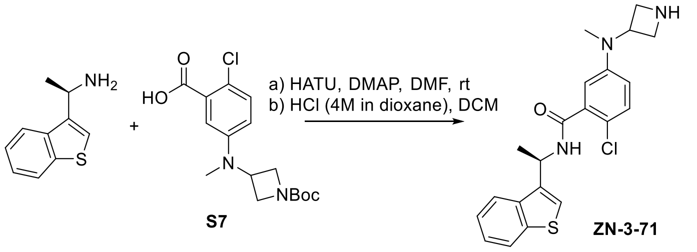


**(R)-5-(azetidin-3-yl(methyl)amino)-N-(1-(benzo[b]thiophen-3-yl)ethyl)-2-chlorobenzamide (ZN-3-71)**. (R)-1-(benzo[b]thiophen-3-yl)ethan-1-amine (41 mg, 0.23 mmol), 5-((1-(tert-butoxycarbonyl)azetidin-3-yl)(methyl)amino)-2-chlorobenzoic acid (**S7**) (51 mg, 0.15 mmol), HATU (70 mg, 0.18 mmol) and DMAP (56 mg, 0.46 mmol) was subjected to general amine coupling procedure with DMF (2 mL). After purification by Prep-HPLC, the product was subjected to general N-Boc deprotection procedure with HCl (4M in dioxane, 200 µL) and DCM (4 mL). The purification by Prep-HPLC afforded the **ZN-3-71** (44 mg, yield 74% for 2 steps) as a white solid: $[\alpha]_{546}^{25}$ = -24.7 (c 1.4, MeOH); $^{1}$H NMR (400 MHz, Methanol-$d_4$) δ 8.47 (s, 1H), 8.00 – 7.96 (m, 1H), 7.90 – 7.86 (m, 1H), 7.55 (d, *J* = 1.0 Hz, 1H), 7.44 – 7.34 (m, 2H), 7.28 (d, *J* = 8.8 Hz, 1H), 6.84 (dd, *J* = 8.8, 3.0 Hz, 1H), 6.75 (d, *J* = 3.0 Hz, 1H), 5.68 – 5.62 (m, 1H), 4.66 – 4.58 (m, 1H), 4.24 – 4.08 (m, 4H), 2.89 (s, 3H), 1.72 (d, *J* = 6.9 Hz, 3H); $^{13}$C NMR (100 MHz, Methanol-$d_4$) δ 169.30, 149.29, 142.06, 139.11, 138.98, 138.07, 131.64, 125.60, 125.11, 123.84, 123.55, 123.19, 122.23, 119.47, 116.93, 52.67, 51.92, 45.00, 35.61, 20.55; HRMS (ESI) calcd for $C_{21}H_{23}ClN_3OS$ $[M+H]^+$ 400.1245, found 400.1252.

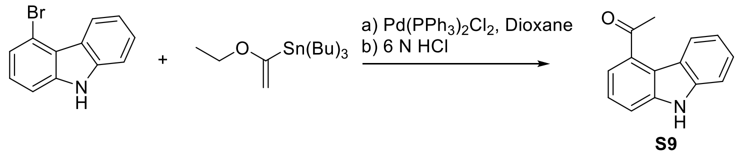


**1-(9H-carbazol-4-yl)ethan-1-one (S9)**. A mixture of the 4-bromo-9H-carbazole (738 mg, 3.0 mmol), $Pd(PPh_3)_2Cl_2$ (105 mg, 0.15 mmol) in anhyd dioxane (4 mL) was placed under argon, and tributyl(1-ethoxyvinyl)tin (1.1 mL, 3.3 mmol) was introduced by a syringe. After heating at 90 $^{o}$C for 6 h, 6 N aq HCl (10 mL) was added to quench the reaction. The solution was refluxed for 1 h, neutralized with $NaHCO_3$ and extracted with EtOAc (3×15 mL). The combined extracts were dried ($NaSO_4$) and concentrated under reduced pressure. To the residue was added sat. KF–MeOH (5 mL), and the mixture was filtered. After removal of the solvent, the residue was chromatographed on Silica Gel to provide the 1-(9H-carbazol-4-yl)ethan-1-one (502 mg, 80%) as a white solid: $^1$H NMR (400 MHz, $CDCl_3$) δ 8.81 – 8.73 (m, 1H), 8.50 (s, 1H), 7.66 (dd, *J* = 7.6, 1.0 Hz, 1H), 7.57 (dd, *J* = 8.1, 1.0 Hz, 1H), 7.48 – 7.37 (m, 3H), 7.25 – 7.20 (m, 1H), 2.80 (s, 3H); LRMS (ESI) calcd for $C_{14}H_{12}NO$ [M+H]$^+$ 210.1, found 210.1.

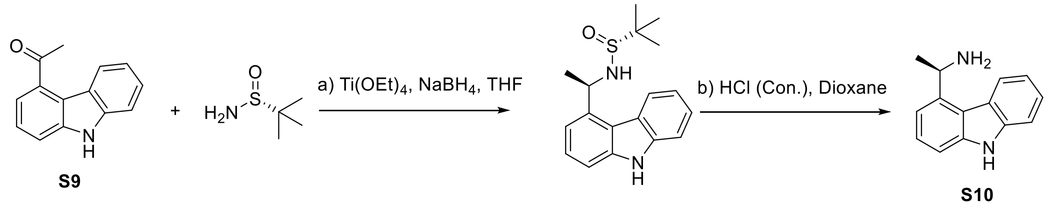


**(R)-1-(9H-carbazol-4-yl)ethan-1-amine (S10)**. To a solution of $Ti(OEt)_4$ (0.3 mL, 1.44 mmol) and 1-(9H-carbazol-4-yl)ethan-1-one (**S9**) (183 mg, 0.87 mmol) in THF (5 mL) under an Ar atmosphere was added (R)-2-methylpropane-2-sulfinamide (127 mg, 1.04 mmol) and the mixture was heated (70 °C). Upon completion, as determined by TLC, the mixture was cooled to −78 °C and $NaBH_4$ (109 mg. 2.88 mmol) was added carefully. The mixture was stirred at −78 °C for 3 h, and then warmed up to room temperature slowly. After another 3 h, MeOH was added dropwise until gas was no longer evolved. The resulting suspension was filtered through a plug of Celite and the filter cake was washed with EtOAc. The filtrate was washed with brine, and the brine layer was extracted with EtOAc. The combined organic portions were dried ($Na_2SO_4$), filtered, and concentrated. Use silica gel column chromatography (Hexanes/EtOAc) to get the major product (178 mg, yield 65%) as a syrup: $^1$H NMR (400 MHz, $CDCl_3$) δ 8.25 (d, *J* = 8.1 Hz, 1H), 7.44 – 7.15 (m, 6H), 4.14 (q, *J* = 7.1 Hz, 1H), 1.78 (d, *J* = 7.1 Hz, 3H), 1.26 (s, 9H); LRMS (ESI) calcd for $C_{18}H_{23}N_2OS$ [M+H]$^+$ 315.2, found 315.1.

The product from the previous step was soluble in dioxane (5 mL) and then concentrated HCl aq. (100 µL) was added. The mixture was stirred at room temperature for 10 min. After the filtration, the compound was generated as a wight solid: LRMS (ESI) calcd for $C_{14}H_{15}N_2$ [M+H]$^+$ 211.1, found 211.1.

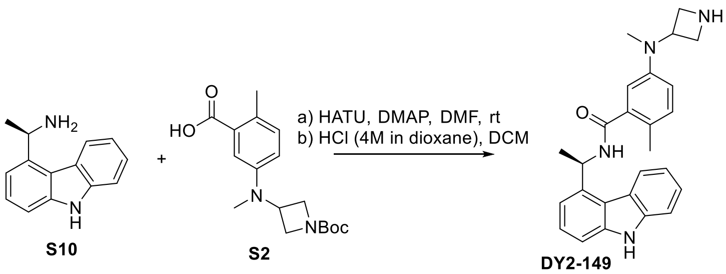


**(R)-N-(1-(9H-carbazol-4-yl)ethyl)-5-(azetidin-3-yl(methyl)amino)-2-methylbenzamide (DY2-149)**. (R)-1-(9H-carbazol-4-yl)ethan-1-amine (32 mg, 0.15 mmol), 5-((1-(tert-butoxycarbonyl)azetidin-3-yl)(methyl)amino)-2-methylbenzoic acid (**S2**) (48 mg, 0.15 mmol), HATU (70 mg, 0.18 mmol) and DMAP (56 mg, 0.46 mmol) was subjected to general amine coupling procedure with DMF (2 mL). After purification by Prep-HPLC, the product was subjected to general N-Boc deprotection procedure with HCl (4M in dioxane, 200 µL) and DCM (4 mL). The purification by Prep-HPLC afforded the product **DY2-149** (53 mg, yield 86% for 2 steps) as a white solid: $[\alpha]_{546}^{25}$= -42.7 (c 0.8, MeOH); $^{1}$H NMR (400 MHz, Methanol-$d_4$) δ 8.37 (s, 1H), 8.24 (d, *J* = 8.1 Hz, 1H), 7.51 (d, *J* = 8.2 Hz, 1H), 7.44 – 7.37 (m, 3H), 7.30 – 7.27 (m, 1H), 7.23 – 7.18 (m, 1H), 7.12 (d, *J* = 8.3 Hz, 1H), 6.79 (dd, *J* = 8.3, 2.8 Hz, 1H), 6.66 (d, *J* = 2.8 Hz, 1H), 6.14 (q, *J* = 6.8 Hz, 1H), 4.39 – 4.30 (m, 1H), 4.12 – 3.94 (m, 4H), 2.77 (s, 3H), 2.30 (s, 3H), 1.79 (d, *J* = 6.8 Hz, 3H); $^{13}$C NMR (100 MHz, Methanol-$d_4$) δ 172.53, 148.29, 141.94, 141.69, 138.86, 132.61, 128.78, 126.54, 126.29, 123.87, 123.43, 121.45, 119.94, 119.83, 117.10, 116.24, 111.85, 111.10, 53.35, 51.94, 48.25, 37.12, 20.53, 18.69; HRMS (ESI) calcd for $C_{26}H_{29}N_4O$ $[M+H]^+$ 413.2336, found 413.2341.

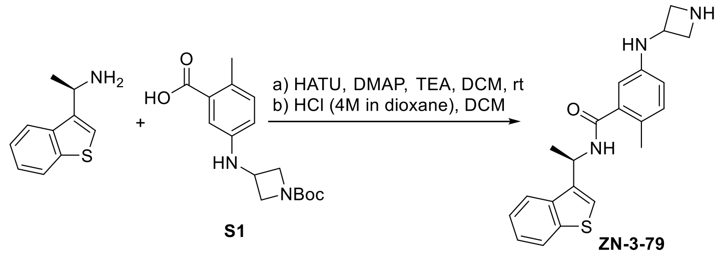


**(R)-5-(azetidin-3-ylamino)-N-(1-(benzo[b]thiophen-3-yl)ethyl)-2-methylbenzamide (ZN-3-79)**. (R)-1-(benzo[b]thiophen-3-yl)ethan-1-amine (41 mg, 0.23 mmol), 5-((1-(tert-butoxycarbonyl)azetidin-3-yl)amino)-2-methylbenzoic acid (**S1**) (46 mg, 0.15 mmol), HATU (70 mg, 0.18 mmol) and DMAP (56 mg, 0.46 mmol) was subjected to general amine coupling procedure with DMF (2 mL). After purification by Prep-HPLC, the product was subjected to general N-Boc deprotection procedure with HCl (4M in dioxane, 200 µL) and DCM (4 mL). The purification by Prep-HPLC afforded the product **ZN-3-79** (46 mg, yield 84% for 2 steps) as a white solid: $[\alpha]_{546}^{25}$= -31.7 (c 2.0, MeOH); $^{1}$H NMR (400 MHz, Methanol-$d_4$) δ 8.48 (s, 1H), 7.99 – 7.94 (m, 1H), 7.90 – 7.85 (m, 1H), 7.52 (s, 1H), 7.44 – 7.33 (m, 2H), 7.00 (d, *J* = 8.3 Hz, 1H), 6.54 (dd, *J* = 8.3, 2.6 Hz, 1H), 6.48 (d, *J* = 2.6 Hz, 1H), 5.66 (q, *J* = 6.9 Hz, 1H), 4.42 (p, *J* = 7.0 Hz, 1H), 4.31 – 4.23 (m, 2H), 3.93 – 3.84 (m, 2H), 2.21 (s, 3H), 1.70 (d, *J* = 6.9 Hz, 3H); $^{13}$C NMR (100 MHz, Methanol-$d_4$) δ 172.33, 145.38, 142.04, 139.31, 139.19, 138.71, 132.62, 125.76, 125.59, 125.13, 123.86, 123.41, 123.07, 115.58, 112.80, 54.76, 46.79, 44.50, 20.58, 18.61; HRMS (ESI) calcd for $C_{21}H_{24}N_3OS$ $[M+H]^+$ 366.1635, found 366.1640.

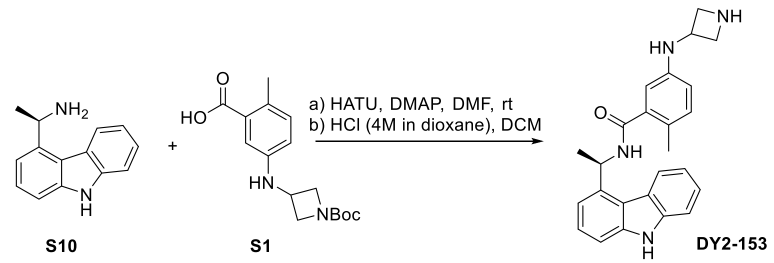


**(R)-N-(1-(9H-carbazol-4-yl)ethyl)-5-(azetidin-3-ylamino)-2-methylbenzamide (DY2-153)**. (R)-1-(9H-carbazol-4-yl)ethan-1-amine (32 mg, 0.15 mmol), 5-((1-(tert-butoxycarbonyl)azetidin-3-yl)amino)-2-methylbenzoic acid (**S1**) (46 mg, 0.15 mmol), HATU (70 mg, 0.18 mmol) and DMAP (56 mg, 0.46 mmol) was subjected to general amine coupling procedure with DMF (2 mL). After purification by Prep-HPLC, the product was subjected to general N-Boc deprotection procedure with HCl (4M in dioxane, 200 µL) and DCM (4 mL). The purification by Prep-HPLC afforded the product **DY2-153** (47 mg, yield 78% for 2 steps) as a white solid: $^1$H NMR (400 MHz, Methanol-$d_4$) δ 8.53 (s, 2H), 8.24 (d, *J* = 8.2 Hz, 1H), 7.53 – 7.48 (m, 1H), 7.44 – 7.36 (m, 3H), 7.30 – 7.26 (m, 1H), 7.23 – 7.17 (m, 1H), 7.02 (d, *J* = 8.2 Hz, 1H), 6.55 (dd, *J* = 8.4, 2.8 Hz, 1H), 6.48 – 6.43 (m, 1H), 6.15 (q, *J* = 6.5 Hz, 1H), 4.39 – 4.30 (m, 1H), 4.25 – 4.15 (m, 2H), 3.88 – 3.80 (m, 2H), 2.27 (s, 3H), 1.77 (d, *J* = 6.8 Hz, 3H); $^{13}$C NMR (100 MHz, Methanol-$d_4$) δ 172.78, 145.33, 141.93, 141.70, 139.03, 138.91, 132.62, 126.52, 126.28, 126.07, 123.89, 123.46, 121.44, 119.96, 116.18, 115.84, 112.78, 111.81, 111.02, 54.83, 48.14, 46.97, 20.68, 18.62; HRMS (ESI) calcd for $C_{25}H_{27}N_4O$ $[M+H]^+$ 399.2179, found 399.2176.

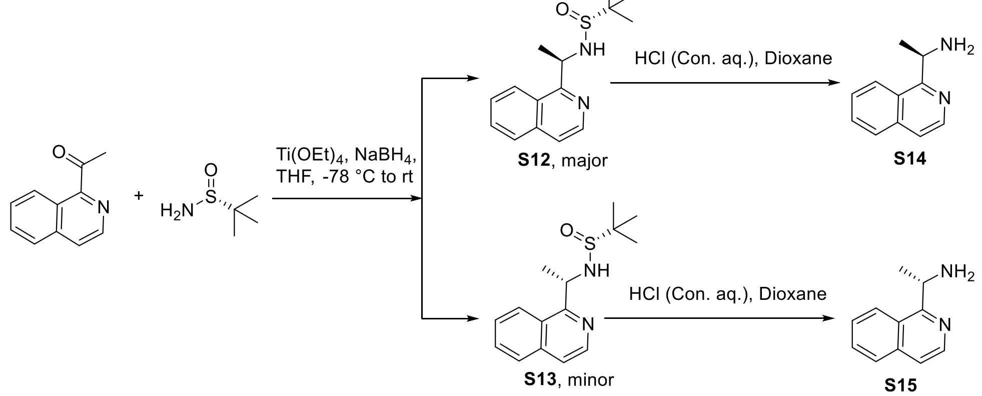


**(R)-N-((R/S)-1-(isoquinolin-1-yl)ethyl)-2-methylpropane-2-sulfinamide**[2]. To a solution of $Ti(OEt)_4$ (0.3 mL, 1.44 mmol) and 1-(isoquinolin-1-yl)ethan-1-one (171 mg, 1.00 mmol) in THF (5 mL) under an Ar atmosphere was added (R)-2-methylpropane-2-sulfinamide (182 mg, 1.50 mmol) and the mixture was heated (70 °C). Upon completion, as determined by TLC, the mixture was cooled to −78 °C and $NaBH_4$ (113 mg. 3.00 mmol) was added carefully. The mixture was stirred at −78 °C for 3 h, and then warmed up to room temperature slowly. After another 3 h, MeOH was added dropwise until gas was no longer evolved. The resulting suspension was filtered through a plug of Celite and the filter cake was washed with EtOAc. The filtrate was washed with brine, and the brine layer was extracted with EtOAc. The combined organic portions were dried ($Na_2SO_4$), filtered, and concentrated. Use silica gel column chromatography (Hexanes/EtOAc) to get the products (**S12**, 199 mg, yield 72%; **S13**, 54 mg, yield 20%) as colorless syrup:

**S12**: $^{1}$H NMR (400 MHz, Chloroform-*d*) δ 8.40 (d, *J* = 5.7 Hz, 1H), 8.08 (dd, *J* = 8.6, 1.3 Hz, 1H), 7.80 – 7.73 (m, 1H), 7.65 – 7.53 (m, 2H), 7.51 – 7.48 (m, 1H), 5.64 (d, *J* = 5.7 Hz, 1H), 5.38 (dt, *J* = 12.5, 6.6 Hz, 1H), 1.53 (d, *J* = 6.7 Hz, 3H), 1.28 (s, 9H); LRMS (ESI) calcd for $C_{15}H_{21}N_2OS$ $[M+H]^+$ 227.14, found 227.11.

**S13**: $^{1}$H NMR (400 MHz, Chloroform-*d*) δ 8.45 (d, *J* = 5.7 Hz, 1H), 8.12 (d, *J* = 8.4 Hz, 1H), 7.82 (d, *J* = 8.1 Hz, 1H), 7.69 – 7.58 (m, 2H), 7.55 (d, *J* = 5.7 Hz, 1H), 5.41 (dq, *J* = 9.0, 6.7 Hz, 1H), 4.88 (d, *J* = 9.0 Hz, 1H), 1.73 (d, *J* = 6.7 Hz, 3H), 1.18 (s, 9H); LRMS (ESI) calcd for $C_{15}H_{21}N_2OS$ $[M+H]^+$ 227.14, found 227.15.

**(R/S)-1-(isoquinolin-1-yl)ethan-1-amine**. **S12** and **S13** were soluble separately in Dioxane (5 mL) and then concentrated HCl aq. (100 µL) was added. The reactions were stirred at room temperature for 10 min. After the filtration and washed with cold EtOAc, the compounds were generated as wight solids.

**S14**: $^{1}$H NMR (400 MHz, Chloroform-*d*) δ 8.47 (d, *J* = 5.7 Hz, 1H), 8.21 – 8.13 (m, 1H), 7.88 – 7.80 (m, 1H), 7.70 – 7.59 (m, 2H), 7.55 (d, *J* = 5.7 Hz, 1H), 5.03 (q, *J* = 6.7 Hz, 1H), 1.56 (d, *J* = 6.7 Hz, 3H); LRMS (ESI) calcd for $C_{11}H_{13}N_2$ $[M+H]^+$ 173.1, found 173.1.

**S15**: $^{1}$H NMR (400 MHz, Chloroform-*d*) δ 8.47 (d, *J* = 5.6 Hz, 1H), 8.22 – 8.12 (m, 1H), 7.90 – 7.80 (m, 1H), 7.70 – 7.58 (m, 2H), 7.55 (d, *J* = 5.7 Hz, 1H), 5.01 (q, *J* = 6.7 Hz, 1H), 1.56 (d, *J* = 6.7 Hz, 3H); LRMS (ESI) calcd for $C_{11}H_{13}N_2$ $[M+H]^+$ 173.1, found 173.1.

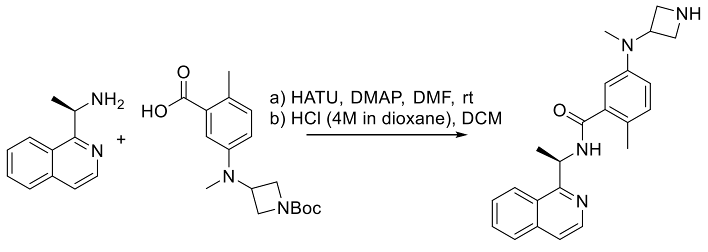


S2 ZN-3-36

**(R)-5-(azetidin-3-yl(methyl)amino)-N-(1-(isoquinolin-1-yl)ethyl)-2-methylbenzamide (ZN-3-36)**. (R)-1-(isoquinolin-1-yl)ethan-1-amine (35 mg, 0.20 mmol), 5-((1-(tert-butoxycarbonyl)azetidin-3-yl)(methyl)amino)-2-methylbenzoic acid (**S2**) (96 mg, 0.30 mmol), HATU (114 mg, 0.30 mmol) and DMAP (110 mg, 0.90 mmol) was subjected to general amine coupling procedure with DMF (1.5 mL). After purification by Prep-HPLC, the product was subjected to general N-Boc deprotection procedure with HCl (4M in dioxane, 200 µL) and DCM (4 mL). The purification by Prep-HPLC afforded the product **ZN-3-36** (63 mg, yield 84% for 2 steps) as a white solid: $[\alpha]_{546}^{25}$ = -49.6 (c 0.8, MeOH); $^{1}$H NMR (400 MHz, Methanol-$d_4$) δ 8.54 (s, 1H), 8.47 – 8.40 (m, 2H), 8.01 – 7.94 (m, 1H), 7.82 – 7.71 (m, 3H), 7.14 (d, *J* = 8.0 Hz, 1H), 6.88 – 6.81 (m, 2H), 6.17 (q, *J* = 6.9 Hz, 1H), 4.56 – 4.46 (m, 1H), 4.26 – 4.19 (m, 2H), 4.11 – 4.04 (m, 2H), 2.87 (s, 3H), 2.28 (s, 3H), 1.69 (d, *J* = 6.8 Hz, 3H); $^{13}$C NMR (100 MHz, Methanol-$d_4$) δ 171.96, 161.77, 148.51, 142.19, 138.47, 138.16, 132.77, 131.71, 128.97, 128.75, 126.89, 125.55, 121.88, 119.91, 117.01, 53.46, 52.13, 48.09, 37.09, 21.35, 18.78; HRMS (ESI) calcd for $C_{23}H_{27}N_4O$ $[M+H]^+$ 375.2179, found 375.2183.

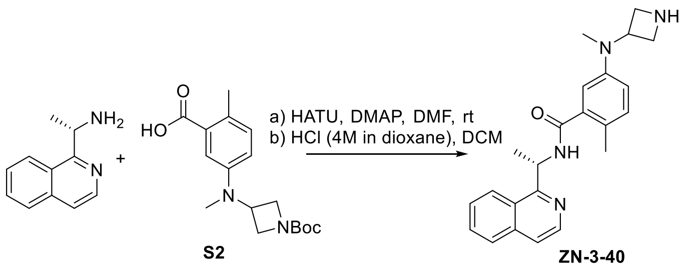


**(S)-5-(azetidin-3-yl(methyl)amino)-N-(1-(isoquinolin-1-yl)ethyl)-2-methylbenzamide (ZN-3-40)**. (S)-1-(isoquinolin-1-yl)ethan-1-amine (10 mg, 0.06 mmol), 5-((1-(tert-butoxycarbonyl)azetidin-3-yl)(methyl)amino)-2-methylbenzoic acid (**S2**) (29 mg, 0.09 mmol), HATU (34 mg, 0.09 mmol) and DMAP (33 mg, 0.27 mmol) was subjected to general amine coupling procedure with DMF (1 mL). After purification by Prep-HPLC, the product was subjected to general N-Boc deprotection procedure with HCl (4M in dioxane, 100 µL) and DCM (2 mL). The purification by Prep-HPLC afforded the product **ZN-2-40** (16 mg, yield 71% for 2 steps) as a white solid: $[\alpha]_{546}^{25}$= +40.7 (c 0.4, MeOH);$^1$H NMR (400 MHz, Methanol-$d_4$) δ 8.51 (s, 1H), 8.44 (t, $J$ = 7.5 Hz, 2H), 7.99 – 7.95 (m, 1H), 7.83 – 7.71 (m, 3H), 7.13 (dd, $J$ = 10.6, 8.1 Hz, 1H), 6.87 – 6.81 (m, 2H), 6.20 – 6.12 (m, 1H), 4.56 – 4.46 (m, 1H), 4.29 – 4.21 (m, 2H), 4.08 (dd, $J$ = 11.2, 7.1 Hz, 2H), 2.87 (s, 3H), 2.28 (s, 3H), 1.69 (d, $J$ = 6.9 Hz, 3H); $^{13}$C NMR (100 MHz, Methanol-$d_4$) 171.90, 161.77, 148.51, 142.18, 138.47, 138.16, 132.78, 131.72, 128.98, 128.76, 125.55, 121.89, 119.96, 117.06, 53.40, 52.13, 48.08, 37.13, 21.36, 18.78; HRMS (ESI) calcd for $C_{23}H_{27}N_4O$ [M+H]$^+$ 375.2179, found 375.2185.

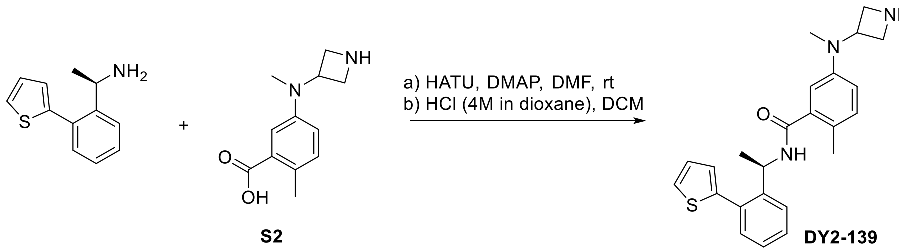


**(R)-5-(azetidin-3-yl(methyl)amino)-2-methyl-N-(1-(2-(thiophen-2-yl)phenyl)ethyl)benzamide (DY2-139)**. (R)-1-(2-(thiophen-2-yl)phenyl)ethan-1-amine (30 mg, 0.15 mmol), 5-(azetidin-3-yl(methyl)amino)-2-methylbenzoic acid (**S2**) (33 mg, 0.15 mmol), HATU (70 mg, 0.18 mmol) and DMAP (56 mg, 0.46 mmol) was subjected to general amine coupling procedure with DMF (2 mL). After purification by Prep-HPLC, the product was subjected to general N-Boc deprotection procedure with HCl (4M in dioxane, 100 µL) and DCM (2 mL). The purification by Prep-HPLC afforded the **DY2-139** (50 mg, yield 82% for 2 steps) as a white solid: $^1$H NMR (400 MHz, Acetone-$d_6$) δ 7.79 – 7.71 (m, 1H), 7.55 (dd, $J$ = 5.2, 1.2 Hz, 1H), 7.46 – 7.27 (m, 4H), 7.18 (dd, $J$ = 5.2, 3.4 Hz, 1H), 7.02 (d, $J$ = 8.4 Hz, 1H), 6.77 (d, $J$ = 2.7 Hz, 1H), 6.72 (dd, $J$ = 8.3, 2.8 Hz, 1H), 5.65 – 5.55 (m, 1H), 3.99 (p, $J$ = 6.7 Hz, 1H), 3.65 – 3.55 (m, 2H), 3.39 – 3.24 (m, 2H), 2.81 (s, 3H), 2.21 (s, 3H), 1.43 (d, $J$ = 7.0 Hz, 3H); $^{13}$C NMR (100 MHz, Acetone-$d_6$) δ 162.86, 148.98, 144.94, 142.60, 138.59, 133.94, 131.91, 129.41, 128.15, 128.06, 127.52, 126.81, 126.64, 126.27, 117.24, 115.29, 52.57, 52.52, 50.31, 46.80, 36.49, 23.20, 18.79; HRMS (ESI) calcd for $C_{24}H_{28}N_3OS$ [M+H]$^+$ 406.1948, found 406.1953

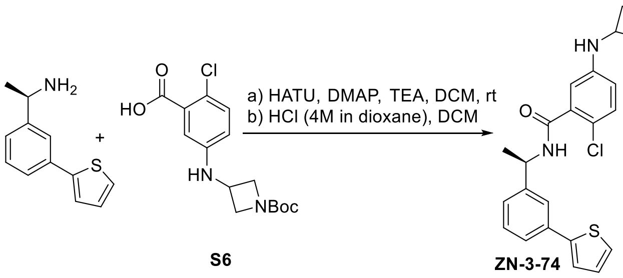


S6 ZN-3-74

**(R)-5-(azetidin-3-ylamino)-2-chloro-N-(1-(3-(thiophen-2-yl)phenyl)ethyl)benzamide (ZN-3-74)**. (R)-1-(3-(thiophen-2-yl)phenyl)ethan-1-amine (30 mg, 0.15 mmol), 5-((1-(tert-butoxycarbonyl)azetidin-3-yl)amino)-2-chlorobenzoic acid (**S6**) (50 mg, 0.15 mmol), HATU (70 mg, 0.18 mmol) and DMAP (56 mg, 0.46 mmol) was subjected to general amine coupling procedure with DMF (2 mL). After purification by Prep-HPLC, the product was subjected to general N-Boc deprotection procedure with HCl (4M in dioxane, 200 µL) and DCM (4 mL). The purification by Prep-HPLC afforded the **ZN-3-74** (48 mg, yield 77% for 2 steps) as a white solid: $[\alpha]_{546}^{25}$ = +58.5 (c 1.5, MeOH); $^1$H NMR (400 MHz, Methanol-$d_4$) δ 8.42 (s, 1H), 7.73 – 7.70 (m, 1H), 7.55 – 7.51 (m, 1H), 7.41 – 7.32 (m, 4H), 7.20 (d, *J* = 8.5 Hz, 1H), 7.08 (dd, *J* = 5.1, 3.6 Hz, 1H), 6.64 – 6.58 (m, 2H), 5.21 (q, *J* = 7.0 Hz, 1H), 4.51 – 4.43 (m, 1H), 4.36 – 4.30 (m, 2H), 3.95 – 3.89 (m, 2H), 1.56 (d, *J* = 7.1 Hz, 3H); $^{13}$C NMR (100 MHz, Methanol-$d_4$) δ 169.47, 146.59, 145.89, 145.40, 138.26, 135.99, 131.68, 130.17, 129.10, 126.43, 125.91, 125.52, 124.65, 124.35, 120.11, 116.39, 113.71, 54.62, 50.74, 46.50, 22.41; HRMS (ESI) calcd for $C_{22}H_{23}ClN_3OS$ [M+H]$^+$ 412.1245, found 412.1250.

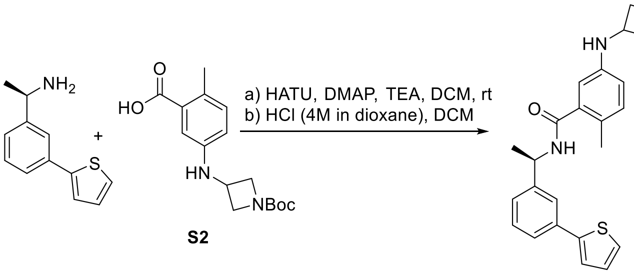


S2 ZN-3-80

**(R)-5-(azetidin-3-ylamino)-2-methyl-N-(1-(3-(thiophen-2-yl)phenyl)ethyl)benzamide (ZN-3-80)**. (R)-1-(3-(thiophen-2-yl)phenyl)ethan-1-amine (30 mg, 0.15 mmol), 5-((1-(tert-butoxycarbonyl)azetidin-3-yl)amino)-2-methylbenzoic acid (**S2**) (46 mg, 0.15 mmol), HATU (70 mg, 0.18 mmol) and DMAP (56 mg, 0.46 mmol) was subjected to general amine coupling procedure with DMF (2 mL). After purification by Prep-HPLC, the product was subjected to general N-Boc deprotection procedure with HCl (4M in dioxane, 200 µL) and DCM (4 mL). The purification by Prep-HPLC afforded the product **ZN-3-80** (47 mg, yield 80% for 2 steps) as a white solid: $[\alpha]_{546}^{25}$ = +25.2 (c 0.8, MeOH); $^1$H NMR (400 MHz, Methanol-$d_4$) δ 8.51 (s, 1H), 7.70 – 7.68 (m, 1H), 7.56 – 7.51 (m, 1H), 7.41 – 7.31 (m, 4H), 7.09 (dd, *J* = 5.1, 3.6 Hz, 1H), 7.03 (d, *J* = 8.2 Hz, 1H), 6.60 – 6.52 (m, 2H), 5.22 (q, *J* = 7.0 Hz, 1H), 4.48 (p, *J* = 7.0 Hz, 1H), 4.38 – 4.29 (m, 2H), 3.95 – 3.88 (m, 2H), 2.22 (s, 3H), 1.55 (d, *J* = 7.1 Hz, 3H); $^{13}$C NMR (100 MHz, Methanol-$d_4$) δ 172.56, 146.28, 145.45, 145.41, 138.93, 136.04, 132.62, 130.23, 129.12, 126.38, 125.93, 125.71, 125.51, 124.62, 124.34, 115.64, 112.68, 54.85, 50.51, 46.86, 22.39, 18.62; HRMS (ESI) calcd for $C_{23}H_{26}N_3OS$ [M+H]$^+$ 392.1791, found 392.1790.

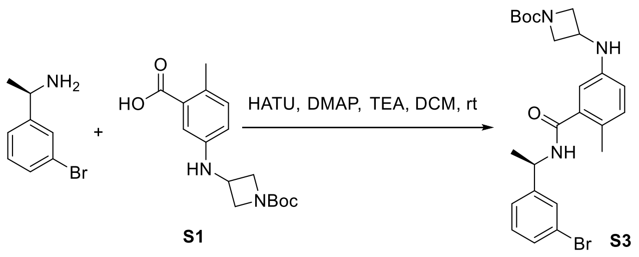


**Tert-butyl (R)-3-((3-((1-(3-bromophenyl)ethyl)carbamoyl)-4-methylphenyl)amino)azetidine-1-carboxylate (S3)**. (R)-1-(3-bromophenyl)ethan-1-amine (200 mg, 1.00 mmol), 5-((1-(tert-butoxycarbonyl)azetidin-3-yl)amino)-2-methylbenzoic acid (**S1**) (306 mg, 1.00 mmol), HATU (380 mg, 1.00 mmol), DMAP (12 mg, 0.10 mmol), and TEA (200 µL) was subjected to general amine coupling procedure with DMF (5 mL). The purification by Prep-HPLC afforded the **S3** (435 mg, yield 89%) as a white solid: $^{1}$H NMR (400 MHz, Chloroform-*d*) δ 7.51 – 7.50 (m, 1H), 7.40 (ddd, *J* = 7.8, 2.0, 1.2 Hz, 1H), 7.32 – 7.28 (m, 1H), 7.24 – 7.19 (m, 1H), 7.01 (d, *J* = 8.2 Hz, 1H), 6.52 (d, *J* = 2.6 Hz, 1H), 6.46 (dd, *J* = 8.2, 2.6 Hz, 1H), 6.02 (d, *J* = 8.0 Hz, 1H), 5.25 (p, *J* = 7.1 Hz, 1H), 4.26 (dd, *J* = 8.7, 7.1 Hz, 2H), 4.20 – 4.13 (m, 1H), 3.68 (dd, *J* = 9.0, 4.6 Hz, 2H), 2.28 (s, 3H), 1.55 (d, *J* = 7.0 Hz, 3H), 1.43 (s, 9H); $^{13}$C NMR (100 MHz, Chloroform-*d*) δ 169.40, 156.30, 145.71, 144.24, 137.28, 132.17, 130.63, 130.45, 129.30, 125.28, 125.09, 122.93, 114.74, 111.81, 79.88, 48.75, 43.41, 38.74, 28.50, 21.99, 18.83; LRMS (ESI) calcd for $C_{24}H_{31}BrN_3O_3$ $[M+H]^+$ 488.2, found 488.2.

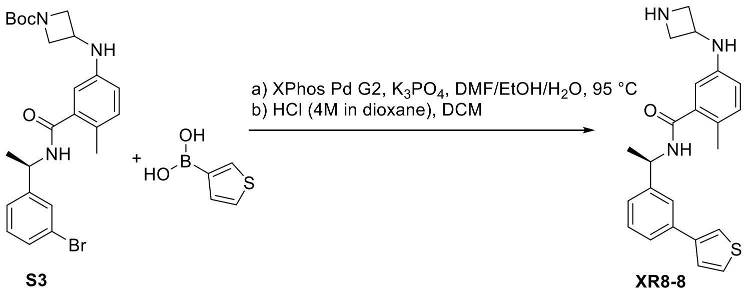


**(R)-5-(azetidin-3-ylamino)-2-methyl-N-(1-(3-(thiophen-3-yl)phenyl)ethyl)benzamide (XR8-8)**. A flask fitted with a rubber septum was charged with tert-butyl (R)-3-((3-((1-(3-bromophenyl)ethyl)carbamoyl)-4-methylphenyl)amino)azetidine-1-carboxylate (**S3**) (49 mg, 0.10 mmol), thiophen-3-ylboronic acid (19 mg, 0.15 mmol), XPhos Pd G2 (8 mg, 0.01 mmol), $K_3PO_4$ (64 mg, 0.3 mmol), DMF/EtOH/$H_2O$ (1 mL/ 1 mL/ 0.5 mL) and then purged with argon. The mixture was stirred at 95 $^{o}$C overnight. The reaction mixture was then cooled to room temperature, diluted with ethyl acetate (20 mL), filtered through celite and concentrated in vacuo. After purification by Prep-HPLC, the product was subjected to general N-Boc deprotection procedure with HCl (4M in dioxane, 100 µL) and DCM (2 mL). The purification by Prep-HPLC afforded the product **XR8-8** (14 mg, yield 35% for 2 steps) as a white solid: $[\alpha]^{25}_{546}$= +12.3 (c 0.6, MeOH); $^{1}$H NMR (400 MHz, Methanol-*d*$_4$) δ 8.41 (s, 1H), 7.72 – 7.69 (m, 1H), 7.63 – 7.60 (m, 1H), 7.58 – 7.53 (m, 1H), 7.50 – 7.45 (m, 2H), 7.41 – 7.31 (m, 2H), 7.02 (d, *J* = 8.2 Hz, 1H), 6.61 – 6.50 (m, 2H), 5.23 (q, *J* = 7.0 Hz, 1H), 4.52 – 4.42 (m, 1H), 4.35 – 4.29 (m, 2H), 3.95 – 3.88 (m, 2H), 2.21 (s, 3H), 1.56 (d, *J* = 7.0 Hz, 3H); $^{13}$C NMR (100 MHz, Methanol-*d*$_4$) δ 172.53, 146.00, 145.45, 143.51, 138.97, 137.47, 132.63, 130.10, 127.36, 127.17, 126.13, 125.92, 125.70, 125.29, 121.38,

115.58, 112.72, 54.84, 50.62, 46.84, 22.47, 18.57; HRMS (ESI) calcd for $C_{23}H_{26}N_3OS$ $[M+H]^+$ 392.1791, found 392.1800.

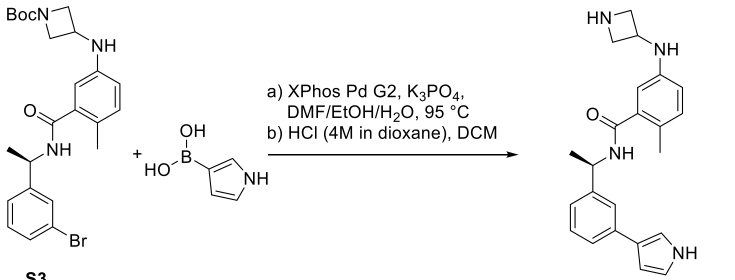


**S3** **XR8-9**

**(R)-N-(1-(3-(1H-pyrrol-3-yl)phenyl)ethyl)-5-(azetidin-3-ylamino)-2-methylbenzamide (XR8-9)**. A flask fitted with a rubber septum was charged with tert-butyl (R)-3-((3-((1-(3-bromophenyl)ethyl)carbamoyl)-4-methylphenyl)amino)azetidine-1-carboxylate (**S3**) (49 mg, 0.10 mmol), (1H-pyrrol-3-yl)boronic acid (17 mg, 0.15 mmol), XPhos Pd G2 (8 mg, 0.01 mmol), $K_3PO_4$ (64 mg, 0.3 mmol), DMF/EtOH/$H_2O$ (1 mL/ 1 mL/ 0.5 mL) and then purged with argon. The mixture was stirred at 95 $^{o}$C overnight. The reaction mixture was then cooled to room temperature, diluted with ethyl acetate (20 mL), filtered through celite and concentrated in vacuo. After purification by Prep-HPLC, the product was subjected to general N-Boc deprotection procedure with HCl (4M in dioxane, 100 µL) and DCM (2 mL). The purification by Prep-HPLC afforded the product **XR8-9** (15 mg, yield 41% for 2 steps) as a white solid: $[\alpha]^{25}_{546}$ = +12.1 (c 1.7, MeOH); $^{1}$H NMR (400 MHz, Methanol-*d*$_4$) δ 8.44 (s, 1H), 7.59 – 7.56 (m, 1H), 7.44 – 7.40 (m, 1H), 7.29 – 7.24 (m, 1H), 7.16 – 7.10 (m, 2H), 7.02 (d, *J* = 8.3 Hz, 1H), 6.79 – 6.76 (m, 1H), 6.56 (dd, *J* = 8.3, 2.6 Hz, 1H), 6.51 (d, *J* = 2.6 Hz, 1H), 6.48 – 6.44 (m, 1H), 5.19 (q, *J* = 7.0 Hz, 1H), 4.49 – 4.39 (m, 1H), 4.33 – 4.27 (m, 2H), 3.90 (ddd, *J* = 11.4, 6.7, 2.0 Hz, 2H), 2.22 (s, 3H), 1.54 (d, *J* = 7.1 Hz, 3H); $^{13}$C NMR (100 MHz, Methanol-*d*$_4$) δ 172.50, 145.42, 139.03, 138.29, 132.64, 132.61, 129.72, 125.71, 125.36, 124.64, 123.70, 119.83, 115.72, 115.69, 115.65, 112.63, 106.47, 54.83, 50.67, 46.83, 22.49, 18.58; HRMS (ESI) calcd for $C_{23}H_{27}N_4O$ $[M+H]^+$ 375.2179, found 375.2183.

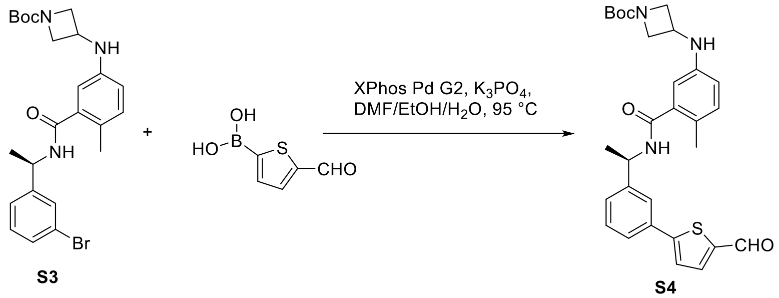


**S3** **S4**

**Tert-butyl (R)-3-((3-((1-(3-(5-formylthiophen-2-yl)phenyl)ethyl)carbamoyl)-4-methylphenyl)amino)azetidine-1-carboxylate (S4)**. A flask fitted with a rubber septum was charged with tert-butyl (R)-3-((3-((1-(3-bromophenyl)ethyl)carbamoyl)-4-methylphenyl)amino)azetidine-1-carboxylate (**S3**) (490 mg, 1.00 mmol), (5-formylthiophen-2-yl)boronic acid (170 mg, 1.50 mmol), XPhos Pd G2 (40 mg, 0.05 mmol), $K_3PO_4$ (531 mg, 2.5 mmol), DMF/EtOH/$H_2O$ (5 mL/ 5 mL/ 2.5 mL) and then purged with argon. The mixture was stirred at 95 $^{o}$C overnight. The reaction mixture was then cooled to room temperature, diluted with ethyl acetate (50 mL), filtered through celite and concentrated in vacuo. The purification by Prep-

HPLC afforded the **XR8-6** (166 mg, yield 32%) as a white solid: $^{1}$H NMR (400 MHz, Chloroform-*d*) δ 9.88 (s, 1H), 7.74 (d, *J* = 3.9 Hz, 1H), 7.67 (s, 1H), 7.61 – 7.57 (m, 1H), 7.46 – 7.38 (m, 3H), 7.02 (d, *J* = 8.2 Hz, 1H), 6.55 (d, *J* = 2.5 Hz, 1H), 6.47 (dd, *J* = 8.2, 2.6 Hz, 1H), 6.06 (d, *J* = 7.9 Hz, 1H), 5.34 (p, *J* = 7.1 Hz, 1H), 4.25 (ddd, *J* = 9.2, 6.9, 2.6 Hz, 2H), 4.12 (q, *J* = 7.2 Hz, 1H), 3.68 (dd, *J* = 8.9, 4.5 Hz, 2H), 2.29 (s, 3H), 1.61 (d, *J* = 7.0 Hz, 3H), 1.43 (s, 9H); $^{13}$C NMR (100 MHz, Chloroform-*d*) δ 182.92, 169.49, 154.08, 144.65, 144.28, 142.72, 137.52, 137.36, 133.64, 132.20, 129.79, 127.32, 125.64, 125.22, 124.47, 124.37, 114.75, 111.84, 79.88, 49.07, 43.42, 28.51, 22.12, 18.89; LRMS (ESI) calcd for $C_{29}H_{34}N_3O_4S$ $[M+H]^+$ 520.2, found 520.3.

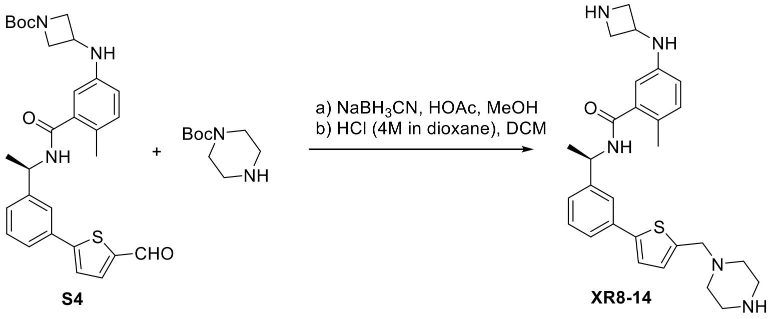


**(R)-5-(azetidin-3-ylamino)-2-methyl-N-(1-(3-(5-(piperazin-1-ylmethyl)thiophen-2-yl)phenyl)ethyl)benzamide (XR8-14)**. Tert-butyl (R)-3-((3-((1-(3-(5-formylthiophen-2-yl)phenyl)ethyl)carbamoyl)-4-methylphenyl)amino)azetidine-1-carboxylate (**S4**) (31 mg, 0.06 mmol) and tert-butyl piperazine-1-carboxylate (17 mg, 0.09 mmol) was subjected to general reductive amination procedure with MeOH (2 mL), HOAc (500 µL) at 50 °C, and then $NaBH_3CN$ (12 mg, 0.18 mmol) was added. After purification by Prep-HPLC, the product was subjected to general N-Boc deprotection procedure with HCl (4M in dioxane, 100 µL) and DCM (2 mL). The purification by Prep-HPLC afforded the product **XR8-14** (20 mg, yield 68% for 2 steps) as a white solid: $[\alpha]_{546}^{25}$ = -6.9 (c 1.7, MeOH); $^{1}$H NMR (400 MHz, Methanol-*d*$_4$) δ 8.34 (s, 1H), 7.67 – 7.64 (m, 1H), 7.53 – 7.48 (m, 1H), 7.41 – 7.32 (m, 2H), 7.26 (d, *J* = 3.7 Hz, 1H), 7.03 (d, *J* = 7.9 Hz, 1H), 6.99 – 6.97 (m, 1H), 6.61 – 6.55 (m, 2H), 5.21 (q, *J* = 7.1 Hz, 1H), 4.56 – 4.45 (m, 1H), 4.36 (dd, *J* = 11.4, 7.4 Hz, 2H), 3.95 (dd, *J* = 11.2, 6.7 Hz, 2H), 3.83 (s, 2H), 3.26 – 3.21 (m, 4H), 2.80 – 2.73 (m, 4H), 2.21 (s, 3H), 1.55 (d, *J* = 7.0 Hz, 3H); $^{13}$C NMR (100 MHz, Methanol-*d*$_4$) δ 172.57, 146.33, 145.64, 145.47, 141.01, 138.86, 135.95, 132.64, 130.28, 129.15, 126.33, 125.73, 125.27, 124.53, 123.90, 115.59, 112.84, 57.59, 54.92, 50.54, 50.31, 46.84, 44.87, 22.36, 18.64; HRMS (ESI) calcd for $C_{28}H_{36}N_5OS$ $[M+H]^+$ 490.2635, found 490.2632.

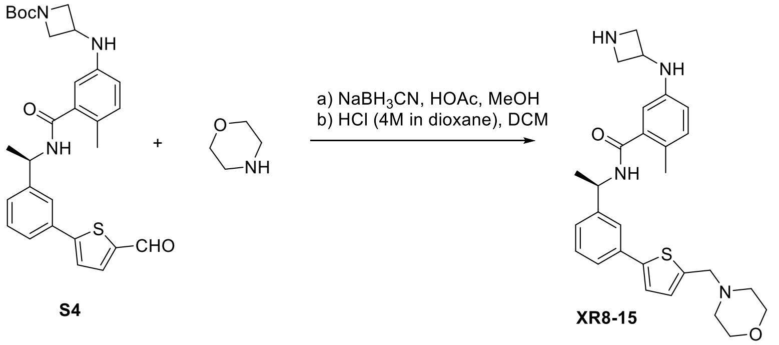


**(R)-5-(azetidin-3-ylamino)-2-methyl-N-(1-(3-(5-(morpholinomethyl)thiophen-2-yl)phenyl)ethyl)benzamide (XR8-15)**. Tert-butyl (R)-3-((3-((1-(3-(5-formylthiophen-2-yl)phenyl)ethyl)carbamoyl)-4-methylphenyl)amino)azetidine-1-carboxylate (**S4**) (31 mg, 0.06

mmol) and morpholine (8 mg, 0.09 mmol) was subjected to general reductive amination procedure with MeOH (2 mL), HOAc (500 µL) at 50 °C, and then $NaBH_3CN$ (12 mg, 0.18 mmol) was added. After purification by Prep-HPLC, the product was subjected to general N-Boc deprotection procedure with HCl (4M in dioxane, 100 µL) and DCM (2 mL). The purification by Prep-HPLC afforded the product **XR8-15** (19 mg, yield 65 % for 2 steps) as a white solid: $[\alpha]_{546}^{25}$ = -9.6 (c 1.2, MeOH); $^1$H NMR (400 MHz, Methanol-*d*$_4$) δ 8.09 (s, 1H), 7.73 – 7.69 (m, 1H), 7.60 – 7.54 (m, 1H), 7.46 – 7.37 (m, 4H), 7.08 (d, *J* = 8.6 Hz, 1H), 6.72 – 6.66 (m, 2H), 5.22 (q, *J* = 7.0 Hz, 1H), 4.64 (s, 2H), 4.59 – 4.52 (m, 1H), 4.42 – 4.32 (m, 2H), 4.13 – 3.99 (m, 4H), 3.89 – 3.79 (m, 2H), 3.52 – 3.43 (m, 2H), 3.29 – 3.19 (m, 2H), 2.22 (s, 3H), 1.56 (d, *J* = 7.1 Hz, 3H); $^{13}$C NMR (100 MHz, Methanol-*d*$_4$) δ 172.36, 149.60, 146.57, 144.17, 138.73, 135.32, 134.98, 132.81, 130.51, 129.40, 127.29, 127.09, 125.68, 124.98, 124.86, 116.51, 113.87, 64.99, 55.70, 54.64, 52.49, 50.63, 47.39, 22.40, 18.71; HRMS (ESI) calcd for $C_{28}H_{35}N_4O_2S$ $[M+H]^+$ 491.2475, found 491.2475.

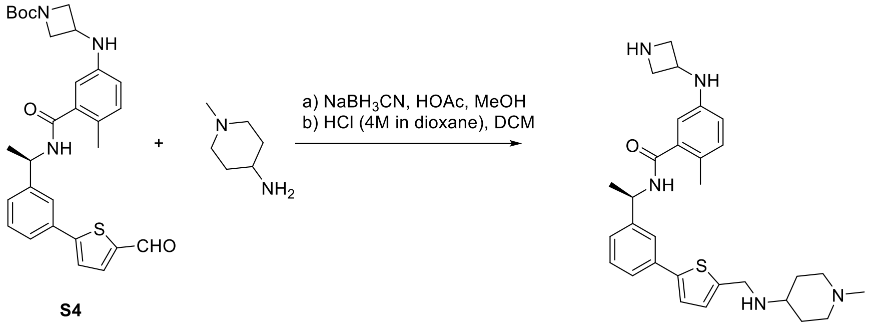


S4

XR8-16

**(R)-5-(azetidin-3-ylamino)-2-methyl-N-(1-(3-(5-(((1-methylpiperidin-4-yl)amino)methyl)thiophen-2-yl)phenyl)ethyl)benzamide (XR8-16)**. Tert-butyl (R)-3-((3-((1-(3-(5-formylthiophen-2-yl)phenyl)ethyl)carbamoyl)-4-methylphenyl)amino)azetidine-1-carboxylate (**S4**) (31 mg, 0.06 mmol) and 1-methylpiperidin-4-amine (10 mg, 0.09 mmol) was subjected to general reductive amination procedure with MeOH (2 mL), HOAc (500 µL) at 50 °C, and then $NaBH_3CN$ (12 mg, 0.18 mmol) was added. After purification by Prep-HPLC, the product was subjected to general N-Boc deprotection procedure with HCl (4M in dioxane, 100 µL) and DCM (2 mL). The purification by Prep-HPLC afforded the product **XR8-16** (23 mg, yield 74 % for 2 steps) as a white solid: $[\alpha]_{546}^{25}$ = -9.6 (c 1.7, MeOH); $^1$H NMR (400 MHz, Methanol-*d*$_4$) δ 8.09 (s, 1H), 7.70 (t, *J* = 1.9 Hz, 1H), 7.56 (dt, *J* = 6.6, 2.1 Hz, 1H), 7.46 – 7.34 (m, 4H), 7.08 – 7.01 (m, 1H), 6.65 – 6.58 (m, 2H), 5.21 (q, *J* = 7.2 Hz, 1H), 4.62 – 4.47 (m, 3H), 4.42 – 4.33 (m, 2H), 4.05 – 3.96 (m, 2H), 3.70 – 3.57 (m, 3H), 3.25 – 3.13 (m, 2H), 2.89 (s, 3H), 2.49 (d, *J* = 14.2 Hz, 2H), 2.21 (s, 3H), 2.20 – 2.06 (m, 2H), 1.56 (d, *J* = 7.1 Hz, 3H); $^{13}$C NMR (100 MHz, Methanol-*d*$_4$) δ 172.52, 148.59, 146.64, 145.18, 138.77, 135.12, 133.42, 132.69, 132.18, 130.47, 127.10, 126.05, 125.58, 124.80, 124.78, 115.75, 113.11, 54.94, 53.48, 52.86, 50.58, 46.94, 44.07, 43.70, 27.36, 22.42, 18.66; HRMS (ESI) calcd for $C_{30}H_{40}N_5OS$ $[M+H]^+$ 518.2948, found 518.2952.

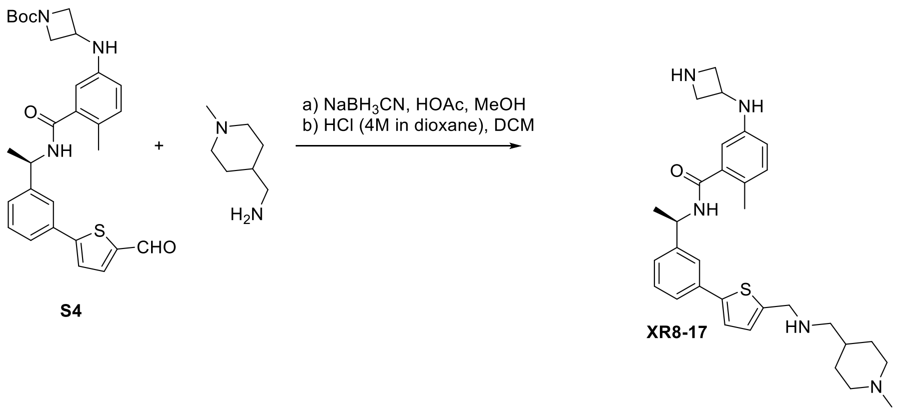


**(R)-5-(azetidin-3-ylamino)-2-methyl-N-(1-(3-(5-((((1-methylpiperidin-4-yl)methyl)amino)methyl)thiophen-2-yl)phenyl)ethyl)benzamide (XR8-17)**. Tert-butyl (R)-3-((3-((1-(3-(5-formylthiophen-2-yl)phenyl)ethyl)carbamoyl)-4-methylphenyl)amino)azetidine-1-carboxylate (**S4**) (31 mg, 0.06 mmol) and (1-methylpiperidin-4-yl)methanamine (12 mg, 0.09 mmol) was subjected to general reductive amination procedure with MeOH (2 mL), HOAc (500 µL) at 50 $^{o}$C, and then $NaBH_3CN$ (12 mg, 0.18 mmol) was added. After purification by Prep-HPLC, the product was subjected to general N-Boc deprotection procedure with HCl (4M in dioxane, 100 µL) and DCM (2 mL). The purification by Prep-HPLC afforded the **XR8-17** (24 mg, yield 76 % for 2 steps) as a white solid: $[\alpha]^{25}_{546}$ = -11.0 (c 0.6, MeOH); $^{1}$H NMR (400 MHz, Methanol-$d_4$) δ 8.16 (s, 1H), 7.72 – 7.67 (m, 1H), 7.55 (tt, *J* = 5.7, 2.1 Hz, 1H), 7.44 – 7.32 (m, 4H), 7.04 (d, *J* = 9.0 Hz, 1H), 6.61 – 6.56 (m, 2H), 5.24 – 5.16 (m, 1H), 4.55 – 4.47 (m, 3H), 4.40 – 4.34 (m, 2H), 4.00 – 3.93 (m, 2H), 3.47 – 3.37 (m, 2H), 3.09 – 2.98 (m, 4H), 2.21 (s, 3H), 2.18 – 2.02 (m, 3H), 1.56 (d, *J* = 7.1 Hz, 5H); $^{13}$C NMR (100 MHz, Methanol-$d_4$) δ 172.60, 148.54, 146.64, 145.47, 138.83, 135.15, 133.43, 132.66, 132.50, 130.47, 127.02, 125.76, 125.60, 124.90, 124.73, 115.55, 112.84, 55.06, 52.47, 50.56, 47.01, 46.85, 44.47, 32.54, 27.57, 22.37, 18.63; HRMS (ESI) calcd for $C_{31}H_{42}N_5OS$ $[M+H]^+$ 532.3105, found 518.2952.

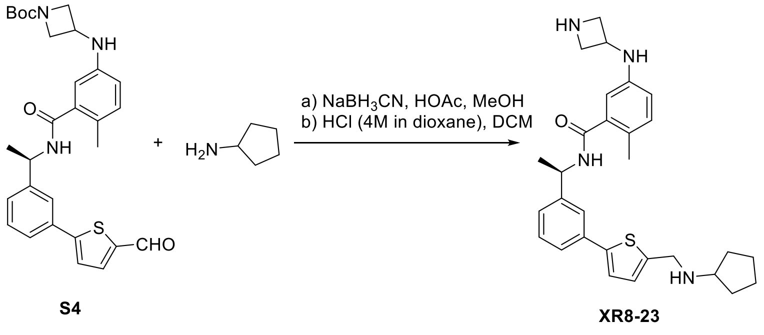


**(R)-5-(azetidin-3-ylamino)-N-(1-(3-(5-((cyclopentylamino)methyl)thiophen-2-yl)phenyl)ethyl)-2-methylbenzamide (XR8-23)**. Tert-butyl (R)-3-((3-((1-(3-(5-formylthiophen-2-yl)phenyl)ethyl)carbamoyl)-4-methylphenyl)amino)azetidine-1-carboxylate (**S4**) (31 mg, 0.06 mmol) and cyclopentanamine (8 mg, 0.09 mmol) was subjected to general reductive amination procedure with MeOH (2 mL), HOAc (500 µL) at 50 $^{o}$C, and then $NaBH_3CN$ (12 mg, 0.18 mmol) was added. After purification by Prep-HPLC, the product was subjected to general N-Boc deprotection procedure with HCl (4M in dioxane, 100 µL) and DCM (2 mL). The purification by Prep-HPLC afforded the product **XR8-23** (21 mg, yield 71 % for 2 steps) as a white solid: $[\alpha]^{25}_{546}$ = -3.1 (c 0.2, MeOH); $^{1}$H NMR (400 MHz, Methanol-$d_4$) δ 8.45 (s, 1H), 7.72 – 7.67 (m, 1H), 7.57 –

7.52 (m, 1H), 7.43 – 7.37 (m, 3H), 7.28 (d, *J* = 3.8 Hz, 1H), 7.03 (d, *J* = 8.1 Hz, 1H), 6.60 – 6.54 (m, 2H), 5.21 (q, *J* = 7.1 Hz, 1H), 4.54 – 4.46 (m, 1H), 4.44 (s, 2H), 4.39 – 4.32 (m, 2H), 3.98 – 3.89 (m, 2H), 3.67 – 3.57 (m, 1H), 2.23 – 2.12 (m, 5H), 1.89 – 1.78 (m, 2H), 1.75 – 1.65 (m, 4H), 1.55 (d, *J* = 7.1 Hz, 3H); $^{13}$C NMR (100 MHz, Methanol-$d_4$) δ 172.60, 148.25, 146.62, 145.50, 138.86, 135.16, 133.08, 132.80, 132.64, 130.46, 126.96, 125.71, 125.61, 124.94, 124.75, 115.56, 112.80, 59.87, 54.88, 50.55, 46.86, 45.41, 30.73, 25.04, 22.34, 18.62; HRMS (ESI) calcd for $C_{29}H_{37}N_4OS$ [M+H]$^+$ 489.2683, found 489.2663.

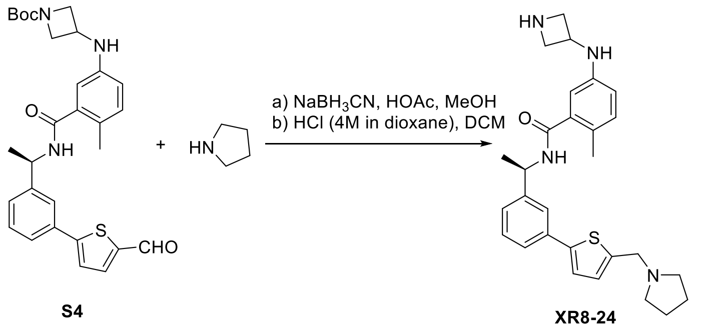


S4 XR8-24

**(R)-5-(azetidin-3-ylamino)-2-methyl-N-(1-(3-(5-(pyrrolidin-1-ylmethyl)thiophen-2-yl)phenyl)ethyl)benzamide (XR8-24)**. Tert-butyl (R)-3-((3-((1-(3-(5-formylthiophen-2-yl)phenyl)ethyl)carbamoyl)-4-methylphenyl)amino)azetidine-1-carboxylate (**S4**) (31 mg, 0.06 mmol) and pyrrolidine (7 mg, 0.09 mmol) was subjected to general reductive amination procedure with MeOH (2 mL), HOAc (500 µL) at 50 °C, and then $NaBH_3CN$ (12 mg, 0.18 mmol) was added. After purification by Prep-HPLC, the product was subjected to general N-Boc deprotection procedure with HCl (4M in dioxane, 100 µL) and DCM (2 mL). The purification by Prep-HPLC afforded the product **XR8-24** (15 mg, yield 54 % for 2 steps) as a white solid: $^1$H NMR (400 MHz, Methanol-$d_4$) δ 8.44 (s, 1H), 7.72 – 7.67 (m, 1H), 7.58 – 7.53 (m, 1H), 7.44 – 7.37 (m, 3H), 7.31 – 7.27 (m, 1H), 7.03 (d, *J* = 7.9 Hz, 1H), 6.62 – 6.55 (m, 2H), 5.22 (q, *J* = 7.0 Hz, 1H), 4.57 (s, 2H), 4.53 – 4.46 (m, 1H), 4.39 – 4.32 (m, 2H), 3.94 (dd, *J* = 11.2, 6.5 Hz, 2H), 3.40 – 3.34 (m, 4H), 2.21 (s, 3H), 2.12 – 2.05 (m, 4H), 1.55 (d, *J* = 7.1 Hz, 3H); $^{13}$C NMR (100 MHz, Methanol-$d_4$) δ 172.59, 148.64, 146.63, 145.49, 138.86, 135.11, 133.50, 132.72, 132.65, 130.47, 127.09, 125.73, 125.63, 124.91, 124.76, 115.56, 112.83, 54.90, 54.47, 53.16, 50.53, 46.86, 23.99, 22.36, 18.62; HRMS (ESI) calcd for $C_{28}H_{35}N_4OS$ [M+H]$^+$ 475.2526, found 475.2524.

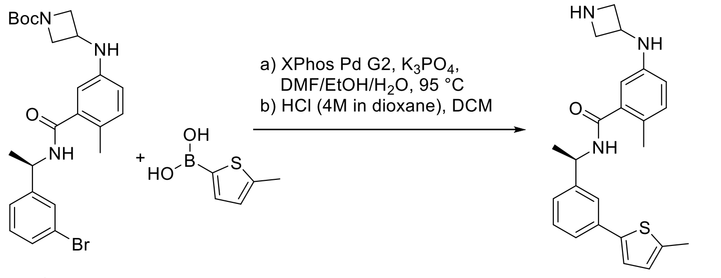


S3 XR8-30

**(R)-5-(azetidin-3-ylamino)-2-methyl-N-(1-(3-(5-methylthiophen-2-yl)phenyl)ethyl)benzamide (XR8-30)**. A flask fitted with a rubber septum was charged with tert-butyl (R)-3-((3-((1-(3-bromophenyl)ethyl)carbamoyl)-4-methylphenyl)amino)azetidine-1-carboxylate (**S3**) (49 mg, 0.10 mmol), (5-methylthiophen-2-yl)boronic acid (21 mg, 0.15 mmol), XPhos Pd G2 (8 mg, 0.01 mmol), $K_3PO_4$ (64 mg, 0.3 mmol), DMF/EtOH/$H_2O$ (1 mL/ 1 mL/ 0.5

mL) and then purged with argon. The mixture was stirred at 95 °C overnight. The reaction mixture was then cooled to room temperature, diluted with ethyl acetate (20 mL), filtered through celite and concentrated in vacuo. After purification by Prep-HPLC, the product was subjected to general N-Boc deprotection procedure with HCl (4M in dioxane, 100 µL) and DCM (2 mL). The purification by Prep-HPLC afforded the **XR8-30** (19 mg, yield 47% for 2 steps) as a white solid: $[\alpha]^{25}_{546}$ = +14.7 (c 1.5, MeOH); $^1$H NMR (400 MHz, Methanol-$d_4$) δ 8.49 (s, 1H), 7.64 – 7.59 (m, 1H), 7.48 – 7.43 (m, 1H), 7.36 – 7.26 (m, 2H), 7.17 (d, *J* = 3.7 Hz, 1H), 7.03 (d, *J* = 8.2 Hz, 1H), 6.75 (dd, *J* = 3.5, 1.2 Hz, 1H), 6.59 – 6.52 (m, 2H), 5.20 (q, *J* = 7.0 Hz, 1H), 4.52 – 4.42 (m, 1H), 4.36 – 4.29 (m, 2H), 3.95 – 3.88 (m, 2H), 2.48 (d, *J* = 1.0 Hz, 3H), 2.22 (s, 3H), 1.54 (d, *J* = 7.1 Hz, 3H); $^{13}$C NMR (100 MHz, Methanol-$d_4$) δ 172.56, 146.15, 145.45, 143.05, 140.67, 138.93, 136.30, 132.63, 130.14, 127.43, 125.92, 125.71, 125.06, 124.17, 124.14, 115.62, 112.71, 54.84, 50.52, 46.84, 22.39, 18.63, 15.29; HRMS (ESI) calcd for $C_{24}H_{28}N_3OS$ $[M+H]^+$ 406.1948, found 406.1958.

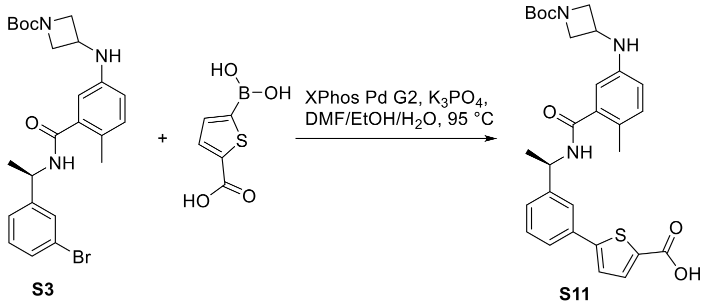


**(R)-5-(3-(1-(5-((1-(tert-butoxycarbonyl)azetidin-3-yl)amino)-2-methylbenzamido)ethyl)phenyl)thiophene-2-carboxylic acid (S11)**. A flask fitted with a rubber septum was charged with tert-butyl (R)-3-((3-((1-(3-bromophenyl)ethyl)carbamoyl)-4-methylphenyl)amino)azetidine-1-carboxylate (490 mg, 1.00 mmol), 5-boronothiophene-2-carboxylic acid (258 mg, 1.50 mmol), XPhos Pd G2 (40 mg, 0.05 mmol), $K_3PO_4$ (531 mg, 2.5 mmol), DMF/EtOH/$H_2O$ (5 mL/ 5 mL/ 2.5 mL) and then purged with argon. The mixture was stirred at 95 °C overnight. The reaction mixture was then cooled to room temperature, diluted with ethyl acetate (50 mL), filtered through celite and concentrated in vacuo. The purification by Prep-HPLC afforded the product **S11** (150 mg, yield 28%) as a white solid: $^1$H NMR (400 MHz, Methanol-$d_4$) δ 7.77 – 7.71 (m, 2H), 7.63 – 7.58 (m, 1H), 7.44 – 7.39 (m, 3H), 7.00 (d, *J* = 8.0 Hz, 1H), 6.58 – 6.51 (m, 2H), 5.22 (q, *J* = 7.0 Hz, 1H), 4.28 – 4.16 (m, 3H), 3.73 – 3.65 (m, 2H), 2.21 (s, 3H), 1.55 (d, *J* = 7.1 Hz, 3H), 1.43 (s, 9H); $^{13}$C NMR (100 MHz, Methanol-$d_4$) δ 172.83, 158.16, 146.73, 146.29, 138.69, 135.32, 135.12, 132.48, 130.44, 127.77, 125.78, 125.06, 124.79, 115.71, 112.60, 81.05, 50.37, 44.18, 28.64, 22.42, 18.58; LRMS (ESI) calcd 536.2, found 536.3.

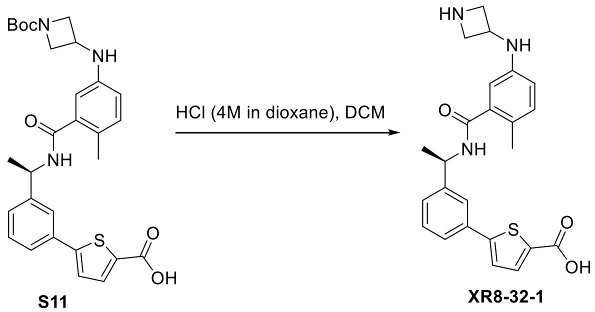

**(R)-5-(3-(1-(5-(azetidin-3-ylamino)-2-methylbenzamido)ethyl)phenyl)thiophene-2-carboxylic acid (XR8-32-1)**. (R)-5-(3-(1-(5-((1-(tert-butoxycarbonyl)azetidin-3-yl)amino)-2-methylbenzamido)ethyl)phenyl)thiophene-2-carboxylic acid (**S11**) (20 mg, 0.04 mmol) was subjected to general N-Boc deprotection procedure with HCl (4M in dioxane, 100 µL) and DCM (2 mL). The purification by Prep-HPLC afforded the **XR8-32-1** (14 mg, yield 86%) as a white solid: $[\alpha]^{25}_{546}$ = +61.7 (c 0.3, MeOH); $^{1}$H NMR (400 MHz, Methanol-$d_4$) δ 7.75 – 7.72 (m, 1H), 7.65 – 7.62 (m, 1H), 7.58 (d, $J$ = 3.8 Hz, 1H), 7.44 – 7.32 (m, 3H), 7.05 (d, $J$ = 8.3 Hz, 1H), 6.68 – 6.61 (m, 2H), 5.21 (q, $J$ = 7.0 Hz, 1H), 4.67 (p, $J$ = 7.3 Hz, 1H), 4.50 – 4.37 (m, 2H), 4.05 – 3.93 (m, 2H), 2.27 (s, 3H), 1.54 (d, $J$ = 7.1 Hz, 3H); $^{13}$C NMR (100 MHz, Methanol-$d_4$) δ 172.77, 149.48, 146.65, 145.45, 138.95, 135.71, 132.75, 132.68, 130.30, 127.54, 125.77, 124.96, 124.73, 123.55, 116.74, 111.35, 55.54, 55.33, 50.12, 46.78, 22.65, 18.47; HRMS (ESI) calcd for $C_{24}H_{26}N_3O_3S$ $[M+H]^+$ 436.1689, found 436.1697.

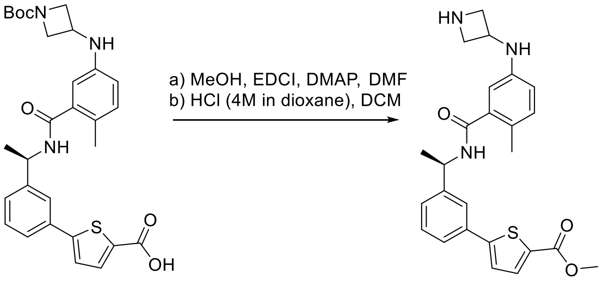


**S11** **XR8-32-2**

**Methyl (R)-5-(3-(1-(5-(azetidin-3-ylamino)-2-methylbenzamido)ethyl)phenyl)thiophene-2-carboxylate (XR8-32-2)**. **S11** (16 mg, 0.03 mmol), MeOH (5 mg, 0.15 mmol), EDCI (10 mg, 0.05 mmol) and DMAP (11 mg, 0.09 mmol) was subjected to general amine coupling procedure with DMF (1 mL). After purification by Prep-HPLC, the product was subjected to general N-Boc deprotection procedure with HCl (4M in dioxane, 100 µL) and DCM (2 mL). The purification by Prep-HPLC afforded the **XR8-32-2** (11 mg, yield 82% for 2 steps) as a white solid: $[\alpha]^{25}_{546}$ = +14.5 (c 0.4, MeOH); $^{1}$H NMR (400 MHz, Acetone-$d_6$) δ 8.21 (s, 1H), 7.89 – 7.86 (m, 1H), 7.79 (d, $J$ = 3.9 Hz, 1H), 7.67 – 7.61 (m, 1H), 7.56 – 7.42 (m, 3H), 6.95 (d, $J$ = 8.4 Hz, 1H), 6.62 (d, $J$ = 2.6 Hz, 1H), 6.55 (dd, $J$ = 8.2, 2.6 Hz, 1H), 5.38 – 5.24 (m, 1H), 4.08 (p, $J$ = 6.7 Hz, 1H), 3.87 (s, 3H), 3.80 – 3.73 (m, 2H), 3.26 – 3.19 (m, 2H), 2.20 (s, 3H), 1.59 (d, $J$ = 7.0 Hz, 3H); $^{13}$C NMR (100 MHz, Acetone-$d_6$) δ 169.76, 162.83, 151.97, 147.31, 146.22, 138.76, 135.33, 134.21, 132.90, 132.08, 130.22, 127.91, 125.32, 125.01, 124.76, 124.44, 114.67, 112.51, 54.53, 54.49, 52.46, 49.35, 43.41, 22.72, 18.81; HRMS (ESI) calcd for $C_{25}H_{28}N_3O_3S$ $[M+H]^+$ 450.1846, found 450.1842.

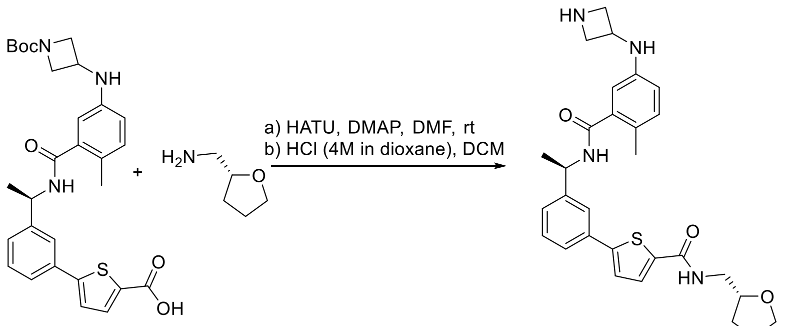


**S11** **XR8-35**

**5-(3-((R)-1-(5-(azetidin-3-ylamino)-2-methylbenzamido)ethyl)phenyl)-N-(((R)-tetrahydrofuran-2-yl)methyl)thiophene-2-carboxamide (S11). XR8-28** (16 mg, 0.03 mmol), (S)-(tetrahydrofuran-2-yl)methanamine (6 mg, 0.06 mmol), HATU (15 mg, 0.04 mmol) and DMAP (11 mg, 0.09 mmol) was subjected to general amine coupling procedure with DMF (1 mL). After purification by Prep-HPLC, the product was subjected to general N-Boc deprotection procedure with HCl (4M in dioxane, 100 µL) and DCM (2 mL). The purification by Prep-HPLC afforded the product **XR8-38** (12 mg, yield 77% for 2 steps) as a white solid: $[\alpha]^{25}_{546}$ = +2.1 (c 0.7, MeOH); $^1$H NMR (400 MHz, Methanol-$d_4$) δ 8.51 (s, 1H), 7.77 – 7.59 (m, 3H), 7.47 – 7.36 (m, 3H), 7.05 (d, *J* = 8.2 Hz, 1H), 6.68 – 6.53 (m, 2H), 5.23 (q, *J* = 7.0 Hz, 1H), 4.64 – 4.54 (m, 1H), 4.49 – 4.36 (m, 2H), 4.10 (dd, *J* = 6.9, 4.9 Hz, 1H), 4.03 – 3.85 (m, 3H), 3.77 (q, *J* = 7.2 Hz, 1H), 3.56 – 3.38 (m, 2H), 2.24 (s, 3H), 2.09 – 1.86 (m, 3H), 1.73 – 1.63 (m, 1H), 1.55 (d, *J* = 7.1 Hz, 3H); $^{13}$C NMR (100 MHz, Methanol-$d_4$) δ 171.29, 163.12, 149.06, 145.29, 144.06, 137.70, 137.54, 133.66, 131.28, 129.17, 129.04, 126.36, 124.33, 124.01, 123.63, 122.90, 114.83, 110.50, 77.77, 67.72, 53.87, 53.80, 48.90, 45.45, 43.40, 28.50, 25.19, 21.06, 17.15; HRMS (ESI) calcd for $C_{29}H_{35}N_4O_3S$ $[M+H]^+$ 519.2424, found 519.2424.

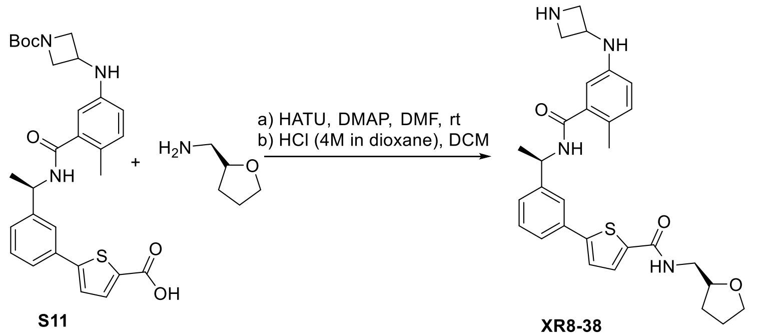


S11

XR8-38

**5-(3-((R)-1-(5-(azetidin-3-ylamino)-2-methylbenzamido)ethyl)phenyl)-N-(((R)-tetrahydrofuran-2-yl)methyl)thiophene-2-carboxamide (XR8-38)**. **S11** (16 mg, 0.03 mmol), (R)-(tetrahydrofuran-2-yl)methanamine (6 mg, 0.06 mmol), HATU (15 mg, 0.04 mmol) and DMAP (11 mg, 0.09 mmol) was subjected to general amine coupling procedure with DMF (1 mL). After purification by Prep-HPLC, the product was subjected to general N-Boc deprotection procedure with HCl (4M in dioxane, 100 µL) and DCM (2 mL). The purification by Prep-HPLC afforded the product **XR8-38** (13 mg, yield 84% for 2 steps) as a white solid: $[\alpha]^{25}_{546}$ = +56.7 (c 1.2, MeOH); $^1$H NMR (400 MHz, Methanol-$d_4$) δ 8.41 (s, 1H), 7.74 – 7.72 (m, 1H), 7.69 (d, *J* = 3.9 Hz, 1H), 7.63 – 7.59 (m, 1H), 7.45 – 7.37 (m, 3H), 7.04 (d, *J* = 8.2 Hz, 1H), 6.64 – 6.56 (m, 2H), 5.22 (q, *J* = 7.1 Hz, 1H), 4.61 – 4.52 (m, 1H), 4.46 – 4.36 (m, 2H), 4.09 (qd, *J* = 6.9, 4.6 Hz, 1H), 4.01 – 3.93 (m, 2H), 3.92 – 3.84 (m, 1H), 3.80 – 3.72 (m, 1H), 3.52 – 3.38 (m, 2H), 2.24 (s, 3H), 2.09 – 1.86 (m, 3H), 1.72 – 1.61 (m, 1H), 1.55 (d, *J* = 7.1 Hz, 3H); $^{13}$C NMR (100 MHz, Methanol-$d_4$) δ 172.68, 164.48, 150.42, 146.71, 145.47, 139.09, 138.89, 135.06, 132.68, 130.59, 130.45, 127.72, 125.72, 125.44, 125.04, 124.34, 116.18, 112.02, 79.18, 69.09, 55.11, 50.32, 46.81, 44.85, 29.93, 26.57, 22.49, 18.57; HRMS (ESI) calcd for $C_{29}H_{35}N_4O_3S$ $[M+H]^+$ 519.2424, found 519.2424.

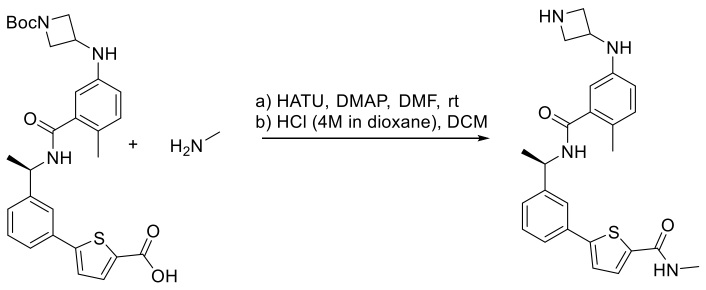

S11 XR8-39

**(R)-5-(3-(1-(5-(azetidin-3-ylamino)-2-methylbenzamido)ethyl)phenyl)-N-methylthiophene-2-carboxamide (XR8-39)**. **S11** (16 mg, 0.03 mmol), methylamine (2 M in THF, 75 µL), HATU (15 mg, 0.04 mmol) and DMAP (11 mg, 0.09 mmol) was subjected to general amine coupling procedure with DMF (1 mL). After purification by Prep-HPLC, the product was subjected to general N-Boc deprotection procedure with HCl (4M in dioxane, 100 µL) and DCM (2 mL). The purification by Prep-HPLC afforded the product **XR8-39** (12 mg, yield 89% for 2 steps) as a white solid: $[\alpha]^{25}_{546}$= +41.5 (c 0.8, MeOH); $^{1}$H NMR (400 MHz, Methanol-*d*$_4$) δ 8.45 (s, 1H), 7.75 – 7.71 (m, 1H), 7.63 – 7.59 (m, 2H), 7.45 – 7.37 (m, 3H), 7.04 (d, *J* = 8.2 Hz, 1H), 6.65 – 6.55 (m, 2H), 5.22 (q, *J* = 7.0 Hz, 1H), 4.60 – 4.52 (m, 1H), 4.46 – 4.36 (m, 2H), 4.01 – 3.92 (m, 2H), 2.92 (s, 3H), 2.23 (s, 3H), 1.55 (d, *J* = 7.1 Hz, 3H); $^{13}$C NMR (100 MHz, Methanol-*d*$_4$) δ 172.68, 164.91, 150.17, 146.69, 145.46, 139.09, 138.90, 135.10, 132.69, 130.44, 130.34, 127.68, 125.76, 125.48, 125.01, 124.40, 116.12, 112.10, 55.13, 50.35, 46.85, 26.77, 22.47, 18.56; HRMS (ESI) calcd for $C_{25}H_{29}N_4O_2S$ $[M+H]^+$ 449.2006, found 449.2015.

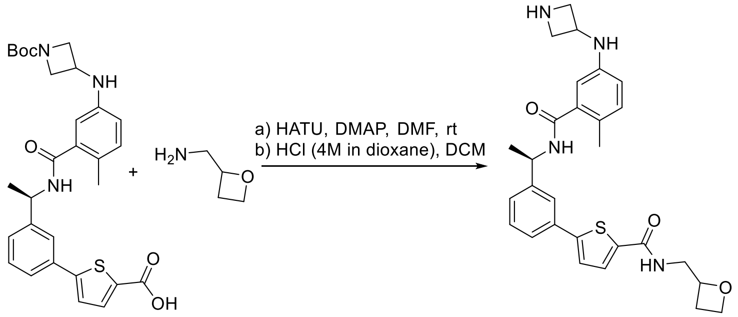

S11 XR8-40

**5-(3-((R)-1-(5-(azetidin-3-ylamino)-2-methylbenzamido)ethyl)phenyl)-N-(oxetan-2-ylmethyl)thiophene-2-carboxamide (XR8-40)**. **S11** (16 mg, 0.03 mmol), oxetan-2-ylmethanamine (5 mg, 0.06 mmol), HATU (15 mg, 0.04 mmol) and DMAP (11 mg, 0.09 mmol) was subjected to general amine coupling procedure with DMF (1 mL). After purification by Prep-HPLC, the product was subjected to general N-Boc deprotection procedure with HCl (4M in dioxane, 100 µL) and DCM (2 mL). The purification by Prep-HPLC afforded the product **XR8-40** (13 mg, yield 86% for 2 steps) as a white solid: $^{1}$H NMR (400 MHz, Methanol-*d*$_4$) δ 8.36 (s, 1H), 7.76 – 7.70 (m, 1H), 7.64 – 7.58 (m, 2H), 7.46 – 7.37 (m, 3H), 7.04 (d, *J* = 8.2 Hz, 1H), 6.64 – 6.52 (m, 2H), 5.23 (q, *J* = 7.3 Hz, 1H), 4.99 – 4.92 (m, 1H), 4.55 – 4.47 (m, 1H), 4.42 – 4.32 (m, 2H), 4.17 – 4.09 (m, 1H), 3.98 – 3.90 (m, 2H), 3.79 – 3.65 (m, 3H), 2.23 (s, 3H), 2.05 – 1.84 (m, 2H), 1.55 (d, *J* = 7.2 Hz, 3H); $^{13}$C NMR (100 MHz, Methanol-*d*$_4$) δ 172.60, 161.81, 150.50, 146.72, 145.44, 138.92, 134.94, 133.18, 132.68, 130.50, 129.71, 127.79, 125.78, 125.08, 124.73, 115.75, 112.52, 79.84, 60.38, 59.15, 55.04, 54.99, 50.47, 46.88, 39.16, 22.44, 18.60; HRMS (ESI) calcd for $C_{28}H_{33}N_4O_3S$ $[M+H]^+$ 505.2268, found 505.2260.

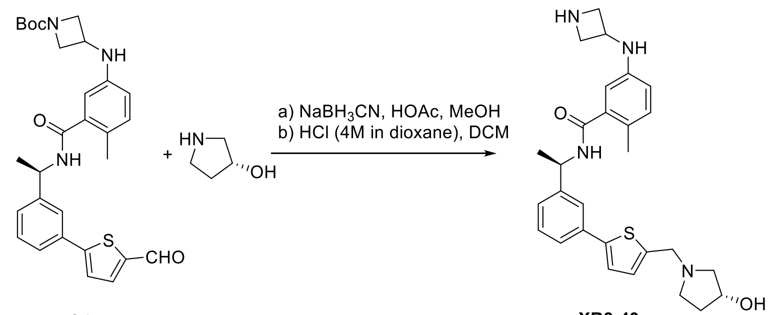


**5-(azetidin-3-ylamino)-N-((R)-1-(3-(5-(((R)-3-hydroxypyrrolidin-1-yl)methyl)thiophen-2-yl)phenyl)ethyl)-2-methylbenzamide (XR8-49)**. Tert-butyl (R)-3-((3-((1-(3-(5-formylthiophen-2-yl)phenyl)ethyl)carbamoyl)-4-methylphenyl)amino)azetidine-1-carboxylate (**S4**) (31 mg, 0.06 mmol) and (R)-pyrrolidin-3-ol (8 mg, 0.09 mmol) was subjected to general reductive amination procedure with MeOH (2 mL), HOAc (500 µL) at 50 °C, and then $NaBH_3CN$ (12 mg, 0.18 mmol) was added. After purification by Prep-HPLC, the product was subjected to general N-Boc deprotection procedure with HCl (4M in dioxane, 100 µL) and DCM (2 mL). The purification by Prep-HPLC afforded the product **XR8-49** (19 mg, yield 63 % for 2 steps) as a white solid: $[\alpha]_{546}^{25}$ = +10.0 (c 0.3, MeOH); $^1$H NMR (400 MHz, Methanol-$d_4$) δ 8.49 (s, 1H), 7.70 – 7.65 (m, 1H), 7.56 – 7.51 (m, 1H), 7.43 – 7.31 (m, 3H), 7.15 (d, *J* = 3.7 Hz, 1H), 7.04 (d, *J* = 8.3 Hz, 1H), 6.58 (dd, *J* = 8.1, 2.6 Hz, 1H), 6.54 (d, *J* = 2.7 Hz, 1H), 5.22 (q, *J* = 7.2 Hz, 1H), 4.55 – 4.44 (m, 2H), 4.38 – 4.32 (m, 2H), 4.29 – 4.20 (m, 2H), 3.96 – 3.90 (m, 2H), 3.28 – 3.19 (m, 1H), 3.17 – 3.10 (m, 1H), 3.06 – 2.94 (m, 2H), 2.27 – 2.16 (m, 4H), 1.95 – 1.86 (m, 1H), 1.55 (d, *J* = 7.1 Hz, 3H); $^{13}$C NMR (100 MHz, Methanol-$d_4$) δ 172.59, 147.10, 146.50, 145.46, 138.93, 135.57, 132.66, 131.36, 130.36, 126.77, 125.76, 125.46, 124.67, 124.36, 115.66, 112.64, 70.90, 62.56, 55.00, 54.86, 53.38, 50.53, 46.90, 34.65, 22.37, 18.61; HRMS (ESI) calcd for $C_{28}H_{35}N_4O_2S$ $[M+H]^+$ 491.2475, found 491.2483.

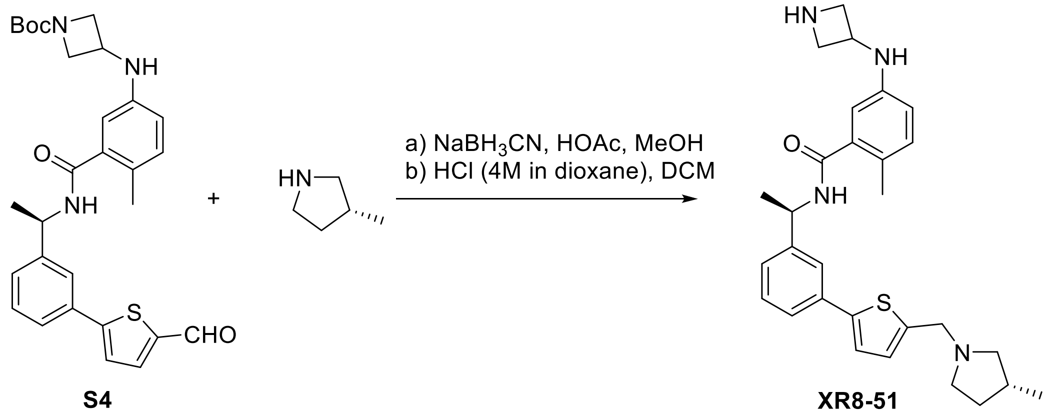


**5-(azetidin-3-ylamino)-2-methyl-N-((R)-1-(3-(5-(((R)-3-methylpyrrolidin-1-yl)methyl)thiophen-2-yl)phenyl)ethyl)benzamide (XR8-51).** Tert-butyl (R)-3-((3-((1-(3-(5-formylthiophen-2-yl)phenyl)ethyl)carbamoyl)-4-methylphenyl)amino)azetidine-1-carboxylate (**S4**) (31 mg, 0.06 mmol) and (R)-3-methylpyrrolidine (8 mg, 0.09 mmol) was subjected to general reductive amination procedure with MeOH (2 mL), HOAc (500 µL) at 50 °C, and then $NaBH_3CN$ (12 mg, 0.18 mmol) was added. After purification by Prep-HPLC, the product was subjected to general N-Boc deprotection procedure with HCl (4M in dioxane, 100 µL) and DCM (2 mL). The purification by Prep-HPLC afforded the product **XR8-56** (13 mg, yield 43 % for 2 steps) as a white solid: LRMS (ESI) calcd for $C_{29}H_{37}N_4OS$ $[M+H]^+$ 489.3, found 489.3.

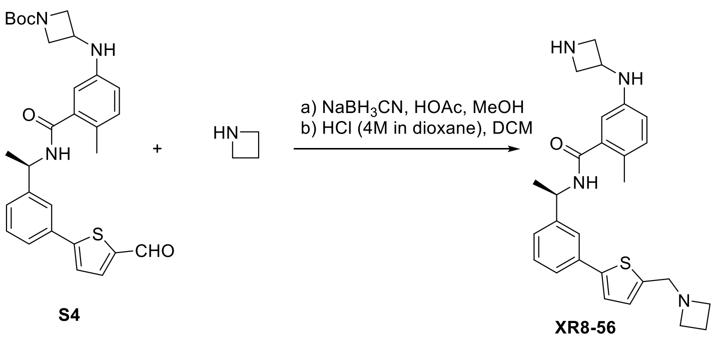


**(R)-N-(1-(3-(5-(azetidin-1-ylmethyl)thiophen-2-yl)phenyl)ethyl)-5-(azetidin-3-ylamino)-2-methylbenzamide (XR8-56)**. Tert-butyl (R)-3-((3-((1-(3-(5-formylthiophen-2-yl)phenyl)ethyl)carbamoyl)-4-methylphenyl)amino)azetidine-1-carboxylate (**S4**) (31 mg, 0.06 mmol) and azetidine hydrochloride salt (8 mg, 0.09 mmol) was subjected to general reductive amination procedure with MeOH (2 mL), HOAc (500 µL) at 50 °C, and then $NaBH_3CN$ (12 mg, 0.18 mmol) was added. After purification by Prep-HPLC, the product was subjected to general N-Boc deprotection procedure with HCl (4M in dioxane, 100 µL) and DCM (2 mL). The purification by Prep-HPLC afforded the product **XR8-56** (14 mg, yield 49 % for 2 steps) as a white solid: $^1H$ NMR (400 MHz, Methanol-$d_4$) δ 8.51 (s, 1H), 7.69 – 7.65 (m, 1H), 7.55 – 7.51 (m, 1H), 7.43 – 7.33 (m, 3H), 7.17 (d, $J$ = 3.8 Hz, 1H), 7.04 (d, $J$ = 8.2 Hz, 1H), 6.60 – 6.53 (m, 2H), 5.21 (q, $J$ = 7.0 Hz, 1H), 4.50 (p, $J$ = 6.5 Hz, 1H), 4.39 – 4.29 (m, 4H), 3.97 – 3.87 (m, 6H), 2.39 (p, $J$ = 7.8 Hz, 2H), 2.21 (s, 3H), 1.55 (d, $J$ = 7.1 Hz, 3H); $^{13}C$ NMR (100 MHz, Methanol-$d_4$) δ 172.60, 146.56, 145.47, 138.91, 135.34, 132.65, 131.71, 130.41, 126.87, 125.73, 125.53, 124.81, 124.61, 115.58, 112.73, 55.19, 55.03, 54.96, 50.55, 46.89, 22.37, 18.62, 17.39; HRMS (ESI) calcd for $C_{27}H_{33}N_4OS$ $[M+H]^+$ 461.2370, found 461.2370.

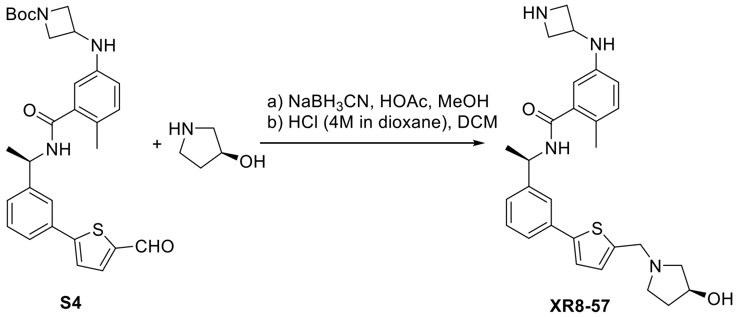


**5-(azetidin-3-ylamino)-N-((R)-1-(3-(5-(((S)-3-hydroxypyrrolidin-1-yl)methyl)thiophen-2-yl)phenyl)ethyl)-2-methylbenzamide (XR8-57)**. Tert-butyl (R)-3-((3-((1-(3-(5-formylthiophen-2-yl)phenyl)ethyl)carbamoyl)-4-methylphenyl)amino)azetidine-1-carboxylate (**S4**) (31 mg, 0.06 mmol) and (S)-pyrrolidin-3-ol (8 mg, 0.09 mmol) was subjected to general reductive amination procedure with MeOH (2 mL), HOAc (500 µL) at 50 °C, and then $NaBH_3CN$ (12 mg, 0.18 mmol) was added. After purification by Prep-HPLC, the product was subjected to general N-Boc deprotection procedure with HCl (4M in dioxane, 100 µL) and DCM (2 mL). The purification by Prep-HPLC afforded the product **XR8-57** (16 mg, yield 55 % for 2 steps) as a white solid: $^1H$

NMR (400 MHz, Methanol-$d_4$) δ 8.37 (s, 1H), 7.72 – 7.66 (m, 1H), 7.58 – 7.53 (m, 1H), 7.44 – 7.37 (m, 3H), 7.28 (d, *J* = 3.7 Hz, 1H), 7.04 (d, *J* = 8.2 Hz, 1H), 6.61 – 6.54 (m, 2H), 5.22 (q, *J* = 7.0 Hz, 1H), 4.61 – 4.46 (m, 4H), 4.40 – 4.33 (m, 2H), 3.98 – 3.91 (m, 2H), 3.53 (dt, *J* = 11.4, 8.0 Hz, 1H), 3.41 – 3.31 (m, 2H), 3.28 – 3.22 (m, 1H), 2.34 – 2.22 (m, 1H), 2.21 (s, 3H), 2.09 – 1.99 (m, 1H), 1.55 (d, *J* = 7.1 Hz, 3H); $^{13}$C NMR (100 MHz, Methanol-$d_4$) δ 172.60, 148.48, 146.61, 145.47, 138.87, 135.16, 133.36, 133.28, 132.65, 130.46, 127.02, 125.74, 125.61, 124.92, 124.70, 115.57, 112.77, 70.41, 62.07, 54.99, 54.37, 53.34, 50.55, 46.86, 34.22, 22.36, 18.62; HRMS (ESI) calcd for $C_{28}H_{35}N_4O_2S$ $[M+H]^+$ 491.2475, found 491.2483.

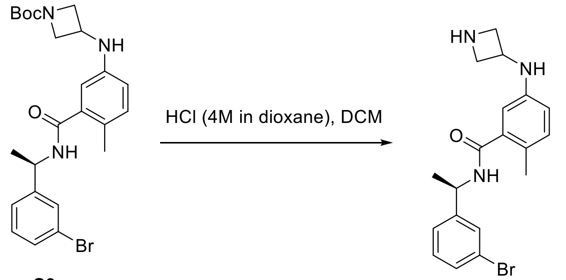


S3

XR8-61

**(R)-5-(azetidin-3-ylamino)-N-(1-(3-bromophenyl)ethyl)-2-methylbenzamide (XR8-61)**. **S3** (20 mg, 0.04 mmol) was subjected to general N-Boc deprotection procedure with HCl (4M in dioxane, 100 µL) and DCM (2 mL). The purification by Prep-HPLC afforded the product **XR8-61** (14 mg, yield 86%) as a white solid: $[\alpha]_{546}^{25}$ = +16.4 (c 0.9, MeOH); $^1$H NMR (400 MHz, Methanol-$d_4$) δ 7.62 – 7.54 (m, 1H), 7.43 – 7.35 (m, 2H), 7.30 – 7.23 (m, 1H), 7.03 (d, *J* = 8.2 Hz, 1H), 6.60 – 6.51 (m, 2H), 5.15 (q, *J* = 7.1 Hz, 1H), 4.50 (p, *J* = 6.8 Hz, 1H), 4.36 (t, *J* = 9.1 Hz, 2H), 3.97 – 3.88 (m, 2H), 2.20 (s, 3H), 1.50 (d, *J* = 7.1 Hz, 3H); 13C NMR (100 MHz, Methanol-$d_4$) δ 172.58, 148.13, 145.47, 138.75, 132.66, 131.41, 131.16, 130.33, 126.19, 125.71, 123.47, 115.63, 112.68, 54.89, 50.15, 46.84, 22.21, 18.53.HRMS (ESI) calcd for $C_{19}H_{23}BrN_3O$ $[M+H]^+$ 388.1019, found 388.1022.

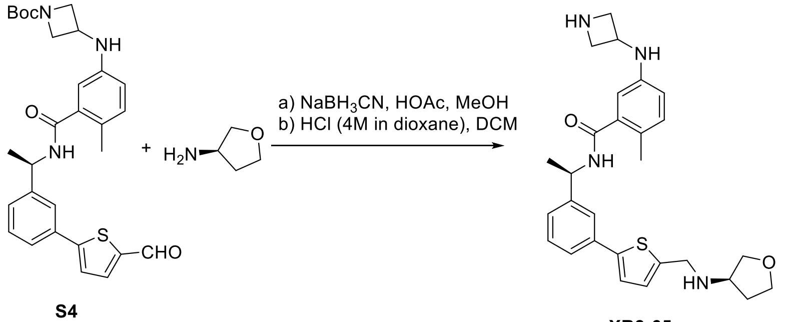


S4

XR8-65

**5-(azetidin-3-ylamino)-2-methyl-N-((R)-1-(3-(5-((((R)-tetrahydrofuran-3-yl)amino)methyl)thiophen-2-yl)phenyl)ethyl)benzamide (XR8-65)**. **S4** (31 mg, 0.06 mmol) and (R)-tetrahydrofuran-3-amine (8 mg, 0.09 mmol) was subjected to general reductive amination procedure with MeOH (2 mL), HOAc (500 µL) at 50 °C, and then $NaBH_3CN$ (12 mg, 0.18 mmol) was added. After purification by Prep-HPLC, the product was subjected to general N-Boc deprotection procedure with HCl (4M in dioxane, 100 µL) and DCM (2 mL). The purification by Prep-HPLC afforded the **XR8-65** (21 mg, yield 71 % for 2 steps) as a white solid: $[\alpha]_{546}^{25}$= +1.5 (c 0.4, MeOH); $^1$H NMR (400 MHz, Methanol-$d_4$) δ 8.36 (s, 1H), 7.71 – 7.66 (m, 1H), 7.57 – 7.52 (m, 1H), 7.44 – 7.34 (m, 3H), 7.22 (d, *J* = 3.7 Hz, 1H), 7.03 (d, *J* = 8.2 Hz, 1H), 6.61 – 6.52 (m,

2H), 5.21 (q, *J* = 7.0 Hz, 1H), 4.50 (p, *J* = 6.9 Hz, 1H), 4.40 – 4.30 (m, 4H), 4.03 (td, *J* = 8.4, 5.4 Hz, 1H), 3.97 – 3.81 (m, 5H), 3.75 (td, *J* = 8.4, 6.8 Hz, 1H), 2.41 – 2.28 (m, 1H), 2.20 (s, 3H), 2.08 – 1.97 (m, 1H), 1.55 (d, *J* = 7.1 Hz, 3H); $^{13}$C NMR (100 MHz, Methanol-$d_4$) δ 172.59, 147.63, 146.56, 145.47, 138.89, 135.36, 135.25, 132.65, 131.92, 130.41, 126.84, 125.74, 125.54, 124.83, 124.62, 115.59, 112.76, 71.32, 68.06, 59.04, 54.94, 50.55, 46.88, 45.74, 31.17, 22.34, 18.61; HRMS (ESI) calcd for $C_{28}H_{35}N_4O_2S$ $[M+H]^+$ 491.2475, found 491.2483.

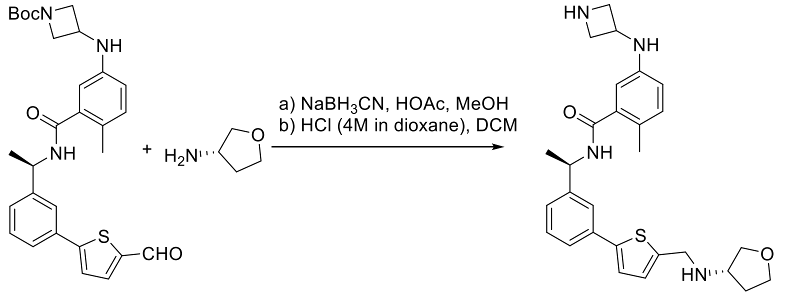


S4 XR8-66

**5-(azetidin-3-ylamino)-2-methyl-N-((R)-1-(3-(5-((((S)-tetrahydrofuran-3-yl)amino)methyl)thiophen-2-yl)phenyl)ethyl)benzamide (XR8-66)**. **S4** (31 mg, 0.06 mmol) and (S)-tetrahydrofuran-3-amine (8 mg, 0.09 mmol) was subjected to general reductive amination procedure with MeOH (2 mL), HOAc (500 µL) at 50 °C, and then $NaBH_3CN$ (12 mg, 0.18 mmol) was added. After purification by Prep-HPLC, the product was subjected to general N-Boc deprotection procedure with HCl (4M in dioxane, 100 µL) and DCM (2 mL). The purification by Prep-HPLC afforded the **XR8-66** (19 mg, yield 65 % for 2 steps) as a white solid: $[\alpha]^{25}_{546}$= +6.7 (c 0.9, MeOH); $^1$H NMR (400 MHz, Methanol-$d_4$) δ 8.43 (s, 1H), 7.67 (t, *J* = 1.8 Hz, 1H), 7.56 – 7.50 (m, 1H), 7.42 – 7.30 (m, 3H), 7.12 (d, *J* = 3.7 Hz, 1H), 7.04 (d, *J* = 8.3 Hz, 1H), 6.58 (dd, *J* = 8.2, 2.6 Hz, 1H), 6.53 (d, *J* = 2.6 Hz, 1H), 5.21 (q, *J* = 7.1 Hz, 1H), 4.54 – 4.45 (m, 1H), 4.39 – 4.31 (m, 2H), 4.22 – 4.12 (m, 2H), 4.03 – 3.89 (m, 3H), 3.86 – 3.64 (m, 4H), 2.30 – 2.23 (m, 1H), 2.21 (s, 3H), 1.98 – 1.87 (m, 1H), 1.55 (d, *J* = 7.1 Hz, 3H); $^{13}$C NMR (100 MHz, Methanol-$d_4$) δ 171.18, 145.03, 144.04, 137.56, 134.30, 131.25, 128.92, 128.75, 125.20, 124.34, 124.02, 123.24, 122.95, 114.23, 111.24, 71.04, 66.72, 57.49, 53.59, 49.13, 45.50, 45.19, 30.79, 20.95, 17.20; HRMS (ESI) calcd for $C_{28}H_{35}N_4O_2S$ $[M+H]^+$ 491.2475, found 491.2479.

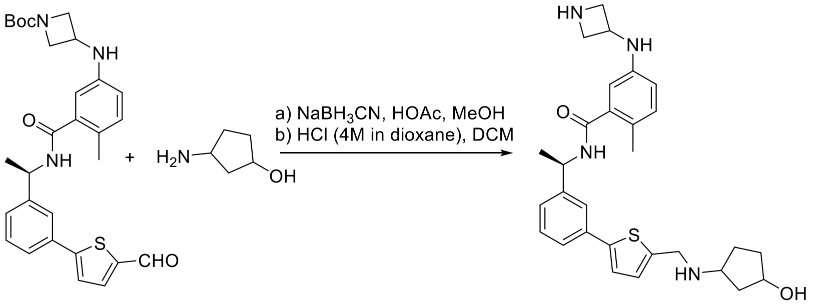


S4 XR8-67

**5-(azetidin-3-ylamino)-N-((1R)-1-(3-(5-(((3-hydroxycyclopentyl)amino)methyl)thiophen-2-yl)phenyl)ethyl)-2-methylbenzamide (XR8-67)**. **S4** (31 mg, 0.06 mmol) and 3-aminocyclopentan-1-ol (9 mg, 0.09 mmol) was subjected to general reductive amination procedure with MeOH (2 mL), HOAc (500 µL) at 50 °C, and then $NaBH_3CN$ (12 mg, 0.18 mmol) was added. After purification by Prep-HPLC, the product was subjected to general N-Boc deprotection procedure with HCl (4M in dioxane, 100 µL) and DCM (2 mL). The purification by Prep-HPLC afforded the product **XR8-67** (22 mg, yield 73 % for 2 steps) as a white solid: $^1$H NMR (400 MHz, Methanol-$d_4$) δ 8.44 (s, 1H), 7.69 (s, 1H), 7.55 (d, $J$ = 6.6 Hz, 1H), 7.44 – 7.37 (m, 3H), 7.27 (d, $J$ = 3.7 Hz, 1H), 7.03 (d, $J$ = 8.0 Hz, 1H), 6.62 – 6.54 (m, 2H), 5.26 – 5.15 (m, 1H), 4.54 – 4.26 (m, 6H), 3.98 – 3.90 (m, 2H), 3.73 – 3.63 (m, 1H), 2.30 – 2.12 (m, 5H), 2.04 – 1.92 (m, 1H), 1.88 – 1.79 (m, 3H), 1.55 (d, $J$ = 7.0 Hz, 3H); $^{13}$C NMR (100 MHz, Methanol-$d_4$) δ 172.60, 148.28, 146.62, 145.49, 138.86, 135.16, 133.09, 132.84, 132.64, 130.46, 126.94, 125.71, 125.60, 124.95, 124.76, 115.56, 112.79, 72.55, 58.48, 54.92, 50.55, 46.87, 45.19, 39.33, 34.42, 28.38, 22.34, 18.61; HRMS (ESI) calcd for $C_{29}H_{37}N_4O_2S$ [M+H]$^+$ 505.2632, found 505.2637.

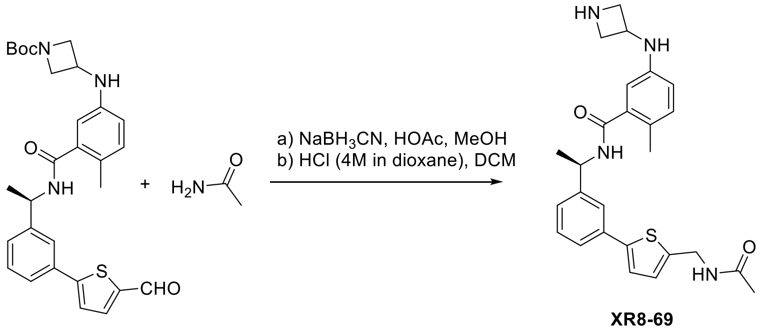


S4

XR8-69

**(R)-N-(1-(3-(5-(acetamidomethyl)thiophen-2-yl)phenyl)ethyl)-5-(azetidin-3-ylamino)-2-methylbenzamide (XR8-69)**. **S4** (31 mg, 0.06 mmol) and acetamide (6 mg, 0.09 mmol) was subjected to general reductive amination procedure with MeOH (2 mL), HOAc (500 µL) at 50 °C, and then $NaBH_3CN$ (12 mg, 0.18 mmol) was added. After purification by Prep-HPLC, the product was subjected to general N-Boc deprotection procedure with HCl (4M in dioxane, 100 µL) and DCM (2 mL). The purification by Prep-HPLC afforded the product **XR8-69** (20 mg, yield 73 % for 2 steps) as a white solid: $^1$H NMR (400 MHz, Methanol-$d_4$) δ 8.55 (s, 2H), 7.65 – 7.61 (m, 1H), 7.52 – 7.48 (m, 1H), 7.39 – 7.29 (m, 2H), 7.23 (d, $J$ = 3.6 Hz, 1H), 7.03 (d, $J$ = 8.2 Hz, 1H), 6.96 (d, $J$ = 3.7 Hz, 1H), 6.58 (dd, $J$ = 8.2, 2.6 Hz, 1H), 6.53 (d, $J$ = 2.6 Hz, 1H), 5.23 – 5.17 (m, 1H), 4.52 (s, 2H), 4.49 – 4.42 (m, 1H), 4.31 – 4.23 (m, 2H), 3.91 – 3.84 (m, 2H), 2.22 (s, 3H), 1.98 (s, 3H), 1.54 (d, $J$ = 7.1 Hz, 3H); $^{13}$C NMR (100 MHz, Methanol-$d_4$) δ 172.63, 146.32, 145.58, 144.99, 142.64, 138.93, 136.03, 132.62, 130.21, 127.84, 126.46, 125.63, 125.20, 124.25, 123.87, 122.97, 115.72, 112.54, 55.04, 50.47, 47.24, 39.15, 22.51, 22.39, 18.60; HRMS (ESI) calcd for $C_{26}H_{31}N_4O_2S$ [M+H]$^+$ 463.2162, found 463.2164.

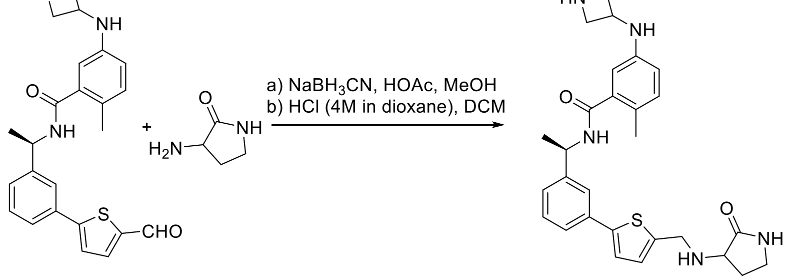


S4

XR8-77

**5-(azetidin-3-ylamino)-2-methyl-N-((1R)-1-(3-(5-(((2-oxopyrrolidin-3-yl)amino)methyl)thiophen-2-yl)phenyl)ethyl)benzamide (XR8-77)**. **S4** (31 mg, 0.06 mmol) and 3-aminopyrrolidin-2-one (9 mg, 0.09 mmol) was subjected to general reductive amination procedure with MeOH (2 mL), HOAc (500 µL) at 50 °C, and then $NaBH_3CN$ (12 mg, 0.18 mmol) was added. After purification by Prep-HPLC, the product was subjected to general N-Boc deprotection procedure with HCl (4M in dioxane, 100 µL) and DCM (2 mL). The purification by Prep-HPLC afforded the product **XR8-77** (19 mg, yield 63 % for 2 steps) as a white solid: $^1H$ NMR (400 MHz, Methanol-$d_4$) δ 8.28 (s, 2H), 7.71 – 7.65 (m, 1H), 7.56 – 7.51 (m, 1H), 7.42 – 7.30 (m, 3H), 7.14 (d, $J$ = 3.7 Hz, 1H), 7.03 (d, $J$ = 8.3 Hz, 1H), 6.59 (dd, $J$ = 8.2, 2.6 Hz, 1H), 6.55 – 6.51 (m, 1H), 5.21 (q, $J$ = 7.0 Hz, 1H), 4.49 (p, $J$ = 7.0 Hz, 1H), 4.39 – 4.27 (m, 4H), 3.98 – 3.90 (m, 2H), 3.79 – 3.72 (m, 1H), 3.44 – 3.33 (m, 2H), 2.55 – 2.46 (m, 1H), 2.22 (s, 3H), 2.11 – 1.99 (m, 1H), 1.55 (d, $J$ = 7.0 Hz, 3H); $^{13}C$ NMR (100 MHz, Methanol-$d_4$) δ 176.76, 172.60, 146.50, 145.47, 138.85, 135.69, 132.66, 130.45, 130.32, 126.75, 125.77, 125.73, 125.37, 124.44, 124.39, 115.82, 112.49, 57.49, 54.86, 50.49, 46.84, 46.14, 40.23, 28.33, 22.42, 18.61; HRMS (ESI) calcd for $C_{28}H_{34}N_5O_2S$ $[M+H]^+$ 504.2428, found 504.2431.

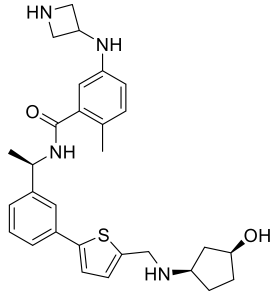


XR8-83

**5-(azetidin-3-ylamino)-N-((R)-1-(3-(5-((((1R,3S)-3-hydroxycyclopentyl)amino)methyl)thiophen-2-yl)phenyl)ethyl)-2-methylbenzamide (XR8-83)**. White solid (yield 67%): $[\alpha]^{25}_{546}$ = -6.8 (c 0.8, MeOH); $^1H$ NMR (400 MHz, Methanol-$d_4$) δ 8.40 (s, 1H), 7.73 – 7.67 (m, 1H), 7.57 – 7.52 (m, 1H), 7.40 (dd, $J$ = 6.8, 4.0 Hz, 3H), 7.28 (d, $J$ = 3.7 Hz, 1H), 7.03 (d, $J$ = 8.0 Hz, 1H), 6.61 – 6.55 (m, 2H), 5.21 (q, $J$ = 7.0 Hz, 1H), 4.55 – 4.43 (m, 3H), 4.39 – 4.30 (m, 3H), 3.97 – 3.89 (m, 2H), 3.68 (tt, $J$ = 8.0, 5.8 Hz, 1H), 2.30 – 2.12 (m, 5H), 2.05 – 1.93 (m, 1H), 1.89 – 1.82 (m, 3H), 1.55 (d, $J$ = 7.0 Hz, 3H); $^{13}C$ NMR (100 MHz, Methanol-$d_4$) δ 172.60, 148.34, 146.62, 145.49, 138.84, 135.15, 132.95, 132.85, 132.64, 130.46, 126.98, 125.71, 125.60, 124.92, 124.77, 115.56, 112.81, 72.52, 58.47, 54.89, 50.55, 46.85, 45.14, 39.26, 34.40, 28.32, 22.35, 18.62; HRMS (ESI) calcd for $C_{29}H_{37}N_4O_2S$ $[M+H]^+$ 505.2632, found 505.2634.

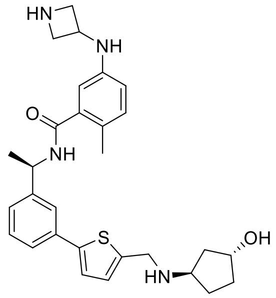


XR8-84

**5-(azetidin-3-ylamino)-N-((R)-1-(3-(5-((((1R,3R)-3-hydroxycyclopentyl)amino)methyl)thiophen-2-yl)phenyl)ethyl)-2-methylbenzamide (XR8-84)**. White solid (yield 65%): $[\alpha]^{25}_{546}$ = -3.2 (c 2.3, MeOH); $^1H$ NMR (400 MHz, Methanol-$d_4$) δ 8.42 (s, 1H), 7.69 (d, $J$ = 1.9 Hz, 1H), 7.57 – 7.52 (m, 1H), 7.43 – 7.36 (m, 3H), 7.28 (d, $J$ = 3.7 Hz,

1H), 7.03 (d, *J* = 8.0 Hz, 1H), 6.60 – 6.54 (m, 2H), 5.21 (q, *J* = 7.0 Hz, 1H), 4.55 – 4.31 (m, 6H), 3.98 – 3.90 (m, 2H), 3.89 – 3.80 (m, 1H), 2.39 – 2.27 (m, 1H), 2.23 – 2.13 (m, 4H), 2.11 – 2.02 (m, 1H), 1.92 – 1.83 (m, 1H), 1.80 – 1.67 (m, 2H), 1.55 (d, *J* = 7.0 Hz, 3H). $^{13}$C NMR (100 MHz, Methanol-$d_4$) δ 172.59, 148.27, 146.61, 145.49, 138.86, 135.17, 133.08, 132.80, 132.64, 130.45, 126.99, 125.73, 125.61, 124.91, 124.75, 115.57, 112.81, 72.45, 58.26, 54.90, 50.54, 46.86, 45.39, 39.88, 34.12, 28.21, 22.34, 18.62; HRMS (ESI) calcd for $C_{29}H_{37}N_4O_2S$ $[M+H]^+$ 505.2632, found 505.2636.

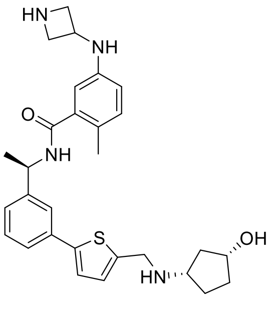


XR8-89

**5-(azetidin-3-ylamino)-N-((R)-1-(3-(5-((((1S,3R)-3-hydroxycyclopentyl)amino)methyl)thiophen-2-yl)phenyl)ethyl)-2-methylbenzamide (XR8-89)**. White solid (yield 72%): $[\alpha]^{25}_{546}$ = +12.9 (c 0.4, MeOH); $^1$H NMR (400 MHz, Methanol-$d_4$) δ 8.45 (s, 1H), 7.72 – 7.67 (m, 1H), 7.58 – 7.51 (m, 1H), 7.44 – 7.38 (m, 3H), 7.26 (d, *J* = 3.7 Hz, 1H), 7.03 (d, *J* = 8.1 Hz, 1H), 6.61 – 6.54 (m, 2H), 5.21 (q, *J* = 7.0 Hz, 1H), 4.50 (p, *J* = 7.0 Hz, 1H), 4.43 (s, 2H), 4.35 (dt, *J* = 8.8, 6.3 Hz, 3H), 3.93 (dd, *J* = 11.1, 6.6 Hz, 2H), 3.71 – 3.62 (m, 1H), 2.29 – 2.12 (m, 5H), 2.04 – 1.92 (m, 1H), 1.89 – 1.80 (m, 3H), 1.55 (d, *J* = 7.1 Hz, 3H); $^{13}$C NMR (100 MHz, Methanol-$d_4$) δ 172.60, 148.23, 146.61, 145.49, 138.89, 135.18, 133.26, 132.74, 132.64, 130.46, 126.92, 125.72, 125.60, 124.98, 124.76, 115.57, 112.79, 72.57, 58.51, 54.93, 50.55, 46.88, 45.24, 39.38, 34.43, 28.44, 22.33, 18.61; HRMS (ESI) calcd for $C_{29}H_{37}N_4O_2S$ $[M+H]^+$ 505.2632, found 505.2638.

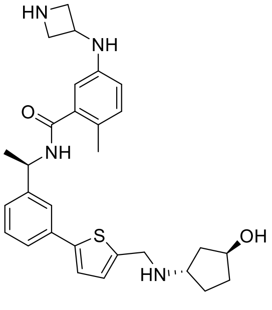


XR8-96

**5-(azetidin-3-ylamino)-N-((R)-1-(3-(5-((((1S,3S)-3-hydroxycyclopentyl)amino)methyl)thiophen-2-yl)phenyl)ethyl)-2-methylbenzamide (XR8-96)**. White solid (yield 72%): $[\alpha]^{25}_{546}$ = -3.2 (c 1.4, MeOH); $^1$H NMR (400 MHz, Methanol-$d_4$) δ 8.43 (s, 1H), 7.71 – 7.68 (m, 1H), 7.57 – 7.52 (m, 1H), 7.44 – 7.37 (m, 3H), 7.27 (d, *J* = 3.7 Hz, 1H), 7.03 (d, *J* = 8.0 Hz, 1H), 6.61 – 6.53 (m, 2H), 5.21 (q, *J* = 7.0 Hz, 1H), 4.51 (p, *J* = 7.0 Hz, 1H), 4.45 (s, 2H), 4.40 – 4.30 (m, 3H), 3.99 – 3.90 (m, 2H), 3.72 – 3.64 (m, 1H), 2.29 – 2.11 (m, 5H), 2.04 – 1.93 (m, 1H), 1.89 – 1.81 (m, 3H), 1.55 (d, *J* = 7.1 Hz, 3H); $^{13}$C NMR (100 MHz, Methanol-$d_4$) δ 172.59, 148.31, 146.62, 145.49, 138.86, 135.16, 132.98, 132.89, 132.64, 130.46, 126.97,

125.73, 125.60, 124.95, 124.77, 115.57, 112.82, 72.55, 58.51, 54.93, 50.55, 46.87, 45.19, 39.31, 34.41, 28.36, 22.34, 18.62; HRMS (ESI) calcd for $C_{29}H_{37}N_4O_2S$ $[M+H]^+$ 505.2632, found 505.2636.

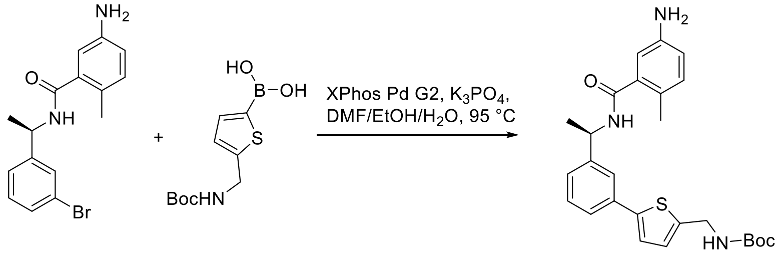


**XR8-98**

**(R)-3-((1-(3-(5-(((tert-butoxycarbonyl)amino)methyl)thiophen-2-yl)phenyl)ethyl)carbamoyl)-4-methylbenzenaminium (XR8-98)**. A flask fitted with a rubber septum was charged with (R)-5-amino-N-(1-(3-bromophenyl)ethyl)-2-methylbenzamide (33 mg, 0.10 mmol), (5-(((tert-butoxycarbonyl)amino)methyl)thiophen-2-yl)boronic acid (39 mg, 0.15 mmol), XPhos Pd G2 (8 mg, 0.01 mmol), $K_3PO_4$ (64 mg, 0.3 mmol), DMF/EtOH/$H_2O$ (1 mL/ 1 mL/ 0.5 mL) and then purged with argon. The mixture was stirred at 95 °C overnight. The reaction mixture was then cooled to room temperature, diluted with ethyl acetate (20 mL), filtered through celite and concentrated in vacuo. The purification by Prep-HPLC afforded the product **XR8-98** (41 mg, yield 88%) as a white solid: $[\alpha]^{25}_{546}$= +22.6 (c 1.5, MeOH); $^1H$ NMR (400 MHz, $CDCl_3$) δ 7.57 – 7.54 (m, 1H), 7.49 – 7.45 (m, 1H), 7.37 – 7.33 (m, 1H), 7.29 – 7.27 (m, 1H), 7.13 (d, *J* = 3.6 Hz, 1H), 6.97 (d, *J* = 8.1 Hz, 1H), 6.90 (d, *J* = 3.6 Hz, 1H), 6.70 (d, *J* = 2.5 Hz, 1H), 6.63 (dd, *J* = 8.1, 2.6 Hz, 1H), 6.00 (d, *J* = 8.0 Hz, 1H), 5.36 – 5.27 (m, 1H), 4.46 (d, *J* = 5.9 Hz, 2H), 2.30 (s, 3H), 1.59 (d, *J* = 6.9 Hz, 3H), 1.47 (s, 9H); $^{13}C$ NMR (100 MHz, $CDCl_3$) δ 169.44, 144.35, 144.05, 143.80, 141.83, 137.19, 134.94, 132.00, 129.43, 126.58, 125.39, 124.94, 123.72, 123.06, 116.83, 113.57, 49.03, 39.95, 28.54, 22.01, 18.90; HRMS (ESI) calcd for $C_{26}H_{32}N_3O_3S$ $[M+H]^+$ 466.2159, found 466.2157.

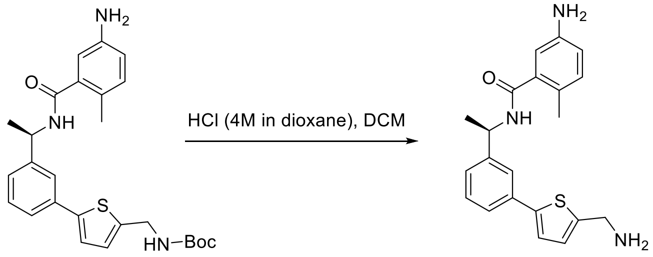


**XR8-98** **XR8-101**

**(R)-5-amino-N-(1-(3-(5-(aminomethyl)thiophen-2-yl)phenyl)ethyl)-2-methylbenzamide (XR8-101)**. Compound **XR8-98** (20 mg, 0.04 mmol) was subjected to general N-Boc deprotection procedure with HCl (4M in dioxane, 100 µL) and DCM (2 mL). The purification by Prep-HPLC afforded the product **XR8-101** (14 mg, yield 89%) as a white solid: $[\alpha]^{25}_{546}$= -2.4 (c 2.0, MeOH); $^1H$ NMR (400 MHz, Methanol-$d_4$) δ 7.71 (q, *J* = 1.5 Hz, 1H), 7.59 – 7.53 (m, 1H), 7.45 – 7.35 (m, 6H), 7.26 – 7.22 (m, 1H), 5.25 (q, *J* = 7.0 Hz, 1H), 4.36 (s, 2H), 2.37 (s, 3H), 1.59 (d, *J* = 7.1 Hz, 3H); $^{13}C$ NMR (100 MHz, Methanol-$d_4$) δ 170.32, 147.64, 146.27, 139.89, 137.97, 135.35, 134.91, 133.53, 131.71, 130.52, 129.57, 126.86, 125.71, 125.18, 124.94, 124.73, 122.66, 50.78, 38.85, 22.27, 19.19; HRMS (ESI) calcd for $C_{21}H_{23}N_3OS$ $[M+H]^+$ 366.1635, found 366.1635

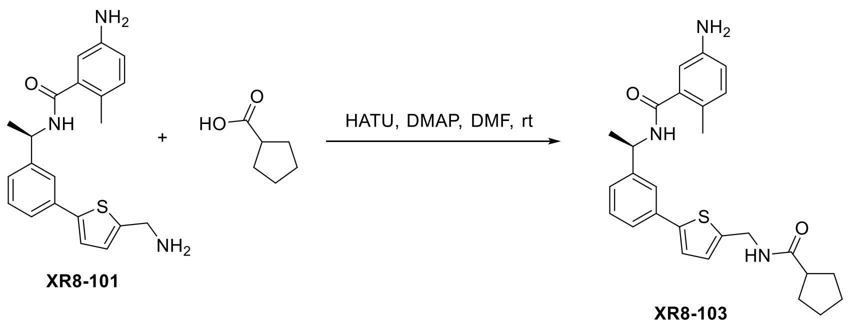


**(R)-5-amino-N-(1-(3-(5-(cyclopentanecarboxamidomethyl)thiophen-2-yl)phenyl)ethyl)-2-methylbenzamide (XR8-103)**. **XR8-101** (20 mg, 0.05 mmol), cyclopentanecarboxylic acid (6 mg, 0.05 mmol), HATU (19 mg, 0.05 mmol), and DMAP (18 mg, 0.15 mmol) was subjected to general amine coupling procedure with DMF (2 mL). The purification by Prep-HPLC gave the product **XR8-103** (20 mg, yield 87%) as a white solid: $[\alpha]^{25}_{546}$ = +23.5 (c 0.8, MeOH); $^{1}$H NMR (400 MHz, Methanol-$d_4$) δ 7.66 – 7.61 (m, 1H), 7.50 – 7.45 (m, 1H), 7.38 – 7.28 (m, 2H), 7.22 (d, *J* = 3.6 Hz, 1H), 6.98 – 6.91 (m, 2H), 6.74 – 6.66 (m, 2H), 5.24 – 5.14 (m, 1H), 4.53 – 4.49 (m, 2H), 2.65 (ddd, *J* = 13.3, 10.5, 7.5 Hz, 1H), 2.20 (s, 3H), 1.91 – 1.82 (m, 2H), 1.80 – 1.69 (m, 4H), 1.65 – 1.56 (m, 2H), 1.53 (d, *J* = 7.1 Hz, 3H); $^{13}$C NMR (100 MHz, Methanol-$d_4$) δ 178.96, 172.83, 146.46, 146.32, 144.88, 143.02, 138.53, 136.03, 132.29, 130.17, 127.60, 126.34, 125.44, 125.17, 124.28, 123.80, 118.01, 115.04, 50.39, 46.46, 39.13, 31.43, 27.02, 22.42, 18.67; HRMS (ESI) calcd for $C_{27}H_{32}N_3O_2S$ $[M+H]^+$ 462.2210, found 462.2214.

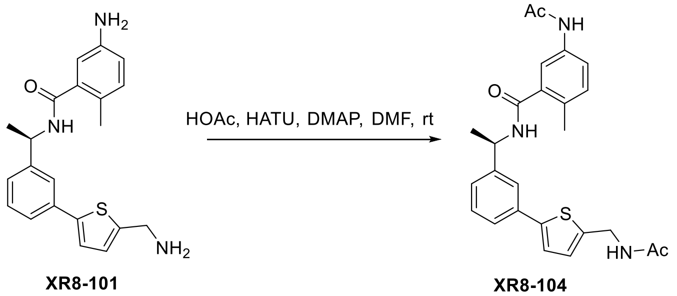


**(R)-5-acetamido-N-(1-(3-(5-(acetamidomethyl)thiophen-2-yl)phenyl)ethyl)-2-methylbenzamide (XR8-104)**. **XR8-101** (20 mg, 0.05 mmol), HOAc (6 mg, 0.10 mmol), HATU (19 mg, 0.05 mmol), and DMAP (18 mg, 0.15 mmol) was subjected to general amine coupling procedure with DMF (2 mL). The purification by Prep-HPLC gave the compound **XR8-104** (17 mg, yield 76%) as a white solid: $^{1}$H NMR (400 MHz, Methanol-$d_4$) δ 7.66 – 7.63 (m, 1H), 7.58 (d, *J* = 2.3 Hz, 1H), 7.50 – 7.43 (m, 2H), 7.38 – 7.30 (m, 2H), 7.23 (d, *J* = 3.6 Hz, 1H), 7.17 (d, *J* = 8.3 Hz, 1H), 6.96 – 6.92 (m, 1H), 5.21 (q, *J* = 7.0 Hz, 1H), 4.51 (d, *J* = 0.9 Hz, 2H), 2.29 (s, 3H), 2.10 (s, 3H), 1.97 (s, 3H), 1.55 (d, *J* = 7.1 Hz, 3H); $^{13}$C NMR (100 MHz, Methanol-$d_4$) δ 172.93, 171.89, 171.72, 146.21, 144.94, 142.48, 138.31, 137.56, 136.02, 132.27, 132.05, 130.24, 127.87, 126.32, 125.25, 124.32, 123.91, 122.43, 119.89, 50.50, 39.14, 23.72, 22.50, 22.40, 19.06; HRMS (ESI) calcd for $C_{25}H_{28}N_3O_3S$ $[M+H]^+$ 450.1846, found 450.1852.

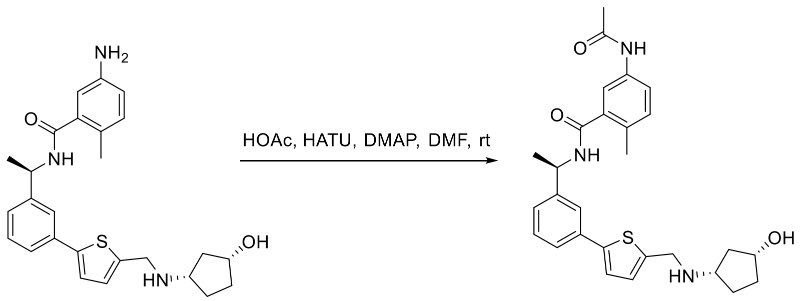


XR8-106

**5-acetamido-N-((R)-1-(3-(5-((((1S,3R)-3-hydroxycyclopentyl)amino)methyl)thiophen-2-yl)phenyl)ethyl)-2-methylbenzamide (XR8-106)**. 5-amino-N-((R)-1-(3-(5-((((1S,3R)-3-hydroxycyclopentyl)amino)methyl)thiophen-2-yl)phenyl)ethyl)-2-methylbenzamide (23 mg, 0.05 mmol), HOAc (3 mg, 0.05 mmol), HATU (19 mg, 0.05 mmol), and DMAP (18 mg, 0.15 mmol) was subjected to general amine coupling procedure with DMF (2 mL). The purification by Prep-HPLC gave the compound **XR8-106** (17 mg, yield 76%) as a white solid: $[\alpha]^{25}_{546}$ = -18.7 (c 0.5, MeOH); $^1$H NMR (400 MHz, Methanol-$d_4$) δ 8.36 (s, 1H), 7.73 – 7.66 (m, 2H), 7.58 – 7.53 (m, 1H), 7.44 – 7.36 (m, 4H), 7.27 (d, *J* = 3.7 Hz, 1H), 7.18 (d, *J* = 8.2 Hz, 1H), 5.22 (q, *J* = 7.0 Hz, 1H), 4.45 (s, 2H), 4.33 (p, *J* = 4.0 Hz, 1H), 3.68 (p, *J* = 7.0 Hz, 1H), 2.30 (s, 3H), 2.27 – 2.14 (m, 2H), 2.12 (s, 3H), 2.03 – 1.92 (m, 1H), 1.89 – 1.80 (m, 3H), 1.56 (d, *J* = 7.0 Hz, 3H); $^{13}$C NMR (100 MHz, Methanol-$d_4$) δ 171.96, 171.77, 148.41, 146.57, 138.25, 137.60, 135.14, 132.97, 132.74, 132.29, 132.08, 130.49, 126.96, 125.56, 124.83, 124.80, 122.41, 119.91, 72.52, 58.47, 50.53, 45.17, 39.27, 34.42, 28.32, 23.74, 22.35, 19.05; HRMS (ESI) calcd for $C_{28}H_{34}N_3O_3S$ $[M+H]^+$ 492.2315, found 492.2319.

**Abbreviations list**

| **Abbreviation** | **Name** |
|---|---|
| aq. | Aqueous solution |
| Ar | Argon |
| Boc | tert-Butyloxycarbonyl |
| DCM | Dichloromethane |
| DMAP | 4,4-Dimethylaminopyridine |
| DMF | Dimethylformamide |
| DIPEA | N,N-Diisopropylethylamine |
| EtOAc or EA | Ethyl acetate |
| equiv. or eq. | Equivalent |

| | |
|---|---|
| Et | Ethyl |
| ESI | Electron spray ionization |
| HATU | Hexafluorophosphate azabenzotriazole tetramethyl uronium |
| HOAc | Acetic acid |
| HRMS | High-resolution mass spectrometry |
| LRMS | Low-resolution mass spectrometry |
| m/z | Ratio of mass to charge |
| Me | Methyl |
| MeCN | Acetonitrile |
| NMR | Nuclear magnetic resonance |
| rt | Room temperature |
| TEA | Triethanolamine |
